# Horizontal transfer promotes allele segregation in multicopy plasmids

**DOI:** 10.64898/2026.03.13.711586

**Authors:** Lisa M. Hartmann, Mario Santer, Nils F. Hülter, Tal Dagan

## Abstract

Plasmids reside within prokaryotic cells in multiple copies and can spread horizontally between hosts. Their multicopy nature enables intracellular allele diversity (heteroplasmy), and their segregation depends on the modes of plasmid replication and partition. Horizontal plasmid transfer has the potential to alter plasmid allele composition, however its impact on plasmid allele dynamics remains poorly understood. Here, we show that conjugative plasmid transfer accelerates the segregation of plasmid heteroplasmy by promoting the emergence of homoplasmic hosts. Using a quantitative experimental system to track plasmid allele dynamics in *Acinetobacter baylyi* under non-selective conditions, we followed the fate of a novel antibiotic resistance allele introduced into an ancestral donor population. While alleles in heteroplasmic donors segregated over time, conjugation produced almost only homoplasmic recipients whose allele composition closely mirrored that of the donor pool. Heteroplasmic recipients were rare and arose primarily from multiple plasmid transfer events prior to plasmid establishment. A mathematical model calibrated to the experimental setup predicts that conjugation accelerates plasmid allele segregation, with effects scaling with plasmid copy number and conjugation frequency. Our findings identify horizontal transfer as a previously unrecognized segregation pathway shaping the evolution of mobile genetic elements.

## Introduction

Plasmids are extrachromosomal genetic elements that play a central role in microbial evolution due to their contribution to horizontal DNA transfer. Mobilizable and self-transmissible (conjugative) plasmids have been implicated as key drivers in the spread of antibiotic resistance genes^1,2^. It has been estimated that 30% of the plasmids in bacteria isolated from humans carry antibiotic resistance genes^1^. Plasmids are frequently maintained in multiple copies per cell, hence the emergence of plasmid mutations generates intracellular genetic diversity, termed heteroplasmy (or plasmid heterozygosity). Multicopy plasmids have a higher mutational supply compared to chromosomes, which are monoploid in many human pathogens (e.g., ESKAPE pathogens). Indeed, the evolution of antibiotic resistance genes is accelerated in high-copy, non-mobile plasmids relative to chromosomal genes^3^. However, the evolutionary rate of multicopy plasmids is also constrained by segregational drift – genetic drift acting on the intracellular plasmid pool – that lowers the establishment probability of novel mutations^4,5^. Random segregation of plasmid copies at cell division can lead to rapid loss of novel mutations (i.e., low-frequency alleles) soon after their emergence^6^. For example, antibiotic resistance genes encoded in multicopy plasmids may be rapidly lost due to segregational drift, also under selection for antibiotic resistance^7^. The plasmid copy number is a major determinant of plasmid allele segregation, affecting the dynamics of both neutral and slightly beneficial alleles^7,8^. Additional determinants are the modes of plasmid replication and segregation, that influence plasmid allele dynamics via their effect on the intracellular plasmid allele composition^6,9,10^.

Studies on the factors influencing the fate of plasmid alleles have so far focussed primarily on vertically inherited, non-mobile plasmids. However, the role of conjugation in plasmid allele dynamics remains poorly understood. Conjugative plasmids are typically maintained in low numbers ranging between two and four copies per cell^11,12^, a range that permits plasmid heteroplasmy and renders plasmid alleles susceptible to segregational drift^7,8^. Conjugation involves the transfer of plasmid DNA from a donor to a recipient via a type IV secretion system, typically mediated by a pilus^13^. During conjugation, a single-stranded copy of the plasmid is transferred and subsequently established in the recipient as double-stranded DNA^14^. Redundant plasmid transfers into the same recipient cell are generally inhibited by entry and surface exclusion mechanisms that prevent lethal zygosis, a phenomenon in which excess ssDNA leads to replication defects and cell death^15–17^. Thus transfer of a single plasmid variant from a heteroplamic donor is expected to generate a homoplasmic recipient. Nonetheless, a delay in the expression of surface exclusion mechanisms after initial plasmid entry may allow multiple plasmid invasions into the same recipient cell^18^, potentially giving rise to heteroplasmic recipients. In the longer term, the composition of plasmid alleles in the recipient population may further depend on fitness effects of the transferred plasmid^19^, particularly if different plasmid variants impose different fitness effects on their host. Notably, under strong selection for plasmid encoded traits, conjugation has no effect on plasmid allele dynamics since plasmid-free cells are removed from the population, hence plasmid alleles dynamics under strong selective regime are affected by vertical inheritance only^20,21^.

Here we establish an experimental system to quantify the effect of conjugation on plasmid allele dynamics. Using an experimental evolution setup, we examine the fate of a novel antibiotic resistance allele in three model conjugative plasmids that vary in their host range or copy number. The results of the evolution experiment are further used to parametrize a mathematical model that enables us to compare the dynamics of plasmid alleles between non-mobile and conjugative plasmids. Our results supply an empirical framework for conjugative plasmid allele dynamics under non-selective conditions, while disentangling the effects of segregational drift and DNA transfer. We show evidence for persistent heteroplasmy for two model conjugative plasmids. Furthermore, we show that conjugation generates mostly homoplasmic cells and thus accelerates plasmid allele segregation. Complementary modelling of timescales beyond the temporal scope of the evolution experiment predicts that the fate of novel plasmid alleles with and without conjugation ultimately converge.

## Results

### An experimental system for following allele dynamics in low-copy conjugative plasmids

To quantify the effect of conjugation on plasmid allele dynamics, we established an experimental system to follow the dynamics of an antibiotic resistance allele in self-transmissible plasmids. Model plasmids were evolved in a ‘donor’ population, where the novel allele is introduced at the onset of the evolution experiment. Daily samples from the donor population were mated with a ‘recipient’ population. Plasmid allele composition in the populations of donors and plasmid recipients was evaluated and compared (Fig. 1a). We constructed three model plasmids, into each of which we inserted a locus containing *gfp*, which codes for green fluorescent protein. The *gfp* locus is flanked by the 5’ and 3’ terminal sections of the *nptII* gene, which encodes neomycin phosphotransferase II, conferring resistance to kanamycin. The focal novel allele (*nptII*) was introduced into the ancestral donor population by natural transformation. Recombination between the taken-up novel allele and the plasmid target locus occurs on only a single plasmid copy within the intracellular plasmid pool due to one-hit kinetics^22^. Consequently, only the transformed plasmid copy replaces the ancestral *gfp* allele with the novel *nptII* allele^38^. The initial donor population thus comprises either homoplasmic cells hosting only the ancestral allele or heteroplasmic cells harboring two plasmid variants carrying either the ancestral or novel allele. Plasmid segregation during heteroplasmic cell growth is expected to yield homoplasmic cells for the novel or ancestral alleles.

**Figure 1.**
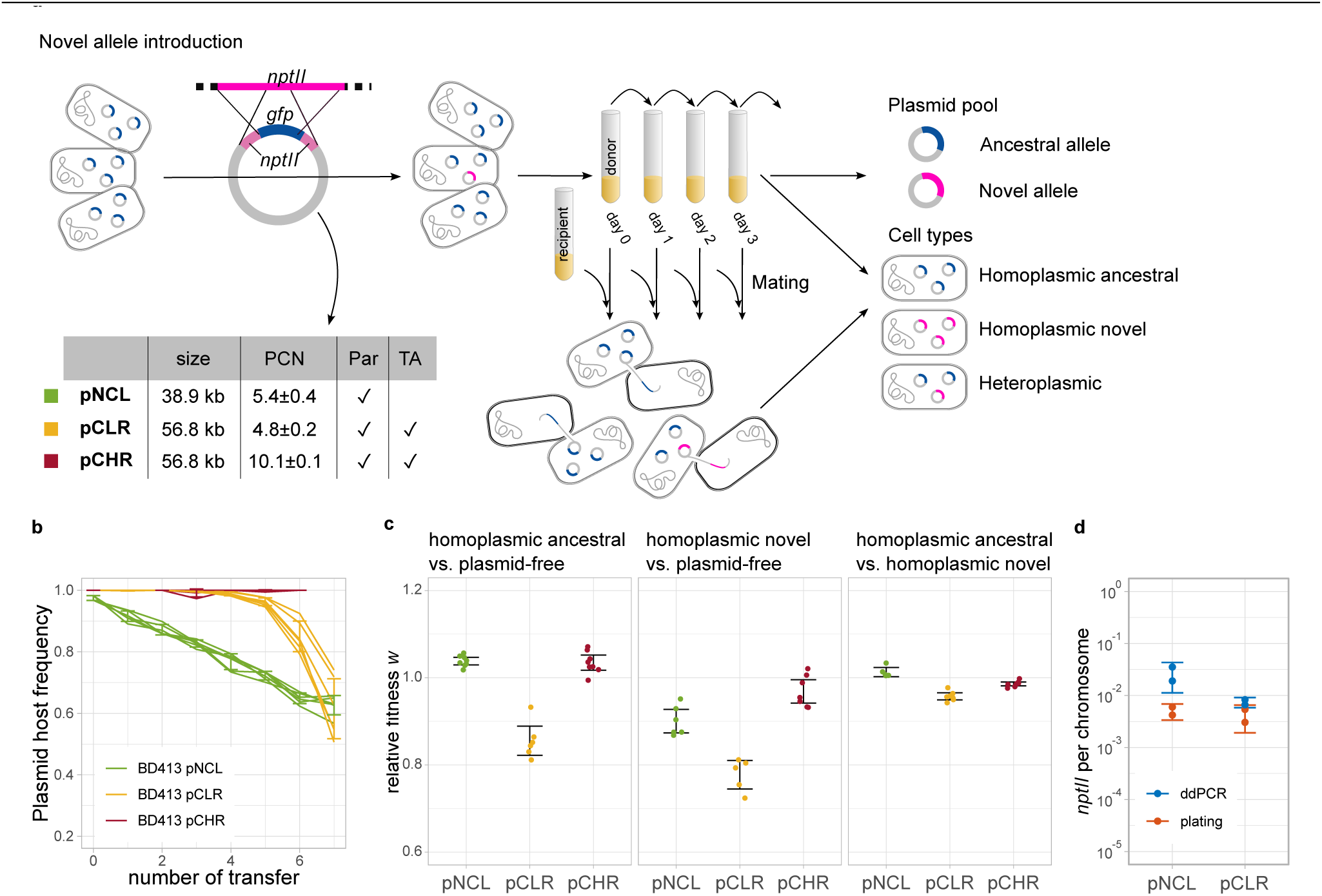
An experimental system to follow the dynamics of conjugative plasmid alleles. Description of the evolution experiment workflow and model plasmid properties. (a) The novel kanamycin resistance allele (*nptII*) was introduced into a homoplasmic population of plasmid hosts where it integrates into a single plasmid copy in a subset of cells. The donor population was evolved by daily serial transfers; agitation prevents conjugation within the donor culture (Table S5). Cell type frequencies are quantified by ddPCR. Donors are mated daily with a plasmid-free recipient population on solid agar. Recipient genotypes are determined by differential plating. This design allows separate observation of conjugation and segregational drift. Plasmids copy number is supplied in the table below as the mean ±95% CI. The presence of partitioning system (Par) and toxin-antitoxin system (TA) is indicated. (b) Model plasmid stability in the *Acinetobacter baylyi* BD413 host over 7 days (transfers). Error bars correspond to standard deviation. (c) Relative fitness *w* of homoplasmic ancestral hosts compared with plasmid-free hosts (with the following sample sizes: n_pNCL_=8; n_pCLR_=6; n_pCHR_=8) (left), homoplasmic novel allele host compared to plasmid-free cells (n_pNCL_=6; n_pCLR_=5; n_pCHR_=7) (middle), and homoplasmic ancestral allele hosts compared with homoplasmic novel allele hosts (n_pNCL_=7; n_pCLR_=5; n_pCHR_=8) (right). Error bars correspond to 95% CI; time series data of the measurements, linear regression analysis and relative fitness values are supplied in Fig. S1. d) Allele frequency as quantified by the established ddPCR protocol was validated by plating. Error bars correspond to 95% CI.

The model plasmid pNCL (<u>N</u>arrow-host-range, <u>C</u>onjugative, <u>L</u>ow-copy) was constructed from the conjugative low-GC plasmid pHHV216 that was captured from soil samples. The plasmid pHHV216 confers resistance to chloramphenicol, streptomycin, sulfonamide and tetracycline and is considered to have a narrow host range in *Acinetobacter*^23^. Model plasmid pCLR (<u>C</u>onjugative, <u>L</u>ow-copy <u>R</u>K2) was constructed from the conjugative IncPα broad-host range plasmid^24,25^. Model plasmid pCHR (<u>C</u>onjugative, <u>H</u>igh-copy <u>R</u>K2) was constructed from pCLR by inserting a point mutation into *trfA* (Arg → Cys at 271 aa position), which controls the replication initiation frequency in RK2^26^. The model plasmid sizes range between 38 Kb and 56 Kb. The three plasmids harbor an active partition system and the RK2 derivatives further harbor a toxin-antitoxin system^27^ (Fig. 1a). The model plasmids were introduced into *Acinetobacter baylyi* BD413 as the host. The plasmid copy number (PCN) in *A. baylyi* was ca. five copies for pNCL and pCLR and ca. ten copies for pCHR (Fig. 1a). The plasmids pNCL and pCLR are unstable in *A. baylyi*, while pCHR is stably maintained over time (Fig. 1b). The fitness of pNCL and pCHR homoplasmic ancestral hosts was slightly higher relative to plasmid-free cells, while pCLR has an intermediate negative fitness effect on the host (Fig. 1c). The fitness of novel homoplasmic hosts was lower relative to plasmid-free cells for all three plasmids (Fig. 1c). Additionally, slight fitness differences between homoplasmic hosts of the ancestral and novel alleles were observed for all plasmids. Homoplasmic novel pNCL hosts have a slightly lower fitness compared to the homoplasmic hosts of the ancestral allele (Fig. 1c). In contrast, homoplasmic novel hosts of pCLR and pCHR have slightly higher fitness compared to the homoplasmic hosts of the ancestral allele (Fig. 1c).

The plasmid allele composition is associated with distinct phenotypic traits: homoplasmic hosts are either fluorescent (ancestral allele carriers) or resistant to kanamycin (novel allele carriers) and heteroplasmic hosts have both traits. To document the host homo- and hetero-plasmy states we established a digital droplet PCR (ddPCR) assay. This procedure further enabled us to document allele frequency within the total population plasmid pool. Using the ddPCR assay, we detected cells carrying the novel allele at a 2- to 5-fold higher frequency than with phenotypic tests, depending on the model plasmid (Fig. 1d). Further validation of the ddPCR approach for the detection of plasmid heteroplasmy was performed with populations in the evolution experiment (Fig. S4). In the following we report the donor cell genotypes as observed using ddPCR measurements, unless stated otherwise.

### The novel allele is stably maintained in the donor population

The evolution experiment was performed with donor populations harboring one of the three model plasmids and propagated daily by serial transfers with a bottleneck of 1:100 (∼1×10^7^ cells) across six replicates in liquid media. The evolution experiment spanned 8-13 transfers (depending on the model plasmid) corresponding to 53-87 generations. Plasmid transfer within the donor populations was exceedingly rare under these culturing conditions (Table S5). At the onset of the evolution experiments, the novel plasmid allele was introduced at a frequency of ca. 5×10^-3^ heteroplasmic donors in the pNCL and pCLR populations. The pCHR ancestral population contained ca. 3×10^-3^ heteroplasmic donors (Table S6).

At the first transfer, 24 h after the novel allele introduction, the majority of pNCL novel allele carriers were heteroplasmic; pNCL hosts that were homoplasmic for the novel allele were observed at a frequency of ca. 5×10^-2^. The ddPCR measurements of host cell types in the pNCL donor population for the first evolution experiment transfer were not significantly different from the phenotypic observations (Fig. S4). At the second transfer of the pNCL donor populations, heteroplasmic and homoplasmic novel cells reached similar frequencies. Afterwards, the frequency of heteroplasmic cells declined rapidly and stabilized after the 7^th^ transfer with a low frequency of ca. 7×10^-3^ (Fig. 2a). At the same time, the frequency of homoplasmic novel pNCL hosts increased to a frequency of ca. 6×10^-2^ and remained stable until the end of the evolution experiment (Fig. 2a). The pNCL novel allele frequency was stable in the donor population throughout the evolution experiment (Fig. 2b).

**Figure 2.**
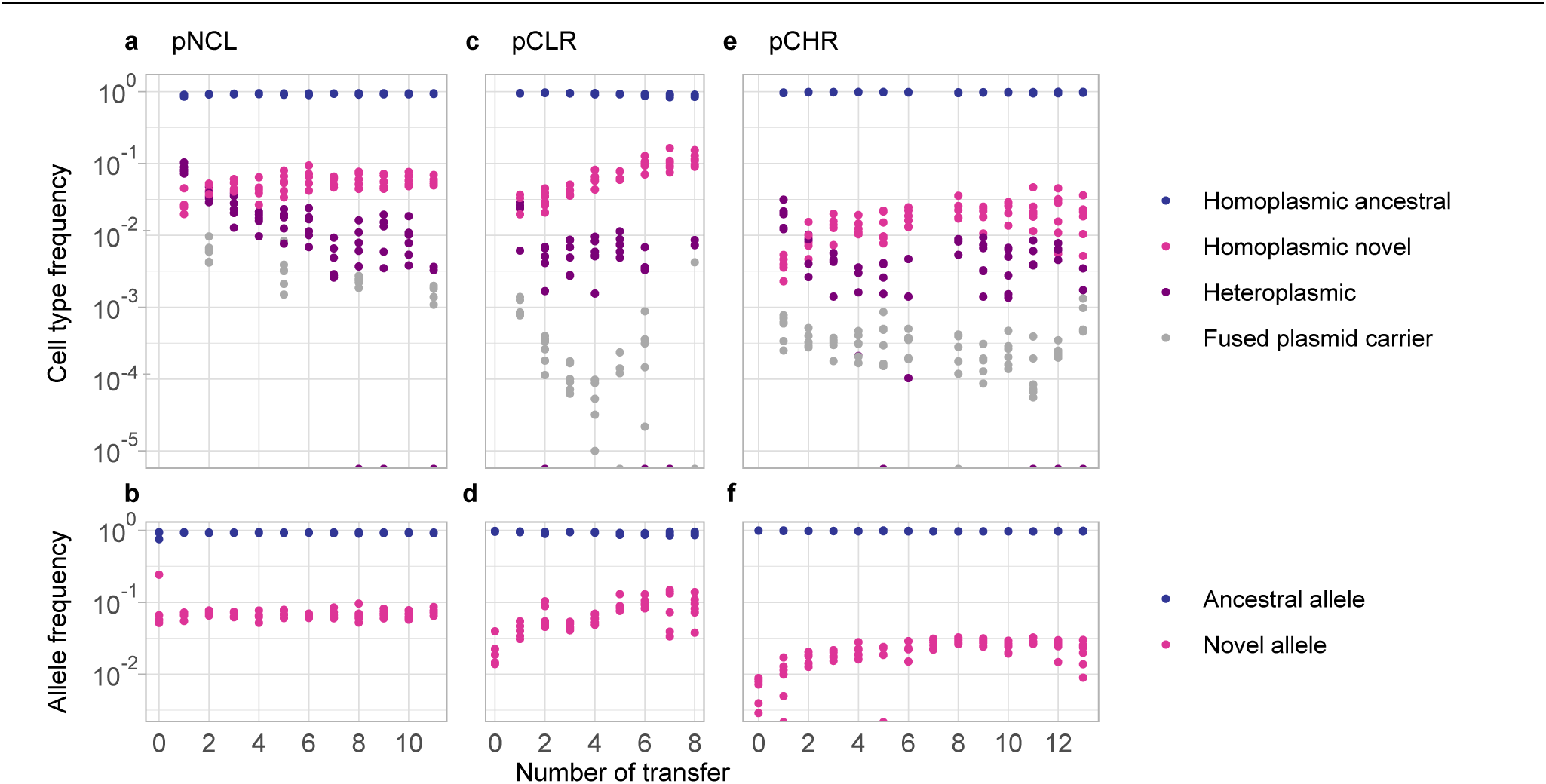
Allele segregation and maintenance of plasmid heteroplasmy in the donor populations over time. (a, c, e) Donor cell type frequencies relative to the total plasmid hosts over time in the model plasmid experimental evolution. The presence of segregating heteroplasmic cells was validated (Fig. S6a-b) and plasmid composition in heteroplasmic hosts was examined for plasmid fusion (Fig. S7a-d). The frequency of fused plasmid carriers was lower compared to segregating heteroplasmic cells. Heteroplasmic cells persisted in the pCLR cultures for an extended period while allele frequency in the population remained stable (Fig S8a-b). (b, d, f) Allele frequency in the donor plasmid pools over the evolution experiment duration as documented by ddPCR.

The plasmid allele dynamics observed for pCLR and pCHR were similar to those of pNCL, with several differences. Heteroplasmic and homoplasmic novel pCLR hosts reached similar frequencies already after 24 h (Fig. 2c). The frequency of heteroplasmic pCLR hosts stabilized after three transfers, while the frequency of homoplasmic novel allele hosts continued to gradually increase afterwards. After six transfers, the proportion of heteroplasmic pCLR hosts further decreased below the detection limit in most replicates. Note that after six transfers, the proportion of pCLR hosts decreased in most populations below 50% (Fig. S5). Unlike pNCL, the novel allele frequency in pCLR and pCHR increased over the first six transfers (Fig. 2d, f). The novel allele segregation dynamics in pCHR donor cells were similar to the dynamics in pNCL, despite the lower initial proportion of heteroplasmic donors (Fig. 2e). The decline of heteroplasmic pCHR host frequency mirrored that of pNCL donors at the onset of the experiment. The overall novel allele frequency in the pCHR plasmid pool over the evolution experiment showed a positive trend, similar to that of pCLR. At the end of the experiment, the novel allele frequency in pCHR was similar to the frequency of homoplasmic novel pCHR host (Fig. 2f). The increase in the novel allele frequency in pCLR and pCHR may be related to higher fitness of pCLR and pCHR homoplasmic novel hosts compared to the respective homoplasmic ancestral hosts (Fig. 1c).

Towards the end of the experiments, the novel allele frequency was similar to the respective homoplasmic novel host frequencies for all three model plasmids. Heteroplasmic cells had a negligible effect on the novel allele frequency, due to their low frequency in the donor population. Taken together, allele segregation changed the intracellular plasmid composition, but not the relative ratio of the ancestral and novel plasmid alleles in the population. Furthermore, plasmid backbone properties, including the plasmid copy number, appear to play a role in the persistent heteroplasmy among conjugative plasmids.

### Maintenance of plasmids heteroplasmy in the donor population exceeds the theoretical expectation

The evolution experiment of all three plasmids revealed that heteroplasmy was maintained for nearly the entire duration of the experiment. For instance, in pNCL, heteroplasmic cells declined by about one order of magnitude between transfer 4 and 8, then stabilized in most replicates. Under random segregation of plasmid copies, we expected faster exponential decay of heteroplasmy with an approximate loss of 20% per generation corresponding to 77% per transfer^9^ (see supplemental text S1 for the mathematical model). Consequently, we considered possible mechanisms for deviation from the theoretical expectation for the maintenance of heteroplasmy. These include non-random plasmid segregation, e.g., due to plasmid fusion or active partition mechanisms as well as the mode of plasmid replication. Fusion of plasmids carrying the novel and ancestral alleles into heteromultimers may lead to detection of heteroplasmic cells that cannot segregate. Testing for the presence of fused plasmid in the donor populations indeed revealed the presence of heteromultimeric plasmids carrying both alleles. The frequency of fused plasmid carriers in the pNCL donor population was ca. 5-fold lower than the frequency of heteroplasmic cells (Fig. 2a) and ca. 10-fold lower in the pCLR and pCHR donor populations (Fig. 2c and 2e). Hence, the frequency of cells carrying fused plasmids does not fully explain the frequency of heteroplasmic cells in the donor populations. According to our model, frequent fission of plasmid heteromultimers can prolong the maintenance of heteroplasmy, but not reproduce the plateau-like dynamics observed in donor cultures (Fig. S9c).

Active partition mechanism, found in the three model plasmids, may promote the partition of isogenic sister plasmids into both daughter cells and, consequently, the maintenance of heteroplasmy (Fig. S9j-k). To test for contribution of an active partition mechanism to pCLR segregation dynamics we generated a pCLR-Δ*incC* variant where active partition is deregulated^28–30^. Evolving that plasmid variant revealed the presence of heteroplasmic hosts throughout the experiment in the absence of fused plasmids (Fig. S10a-c). Consequently, we conclude that the pCLR-encoded active partition mechanism is unlikely contributing to the maintenance of pCLR heteroplasmy. Finally, to test for an effect of the plasmid origin of replication we generated pNCL::*ori*_pTAD_ by replacing the pNCL origin of replication, including the predicted replication initiation protein (*rep)* with that of pTAD (Fig S10d), which is considered to undergo regular replication^7^. This pNCL derivative had a plasmid copy number of five copies per cell. Conducting the evolution experiment with pNCL::*ori*_pTAD_ showed a rapid decline in the proportion of heteroplasmic cells in some replicates, while heteroplasmic cells were maintained in other replicates in lower proportion compared to pNCL (Fig. S.10e-f). Both scenarios matched our mathematical model of random segregation for the first four to six transfers (Fig. S9b). These results suggest an effect of the pNCL origin of replication on the observed pNCL allele dynamics.

### The novel allele frequency in recipient cells resembles that of the donor plasmid pool

To quantify the contribution of conjugation to plasmid allele dynamics, aliquots of evolving donor populations from successive days of the serial evolution experiment were mated with recipient populations at equal densities (1:1) in six replicates (Fig. 1a). Mating durations were minimized to below five cell divisions in order to limit the influence of plasmid recipient growth on plasmid genotype frequencies, e.g., due to growth deficit after plasmid acquisition. The frequencies of hosts carrying either the novel or ancestral allele, in homo- and heteroplasmic states, were determined within the plasmid-recipients (exconjugants) by plating. Note that conjugation frequencies of plasmid variants carrying the novel and ancestral alleles are not significantly different (Fig. S12a).

Plasmid hosts in the pNCL recipient population consisted predominantly of cells carrying the ancestral allele. Hosts harboring the novel allele were detected at low frequencies, ranging between 3×10^-2^ and 5×10^-2^ throughout the experiment (Fig. 3a). Heteroplasmic hosts carrying both alleles were rarely observed in the recipient population. Further analysis revealed that these colonies harbored fused plasmids that were detected at a low abundance from day 3 until the end of the experiment (Fig. 3a; Fig. S11a-d). The frequency of homoplasmic novel hosts in the pCLR recipient population was ca. 1×10^-1^ and stabilized after three days (Fig 3b). The novel allele frequency among pCHR recipients was initially low and increased steadily by ca. 3-fold (Fig. 3c). No evidence for transfer of fused plasmids was detected in pCLR and pCHR recipients, consistent with the low frequency of fused plasmids in the donor population (Fig. 2c, 2e).

**Figure 3:**
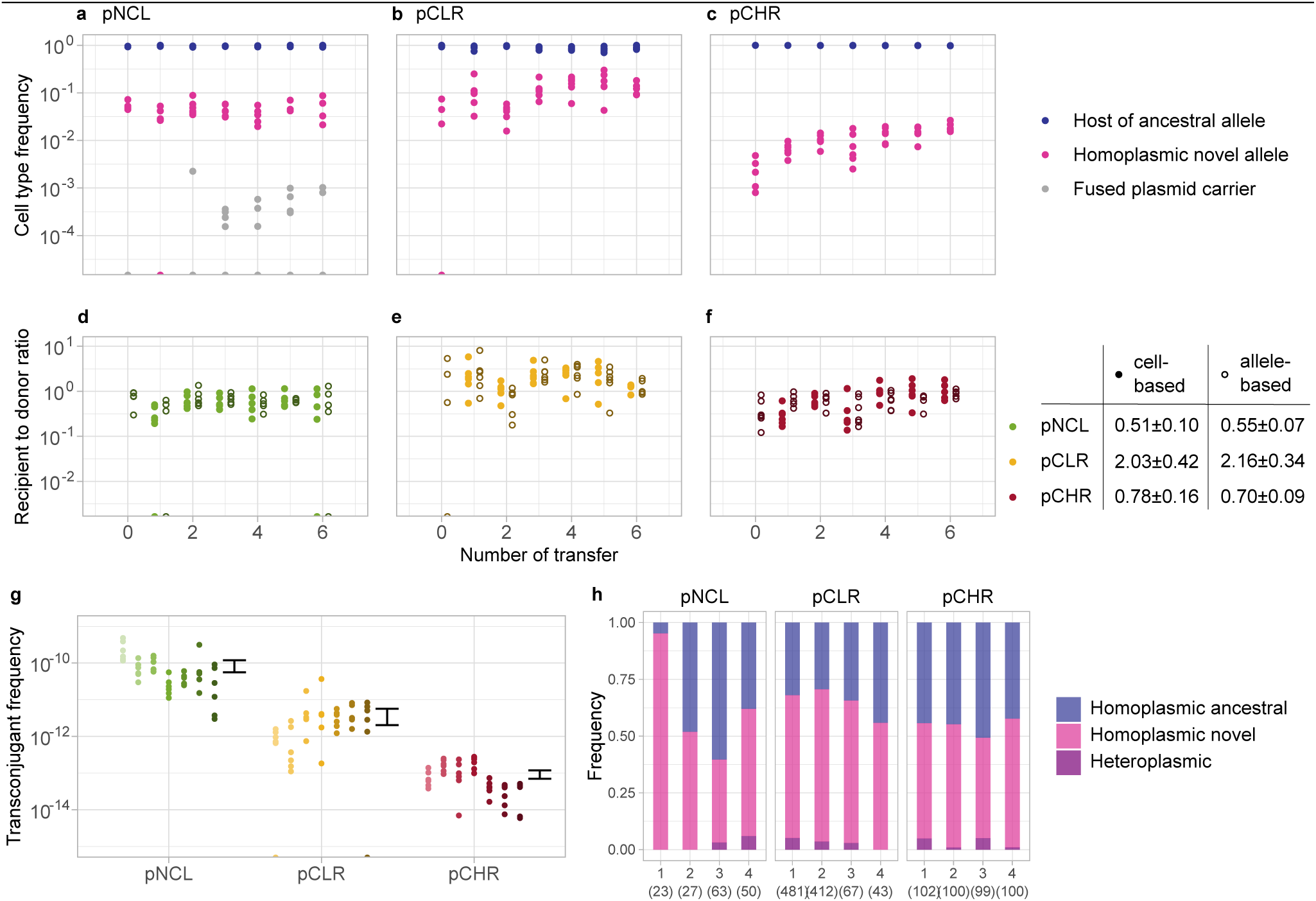
The donor allele composition is reflected in the recipient population. (a, b, c) Cell type frequency in recipient populations over the evolution experiment duration. The presence of heteroplasmic recipients carrying fused plasmids was validated in pNCL recipient populations (Fig S11 a-d). (d, e, f) Comparison of novel allele frequency to donor populations using the cell- and allele-based recipients to donors ratio. The table on the left shows the mean ± 95% CI ratio. (g) Conjugation frequency over the evolution experiment, ordered by day, was significantly different among the model plasmids. The highest conjugation frequency was observed for pNCL followed by pCLR. Over the experiment duration, the pNCL and pCHR conjugation frequency decreased, while pCLR conjugation increased in frequency. (h) Cell type frequencies among plasmid-harboring recipients resulting from matings of homoplasmic donors to the novel and ancestral allele in equal frequency with plasmid-free recipients. Four independent replicates per model plasmid are shown (total tested colonies are shown below in brackets). Heteroplasmic recipients occurred at low frequency in two of four pNCL replicates, three out of four pCLR replicates and all pCHR replicates, consistent with occasional acquisition events from multiple donors.

To quantify the effect of conjugation on the novel allele dynamics we compared the recipient allele composition to that of the donor population. Here we examine the usefulness of two different measures based either on the donor cell type or the donor allele pool. The cell-based recipient to donor ratio, the frequency of homoplasmic novel allele hosts in the recipient population (nominator) was divided by the frequency of novel allele carrying donor cell (denominator). In the allele-based ratio, the denominator was the frequency of the novel allele in the total donor plasmid pool. The cell-based ratio is expected to be 1 if transfer probability of the novel allele from novel allele carries in the donor population is independent of the donor cell type (i.e., homo- or hetero-plasmic for the novel allele). The allele-based ratio is expected to be 1, if plasmid allele acquisition depends only on the relative frequency of novel and ancestral alleles in the donor population.

The cell-based recipient to donor ratio for pNCL and pCHR were lower at transfer 1 than at subsequent time points, whereas those for pCLR remained stable throughout the experiment. The ratio remained below 1 for pNCL and pCHR and mostly exceeded 1 for pCLR. The allele-based ratios were stable over time: for pNCL, novel allele hosts in the recipient population were approximately twofold less frequent than expected (Fig. 3d), and for pCHR the ratio was similarly low (Fig. 3f). In contrast, the allele-based ratio for pCLR averaged 2.24 ± 0.50 (95% CI) over the course of the experiment (Fig. 3e). Within each model plasmid, there was no significant difference between the cell-based and allele-based ratios (using t-test on the pooled data, α=0.05). Both recipient to donor ratios were significantly different from 1 for all model plasmids (using t-test on the pooled data, α=0.01). The two ratio measures are expected to be similar when the frequency of heteroplasmic donors is low. Indeed, the allele-based ratio is slightly closer to 1 in day 1 compared to the cell-based ratio, however, there is no significant different between the two measures also for that day in all model plasmids (using t-test on the pooled data, α=0.01). Note that both ratios were stable over time, despite variation in the conjugation frequencies among individual matings and decline in the mean conjugation frequency towards the end of the experiment (Fig. 3g). Taken together, the novel allele frequency in the recipient population does not reflect directly the donor cell types or allele composition. Furthermore, the novel allele frequency in the recipient population seems closer to the donor plasmid pool composition compared to the donor cell type composition.

To further examine the occurrence of heteroplasmic hosts in the recipient population, we focused on the first evolution experiment day, where the frequency of heteroplasmic donors was the highest. Phenotypic screening using streak assays revealed segregating heteroplasmic hosts in the recipient population at low frequencies, with 1/300 colonies for pNCL, 3/300 colonies for pCLR and 0/900 colonies for pCHR (Fig. S12b). Heteroplasmic recipients can arise either from the transfer of two plasmids from a single donor or plasmid transfer from multiple donors into the same recipient. Multiple plasmid transfers may occur putatively either before or after plasmid establishment. For all model plasmids, the frequency of plasmid transfer into plasmid-carrying host cells was ca. two orders of magnitude lower than transfer into plasmid-free cells (Fig. S12c). Consequently, double plasmid transfer may occur prior to plasmid establishment within the host in our experimental system. To distinguish between plasmid transfer from a single or multiple donors, we performed matings with an equal mixture of donors that were homoplasmic for either allele. These assays yielded an average (±SD) of 2.3 ± 2.9% heteroplasmic recipients for pNCL, 3.0 ± 2.2% for pCLR and 3.0 ± 2.2 % (Fig. 3h). Based on the pooled evolution experiment data, the probability of double plasmid transfer events (including transfers of identical alleles) was estimated at 0.11 for pCLR and 0.026 of pNCL conjugation events. No heteroplasmic cells in recipients were detected for pCHR, indicating that the probability of double plasmid transfer was estimated to be <0.07 of conjugation events (Fig. S12b). Taken together, plasmid recipients were predominantly homoplasmic and the proportion of recipient cells carrying the novel allele was slightly lower than the corresponding donor novel allele frequency for pNCL and pCHR, and slightly higher for pCLR.

### Conjugation modulates allele segregation dynamics in a PCN-dependent manner with minimal long-term effects

To predict the effect of conjugation on plasmid allele dynamics beyond the duration of the evolution experiment, we performed simulations of plasmid allele dynamics. For this purpose, we extended a model of random plasmid segregation to include plasmid conjugation (Fig. 4a; Supplementary Text S2)^7,9^. Plasmid conjugation was modeled using the classic mass-action kinetics framework^31^. In the model, plasmid transfer occurs initially at a rate of *βDR*, where *β* is the plasmid-specific transfer-rate constant, and *D* and *R* denote the abundances of donor and plasmid-free recipient cells, respectively. Recipient cells that have received a plasmid (denoted *T*) are also assumed to later conjugate at a rate *βTR*. In each conjugation event, a single plasmid copy is randomly selected from the donor’s plasmid pool and transferred to a plasmid-free recipient cell. Under this assumption, plasmid recipients always become homoplasmic, and the probability that a recipient inherits the ancestral or the novel allele is proportional to the respective allele frequency in the donor cell. In subsequent model extensions, we incorporate plasmid loss and allow the transfer of multiple plasmid copies per conjugation event. To describe allele segregation arising from conjugation and random plasmid assortment during cell division, we track the abundances of recipient and donor cells – including homoplasmic and heteroplasmic types – in continuous time^9^.

**Figure 4.**
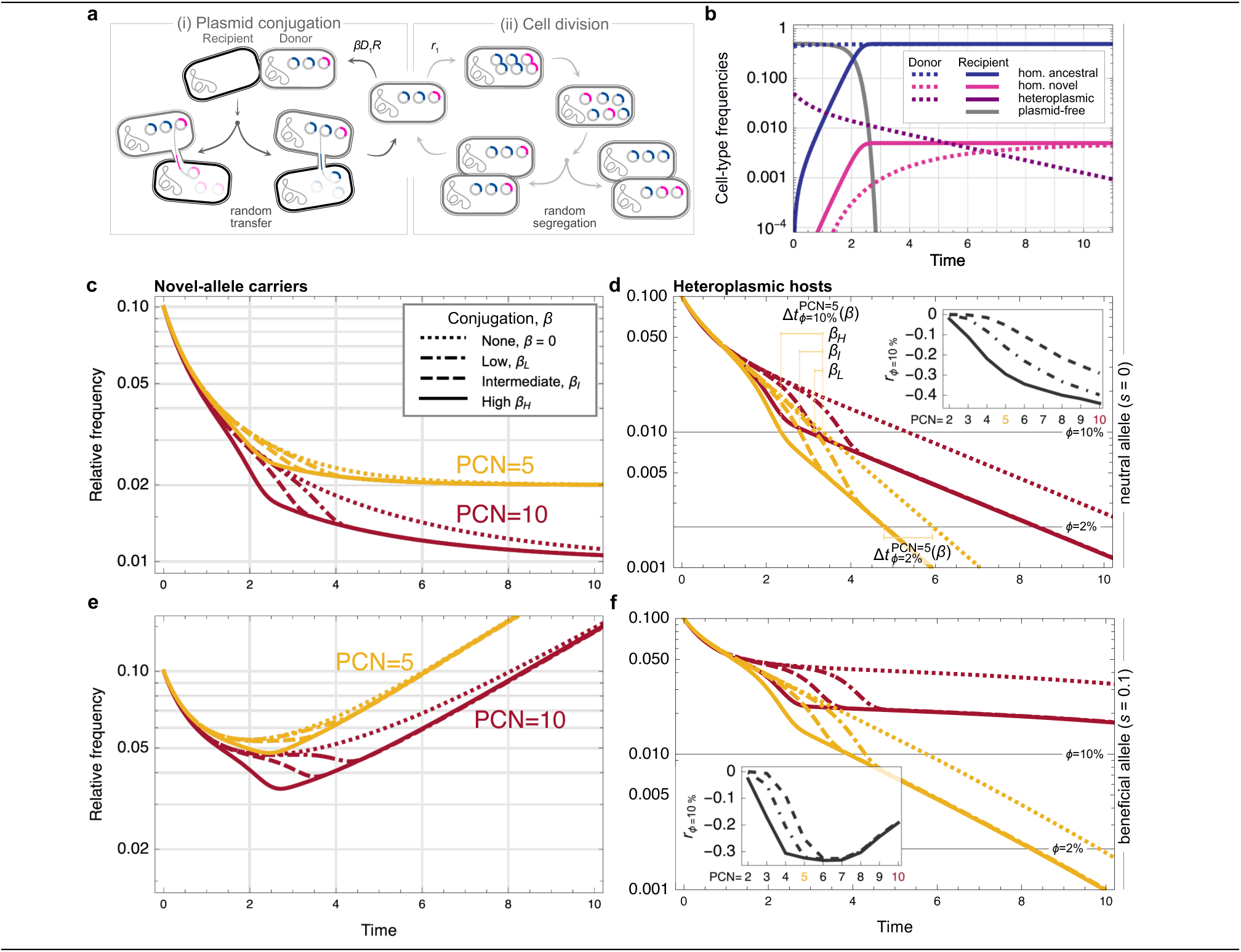
A calibrated mathematical model predicts rapid allele segregation in the presence of conjugation. (a) Plasmid segregation and transfer in the mathematical model. (i) Transfer of a single plasmid copy during conjugation generates homoplasmic cells for either the novel or the ancestral allele. (ii) Prior to cell division, all plasmid copies are replicated and randomly distributed in equal numbers to the two daughter cells, resulting in stochastic plasmid compositions. (b) Frequency of donor and recipient cell types in the course of simulated plasmid evolution for plasmid copy number *n_PCN_* = 10 with (i) a high transfer-rate constant (*β_H_* = 10^−10^) or (ii) no conjugation. (c, e) Relative frequency of novel-allele carriers, defined as the proportion of plasmid-host cells (donors and recipients) carrying at least one plasmid copy with the novel allele (d), (f) Relative frequency of heteroplasmic hosts, defined as the proportion of plasmid-host cells carrying both alleles. Timeline segments indicate the absolute reduction in segregation time due to conjugation, Δ*t*_*ϕ*_(β), for different transfer-rate constants (*β_H_*, *β_I_* = 10^−11^, *β_L_* = 10^−12^) and plasmid copy number *n_PCN_* = 5. Inset: Relative reduction in segregation time due to conjugation across combinations of plasmid copy number (PCN) and conjugation strength. Parameters: Cell division rates *r_i_* = *r* ≔ log_2_(4) for novel allele carriers and *r*_0_ = *r*(1 − *s*) for other cells. Initial frequency of plasmid-host cells carrying the novel allele in one plasmid copy *f* = 0.1. Initial population size of donors and recipients *D* = *R* = 2 × 10^7^.

The coupled dynamics of cell growth and plasmid conjugation were simulated for plasmids with copy numbers *n*_PCN_ = 5 (pNCL, pCLR) or 10 (pCHR). Estimates of the transfer rate constant *β* for the three plasmids range between 9 × 10^−12^ and 8 × 10^−11^; thus, we conducted simulations with three transfer-rate constants of similar orders of magnitude: high (*β_H_* = 10^−10^), intermediate (*β_I_* = 10^−11^), and low (*β_L_* = 10^−12^). We compared the results to the dynamics without conjugation (*β* = 0). Simulations were initialized with equal abundances of plasmid-free recipients and plasmid-carrying donors, *R*_0_ = *D*_0_ = 5 × 10^−7^ as in the conjugation assay. Similarly to the evolution experiment, a small fraction of donor cells (*f*_0_ = 0.1) was initially heteroplasmic, carrying the novel allele in a single plasmid copy, while the remaining donors were homoplasmic for the ancestral allele. The cell division rate was set to, *r* = ln(4), for both donor and recipient cells, which yields four cell divisions – as occurred during the experimental matings – per time unit. For beneficial alleles, homoplasmic ancestral and plasmid-free cells were assumed to divide at a reduced rate *r*(1 − *s*), where *s* > 0 is the selection coefficient.

Our model describes the conversion of plasmid-free cells into novel or ancestral homoplasmic recipients in a ratio reflecting the novel-allele frequency *f*/*n*_PCN_. The abundance of homoplasmic novel recipients exceeded that of homoplasmic novel donors for neutral and slightly beneficial alleles because single-plasmid conjugation produces exclusively homoplasmic recipients, whereas segregational drift generates homoplasmic novel donors only rarely (Fig. 4b). In case of neutral alleles, the relative frequency of novel allele carriers among plasmid hosts declined as heteroplasmic cells were converted into homoplasmic cells through cell division and conjugation (Fig. 4c,d). For neutral alleles, the relative frequency of novel-allele carriers converged to the initial novel-allele frequency, *f*/*n*_PCN_ = 0.01, independent of whether conjugation occurred. Note that the equilibrium is reached once all plasmid-host cells have become homoplasmic. Conjugation accelerated allele segregation – with stronger effects as the transfer-rate coefficient increases – by continuously adding homoplasmic recipients in the host population, thereby accelerating the decline of novel allele carriers. To quantify the influence of conjugation on allele segregation dynamics, we defined the segregation time, *t*_*ϕ*_(*β*), as the time at which the relative frequency of heteroplasmic hosts declines to a fraction *ϕ* of its initial value. For example, for *ϕ* = 10% (Fig. 4d), the heteroplasmic cell frequency decreases from 0.1 to 0.01 at *t*_*ϕ*_(*β*) as heteroplasmic cells are converted into homoplasmic cells. We then quantified the effect of the transfer-rate constant, *β*, by calculating the relative change in segregation time,

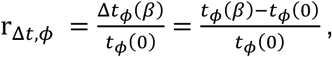

where *t*_*ϕ*_(0) denotes the segregation time in the absence of conjugation (see timeline segments in Fig. 4d). Negative values of r_Δ*t*,*ϕ*_ indicate that conjugation accelerates segregation. Segregation occurred earlier for intermediate to high plasmid copy numbers (*n*_PCN_ > 2) and sufficiently large transfer rate coefficient (Fig. 4d, inset). By contrast, for low PCNs or weak conjugation, r_Δ*t*,*ϕ*=10%_remained close to zero, indicating little influence of conjugation. For smaller thresholds *ϕ*, the absolute differences in segregation times Δ*t*_*ϕ*_(β) became less dependent on the transfer rate coefficient, *β*. For *ϕ* = 2%, there is no observable effect of the transfer rate coefficient *β* on Δ*t*_*ϕ*_ (Fig. 4d), indicating that segregation can be accelerated even when conjugation is weak. However, the relative change in segregation time, |r_Δ*t*,*ϕ*_|, decreased for very small thresholds *ϕ*, because the absolute segregation time increases as the threshold *ϕ* becomes smaller (Fig. S13; cf. *ϕ* = 10% and *ϕ* = 2% in Fig. 4d).

For intermediate positive selection for the novel allele, we found that segregational drift can transiently reduce the relative frequency of novel allele carriers (Fig. 4e, S14). The effect was stronger as the transfer-rate coefficients increased with a more reduced frequency of novel-allele carriers compared to no conjugation. Similar to the influence of conjugation on the segregation of neutral alleles, the rise of novel allele carriers was delayed by conjugation with the magnitude of delay depending also on the plasmid copy number. In contrast to neutral alleles, the relative change in segregation time of beneficial alleles reached its minimum for intermediate plasmid copy numbers caused by the slow decline of heteroplasmic hosts for high plasmid copy numbers (Fig. 4f). For strongly beneficial alleles, conjugation showed little influence on allele dynamics because plasmid-free cells proliferate slowly; the low abundance of plasmid-free cells limited the frequency of conjugation events that would otherwise affect the allele dynamics (Fig. S15).

To examine the effect of double plasmid transfer, we extended our model to allow transfer of multiple plasmid copies from single or multiple donor cells. Simulating up to 10% of two simultaneous plasmid transfers into the same recipient yielded dynamics similar to single-copy transfer. However, under extreme scenarios in which many plasmid copies were transferred during conjugation – particularly from multiple donors into a single recipient – allele segregation was slower and even temporarily reversed (Fig. S16). Note that these predictions do not take into account the negative fitness effect of multiple plasmid transfer into a single recipient due to genotoxic stress^17^.

Under faithful vertical plasmid transfer, our model assumes no influence of conjugation on the allele segregation in the donor population. Yet, two of the model plasmids we examined – pNCL and pCLR – are unstable (Fig. 1b). Plasmid loss generates plasmid-free cells that can become recipients in later conjugation. At the same time, allele segregation is slowed down in unstable plasmids, as a heteroplasmic (mother) cell remains heteroplasmic, thereby suppressing segregational drift. Simulations including plasmid loss showed that allele segregation is initially slow; However, once plasmid-free cells accumulated due to segregational loss, conjugation into plasmid-free cells substantially speeded up the allele segregation process (Fig. S17).

## Discussion

Multicopy extrachromosomal genetic elements are widespread across the tree of life, ranging from prokaryotic plasmids and viruses to eukaryotic organelles. Fundamental principles governing the evolution of these elements appear to be shared across prokaryotic and eukaryotic systems. Plasmids, viruses, and mitochondrial DNA (mtDNA) commonly occur in multiple copies per cell, where their multicopy state increases mutational supply and generates intracellular genetic diversity (heteroplasmy)^32,33^. The establishment and fixation of novel alleles in multicopy genetic elements are constrained by segregational drift, which arises from repeated bottlenecks affecting the intracellular pool of genetic elements during cell division^3,4^. Horizontal transfer is a well-known property of conjugative plasmids and viruses, whereas evidence for intercellular transfer of mtDNA is accumulating^34–36^, including observations in cancer tissues^37^. Our study provides empirical evidence and a theoretical framework demonstrating the influence of DNA transfer on allele segregation dynamics in multicopy genetic systems.

The maintenance of the novel allele in all three model conjugative plasmids exceeded the expectations derived from theoretical predictions (Fig. S9b). Previous studies employing a non-mobile plasmid with higher copy number (PCN=15) reported rapid decline in the frequency of heteroplasmic hosts under non-selective conditions^38,39^. The prolonged heteroplasmy state observed here therefore suggests the operation of additional mechanisms influencing plasmid allele segregation. Transient plasmid fusion into heteromultimers and fission into monomers represent one possible explanation and may arise through homologous recombination between plasmid variants^40,41^. Nonetheless, documentation of multimeric conjugative plasmids remains limited in sequenced genomes^42,43^, potentially reflecting detection constraints associated with standard sequencing approaches. Exploration of plasmid fusion–fission dynamics identified parameter regimes compatible with extended heteroplasmy (Fig S9c), suggesting that transient fusion events escaping our detection may contribute to the observed extended heteroplasmy. Such dynamics are plausible for our model RK2-derivative plasmids, pCLR and pCHR, that encode a multimer resolution system that may permit reversible plasmid multimerization^24^. In contrast, multimer resolution system has not been described for pHHV216^23^ and we did not observe size heterogeneity in the ancestral pNCL. Taken together, transient plasmid fusion and fission are unlikely to occur at sufficient frequency to sustain heteroplasmy in the donor populations.

We next considered whether plasmid backbone properties, i.e., active partition system and replication dynamics, could contribute to the maintenance of heteroplasmy. Plasmid RK2 harbors a tightly regulated active partition system that promotes plasmid clusters and faithful plasmid partition into daughter cells^44–46^. The novel allele dynamics observed for pCLR-Δ*incC* (Fig. S10) indicate that this mechanism is unlikely to explain the observed allele dynamics (Fig. S9b). Nonetheless, we cannot fully exclude an interaction of pCLR with chromosomally encoded partition system, as previously reported for RK2 in *Pseudomonas putida*^47^. Given the tight regulation of plasmid replication, we next considered whether replication dynamics influence the maintenance of heteroplasmy in our experimental system. Indeed, our results indicate a role of the *oriV* and the replication machinery in the persistence of heteroplasmy and the novel allele. However, heteroplasmy was maintained more efficiently than predicted by regular replication models, suggesting the operation of additional mechanisms. In particular, elements in the *oriV* region of pNCL may contribute to prolonged heteroplasmy. Deviation from random plasmid segregation may occur due to linkage of plasmid monomers^9,38^. Indeed, linkage of plasmid monomers via a handcuffing mechanism has been proposed for RK2, in which the replication control protein TrfA may bind two plasmid molecules simultaneously^48,49^. Such physical coupling could, in principle, generate deviations from random segregation, either by linking plasmids carrying different alleles or through synergistic interactions with active partitioning systems that promote the separation of sister replicons into daughter cells.

Plasmid transfer from heteroplasmic donors during the evolution experiment predominantly gave rise to homoplasmic recipients. This result is consistent with the workings of entry exclusion systems in blocking the entry of multiple plasmids copies into recipient cells^50^. The rare heteroplasmic recipients we observed likely emerged following the transfer of two plasmid copies into a single recipient prior to plasmid establishment and expression of entry exclusion systems. This interpretation is compatible with known kinetics of entry exclusion, which becomes effective about 20 min after F plasmid entry into the recipient cell^18^. The novel allele frequency in the donor plasmid pool was reflected in recipient populations, albeit with certain deviations: in the pNCL and pCHR recipient populations the novel allele was underrepresented, whereas in the pCLR recipient population the novel allele was overrepresented. These differences are unlikely to result from variation in conjugation frequencies among plasmids or alleles (Fig. 1c, S12a). Instead, allele-specific fitness effects may contribute to these deviations, as homoplasmic novel allele hosts exhibited a slight fitness advantage for pCLR but not for pNCL or pCHR (Fig. 1c). Together, these observations suggest that deviations from donor allele frequencies primarily reflect differences in donor and recipient growth dynamics modulated by plasmid-associated fitness effects.

Previous theoretical studies predicted that resistance evolution in conjugative plasmids is governed by an interplay between mutational supply, plasmid vertical transfer (inheritance) and horizontal dissemination in the host population^51,52^. Segregational drift during plasmid inheritance imposes a fundamental constraint on establishment of alleles in multicopy plasmids^4,5^. Our results show that conjugation partially alleviates this constraint by increasing the abundance of cells that are homoplasmic for the novel allele. However, the probability that a novel allele successfully disseminates to a recipient cell decreases with increasing plasmid copy number. Indeed, our results show that the allele fixation probabilities in recipients scaled proportionally with allele frequencies in the donor plasmid pool. Conjugation thus bypasses the slow emergence of homoplasmic novel cells during plasmid inheritance, and the fitness consequences of this process depend on the historical contingency. Accelerated allele segregation by conjugation may transiently reduce the proportion of novel allele carriers, and thereby lower the establishment probability of resistance-conferring alleles under strong population bottlenecks. In contrast, conjugation may provide a direct route to cells that are homoplasmic for novel alleles conferring dose-dependent beneficial traits (e.g., efflux-mediated resistance^5,53^), thereby facilitating rapid bacterial adaptation. Under conditions that induce balancing selection for the novel and ancestral plasmid alleles (termed also heterozygote advantage; e.g., combined antibiotic therapy^54^), conjugation is expected to have a negative fitness effect due to its contribution to the reduction in the proportion of heteroplasmic cells.

Plasmid evolutionary success is shaped by interactions at multiple levels, including within-host competition between plasmid variants^10,55^. Example is plasmid variants exploiting functions encoded by co-resident plasmids for their replication or transfer. These include satellite plasmid variants devoid of essential replication components that can persist only in a heteroplasmic state with the ancestral plasmid supplying the missing functions^56^. Functional interactions in the context of plasmid transfer are exemplified by high copy number plasmid variants associated with elevated conjugation frequencies due to increased expression level of the transfer machinery^57^; ancestral plasmids coexisting with such high-copy variants in heteroplasmic cells may experience enhanced transfer frequency. Under conditions permitting conjugation, plasmid variants having a low fitness due to reduced ability to replicate or transfer are expected to be purged more rapidly from populations. Rapid segregation of plasmid variants via conjugation can facilitate early escape from intracellular competition between plasmid variants exhibiting a functional interaction. By modulating allele segregation and constraining functional interactions, conjugation thus emerges as a determinant of plasmid genome evolution.

## Methods

### Strains and culturing conditions

All experiments were performed using the model strain *Acinetobacter baylyi* BD413 and BD4 (DSM 588 and DSM 586; German Collection of Microorganisms and Cell Cultures, DSMZ). A complete list of strains is provided in Table S1. Bacterial cultures were routinely grown at 30°C in liquid lysogeny broth (LB) medium with agitation at 750 rpm or on LB agar plates. When required, media were supplemented with antibiotics at the following final concentrations: spectinomycin (20 µg/ml), kanamycin (10 µg/ml), trimethoprim (50 µg/ml), gentamicin (5 µg/ml), and ampicillin (200 µg/ml). GFP expression was induced by addition of 0.5 mM IPTG. During stability assays, fitness competitions, and allele dynamics experiments (donor cultures and matings), DNase I (0.1 mg/ml) was included to prevent an effect of natural transformation on allele frequency^38^. Cloning procedures were performed in *Escherichia coli* DH5α^58^. *E. coli* cultures were grown in LB at 37°C, shaking at 160 rpm and supplemented with antibiotics as appropriate.

### Plasmid and strain construction

Primer sequences and detailed cloning procedures are listed in Supplementary Tables S2 and S3. To eliminate natural transformation during mating experiments, designated recipients were made transformation-deficient by deletion of the *comFECB* gene cluster (*comB*-*F,* DNA binding and translocation) or deletion of *dprA* (prepares incoming DNA for recombination). The recipient BD4 Δ*comB-F* strain (*comB-F*::*dhfr-1*) was generated by transforming BD4 with genomic DNA from strain KOM130 (*trpE27 rpoB1 alkM*::*nptII’*::*tg4 comB-F*::*dhfr-1*)^59^, followed by trimethoprim selection. Correct allelic replacement was confirmed by colony PCR. Cotransformation of unwanted markers from KOM130 was excluded phenotypically (rpoB^wt^ (Rif^S^) and *alkM^+^* (growth with hexadecane as sole carbon source)). Strain BD4 Δ*dprA* (Fig. S11b) was constructed by natural transformation of the parental strain BD4 by a PCR product containing the Δ*dprA::aacC1* allele (selection for gentamicin resistance) from *A. baylyi* strain NH24 (BD413 Δ*dprA*::*aacC1)*^59^.

Model plasmids pNCL and pCLR were constructed from pHHV216^23^ and RK2^24^, respectively, in *A. bayly*i, where all subsequent genetic modifications were also performed. Mobile genetic elements were deleted in pHHV216/pNCL using a two-step allelic exchange strategy with pGT42-derived suicide vectors (see below) In the model plasmid RK2/pCLR, IS*21* and *aphA*, conferring kanamycin resistance, were replaced with *aadA*, conferring spectinomycin resistance, by allelic replacement using a three-fragment overlap PCR construct (Table S3). Subsequently the genetic system to introduce a novel allele by natural transformation was inserted^38^ into both model plasmid backbones. The ancestral cassette (AD-*gfp*) comprised the lac repressor gene *lacI^adi^* driven by the *lacI*^q^ promoter and *gfpmut3.1* (*gfp*) driven by the P*_trc_* promoter, with *gfp* inserted into the middle of *nptII*, thereby disrupting the gene and conferring kanamycin sensitivity. The novel cassette (AD-*nptII*) encoded *lacI^adi^* and the intact *nptII* gene. The *lacI^adi^*variant was chosen to minimize clustered plasmid segregation. The AD-*nptII* cassette was first inserted into the Tn*1* locus of RK2 and downstream of the gene *V216_45* in pHHV216 via SOE-PCR and natural transformation. Transformants were selected on kanamycin and verified by PCR and Sanger sequencing. The AD-*gfp* cassette was then delivered by naturally transforming the AD-*nptII*-carrying variants of RK2 and pHHV216 in *A. baylyi* via a PCR product of the AD-*gfp* cassette. Successful integration of gfp was identified by green fluorescence and kanamycin sensitivity of resulting transformants.

Construction of pNCL, pCHR, pCLR-Δ*incC* and pNCL::*ori*_pTAD_ was performed using the gene-targeting vector pGT42^60^ in a two-step allelic exchange strategy. Homologous regions flanking the plasmid target locus to be removed or modified were inserted upstream and downstream of an *nptII*-*sacB* cassette (conferring kanamycin resistance and sucrose sensitivity) in the vector using Gibson Assembly^61^. *A. baylyi* cells carrying the respective target plasmid were then naturally transformed with the vector containing the *nptII-sacB* cassette, and double-crossover integrants were selected on kanamycin. To excise the *nptII-sacB* cassette from the resulting target plasmids, the cassette was removed from the vector, leaving only the flanking homologous regions, and this vector was used as donor DNA for natural transformation. Recombinants in which the cassette had been excised by homologous recombination were selected on LB plates supplemented with 7% sucrose. Sucrose-resistant, kanamycin-sensitive colonies were screened by colony PCR and confirmed by sequencing. As pGT42 does not replicate in *A. baylyi*, curing of that plasmid was not required. Alternatively, overlap-PCR was performed on the fragments fusing them without constructing a cloning vector. Then the PCR product was used as template for natural transformation of *A. baylyi*. Point mutations were introduced by PCR-mediated site-directed mutagenesis. Template plasmids (pLH13 and pLH14) for allele introduction were generated by amplifying 7.7 kb (pCLR derivatives) or 8.7 kb (pNCL derivatives) regions surrounding the *nptII* locus and cloning into pJET1.2 (Table S3).

Natural transformation of *A. baylyi* during cloning work was performed using either 1:40 diluted competent cells prepared as described^62^ or bacterial stationary cultures were initially diluted 1:4. Template DNA (≤1 ng) was added, followed by incubation at 30°C for 90 min, shaking at 750 rpm before plating on selective media.

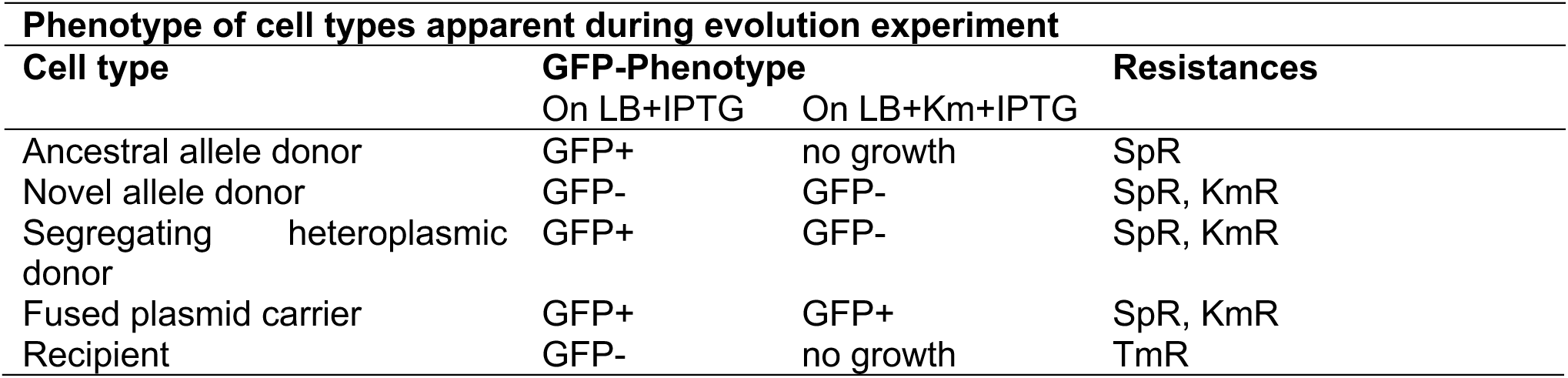

### Plasmid Isolation

Cloning vectors and template plasmids were isolated using the Monarch Spin Miniprep Kit (NEB) according to the manufacturer’s instructions. Model plasmids (pNCL, pCLR, and derivatives) were purified from 300 ml overnight cultures using the NucleoBond Xtra Maxi kit (Macherey-Nagel) with modifications optimized for low-copy plasmids. Briefly, cell pellets were resuspended in Buffer S1 supplemented with lysozyme (1 mg/ml final concentration) and incubated at 37°C for 60 min prior to alkaline lysis. After clarification by centrifugation (24,446 × g, 45 min), the supernatant was filtered before loading onto AX500 columns. DNA was eluted with prewarmed (50°C) N5 buffer.

### Plasmid stability assay

Plasmid stability was assessed using the donor genotype as the host. Six replicate cultures were serially transferred daily (1:100 bottleneck) for up to 7 days (∼47 generations). At each time point, cultures were plated on LB+IPTG for total viable counts. Plasmid hosts were quantified by scoring GFP⁺ colonies and their frequency calculated relative to total viable counts.

### Fitness experiments

Plasmid fitness effect was quantified via competitions of BD413 strains carrying pNCL, pCLR, or pCHR against plasmid-free strains. Overnight cultures were inoculated from the same competent stocks used for the evolution experiment. Cultures of plasmid-free BD4 were inoculated from single colonies. For competitions involving homoplasmic novel allele plasmid variants (pNCL::*nptII*, pCLR::*nptII* and pCHR::*nptII*), cultures were inoculated from selective plates. Overnight cultures were adjusted to an OD600 of 1.6, mixed at a 1:1 ratio of both competitors, diluted 1:100 into 1.1 ml LB in 24-well plates, and propagated for 3 days by daily transfers with a 1:100 bottleneck. At competition start and every 24 h, strain frequencies were determined by differential plating. M9 medium supplemented with 2% citrate was used to select plasmid-free competitors. Hosts of the novel allele plasmids were scored after plating on LB plates supplemeted with kanamycin. Hosts of the ancestral allele plasmids were scored after plating on LB+IPTG.

Competitions were initiated with eight biological replicates. Replicates with initial competitor ratio below 0.6 or above 1.66 were excluded from further analysis. Selection coefficients were calculated for each replicate^63,64^, and relative fitness values were inferred as previously described^65^.

### Establishment of ddPCR protocol for evaluation of donor population

A multiplex ddPCR assay was established to quantify genotype frequencies within donor populations, analyzing up to ∼6 × 10^3^ cells per reaction. In brief, PCR reaction mixtures containing template molecules were partitioned into up to 20,000 droplets, resulting in stochastic target distribution according to Poisson statistics^66^. Endpoint fluorescence of each droplet was measured following multiplex PCR using TaqMan chemistry^67^.

To resolve plasmid genotype composition, one primer-probe set was designed for each plasmid allele: A11 targeting *gfp* (ancestral allele) and A2 targeting *nptII* (novel allele) (Table S4). Two chromosomal loci, *dnaX* (A3) and *rpsR* (A4), separated by 438 kb and previously classified as essential^68^, were included to identify droplets containing intact cells. Because DNA fragmentation during sample preparation is expected to predominantly affect extracellular DNA, intact cells were defined as droplets positive for both chromosomal markers, whereas fragmented chromosomal DNA would segregate into separate droplets. All probes were labeled with distinct fluorophores to enable simultaneous multiplex detection. For genotype determination, only droplets positive for both chromosomal markers were considered intact cells. Among these, droplets lacking plasmid targets were classified as plasmid-free cells; droplets positive for either A11 or A2 alone were classified as homoplasmic for the respective allele; and droplets positive for both plasmid targets were classified as heteroplasmic (Fig. S2, left). To determine allele frequencies within the total plasmid pool independent of cellular allele composition, samples were subjected to controlled lysis prior to droplet generation. Droplets positive for both chromosomal markers were excluded from analysis to avoid masking effects from multiple plasmid copies per cell (Fig. S2, left).

Primer and probe concentrations as well as annealing temperatures were optimized using plasmid-free cells (BD413) and cells harboring both alleles on each plasmid (ABLH044: BD413 pCHR-[*sacB*-*nptII*]). The established conditions (400 nM primers, 100 nM probes) yielded stable cluster separation with minimal background signal (Fig. S3). Reactions were prepared using ddPCR™ Supermix for Probes (No dUTP; Bio-Rad) in 25 µl volumes containing 2.5 × 10^3^ to 6 × 10^3^ target molecules. Droplets were generated using the QX200™ Droplet Generator (Bio-Rad). Thermal cycling was performed as follows: 95°C for 10 min; 40 cycles of 95°C for 30 s and 59°C for 1 min (ramp rate 2°C/s); followed by 98°C for 10 min and cooling to 4°C. Droplets were analyzed using the QX600™ Droplet Reader. Fluorescence amplitude data were exported and analyzed with RStudio (version 2023.03.1+446). Thresholds were set manually relative to negative clusters. Samples subjected to allele frequency analysis were lysed prior to ddPCR using a modified lysozyme-based protocol. Samples resuspended in T4 Lysozyme Lysis Buffer were thawed and supplemented with NEBExpress T4 Lysozyme (10 µg/ml final concentration), incubated on ice for ∼6 h, stored at -20°C overnight, and incubated again on ice for ∼6 h. Samples were stored once more at -20°C prior to use. Immediately before ddPCR, lysates were diluted 10^-4^ in ddH_2_O and 2.5 µl were added per reaction (1.5 µl for pCHR to account for its higher plasmid copy number).

### Initial validation of ddPCR as method to evaluate novel allele frequencies in donors

The established ddPCR protocol was validated by comparing the quantified frequency of novel allele carriers in pNCL and pCLR host populations to the observed phenotypes (Fig. 1d). Competent stocks of BD413 pNCL and BD413 pCLR were prepared as described above, transformed with the respective template (pLH13 or pLH14), treated with DNase I after 90 min to remove residual DNA, diluted 1:10, and grown for 24 h. Approximately 2.5 × 10^3^ cells per replicate were subjected to ddPCR. In parallel, cultures were plated to determine total viable counts (LB) and the frequency of novel allele carriers (LB+Km). Among droplets with two chromosomal markers present, the ones additionally positive for A2 (*nptII*) were classified as cells carrying the novel allele. Frequencies obtained by ddPCR and plating were calculated independently for each replicate by dividing novel allele positive counts by total viable counts. No novel allele signal was detected in control cultures lacking template DNA by either method.

### Plasmid copy number determination

Plasmid copy number (PCN) was determined using ddPCR. Overnight cultures (2 ml) grown in LB supplemented with spectinomycin (20 µg/ml) were harvested at stationary phase and resuspended in T4 Lysozyme Lysis Buffer. Samples were processed using lysis protocol described above. Reactions included primer-probe sets A3 (*dnaX*), A4 (*rpsR*), and A11 (*gfp*). PCN was calculated for each replicate as the ratio of plasmid target counts (A11) to the mean chromosomal target counts (A3 and A4). Measurements were performed in four biological replicates for pNCL and pCLR and in two replicates for pCHR and pCLR::*oriV*_pTAD_.

### Experimental evolution experiment

The ancestral donor population was prepared by transformation of the donor population with the respective novel allele template. Competent cells of *A. baylyi* BD413 carrying pNCL, pCLR, pCHR, pCLR-Δ*incC*, or pNCL::*oriV*_pTAD_ were prepared as described^62^. Competent stocks were thawed and diluted in prewarmed LB to approximately 2.5 × 10^8^ cells/ml. Template DNA was added at a final concentration of 100 ng/ml (30 ng/ml for pCHR). Six independent replicates were prepared per model plasmid. An additional control culture was incubated without template DNA.

Cultures were incubated for 90 min at 30°C, shaking at 750 rpm to allow integration of the novel allele via recombination. Subsequently, DNase I was added to a final concentration of 0.1 mg/ml to degrade residual extracellular DNA. Total viable counts were determined by plating appropriate dilutions on LB+IPTG. Successful transformants were quantified from LB+Km+IPTG plates and fused plasmid carriers scored from GFP^+^ colonies under kanamycin selection.

Following the 90 min incubation, cultures were diluted 1:10 into individual wells of a 24-well plate containing 1.1 ml LB per well and incubated for 24 h at 30°C, shaking at 250 rpm (transfer 0). Thereafter, donor populations were propagated by daily 1:100 serial transfers. Unless indicated otherwise, cultures were propagated without antibiotic selection. For pCLR-Δ*incC* and the corresponding pCLR control, spectinomycin (20 µg/ml) was included during the first 15 transfers to maintain selection for the plasmid and removed thereafter.

From transfer 1 onward, genotype frequencies within donor populations were quantified by ddPCR. Fused plasmid carriers were monitored independently by differential plating as described below. For ddPCR-based genotype determination, 2 µl of a 10^-3^ dilution of the respective culture in ddH_2_O were used per reaction. To determine allele frequencies within the total plasmid pool, 300 µl of culture were harvested daily and resuspended in T4 Lysozyme Lysis Buffer (50 mM Tris-HCl pH 7.5, 0.5 mM EDTA) prior to processing as described below.

For pNCL::*oriV*_pTAD_, genotype frequencies were determined by differential plating rather than ddPCR due to altered segregation behavior. Total viable counts were obtained on LB+IPTG, plasmid hosts on LB+Sp, homoplasmic novel allele cells as KmR GFP^-^ colonies on LB+Km+IPTG, and heteroplasmic cells as KmR GFP⁺ colonies on LB+Km+IPTG. Frequencies were calculated relative to total plasmid-host titer.

The evolution experiment was continued until heteroplasmic cells were no longer detectable by ddPCR in the donor population (for pCLR) or until genotype frequencies remained similar for three days (for pNCL and pCHR). Cultures of pNCL and pCHR donors were propagated for 10 transfers, pCLR for 8 transfers (with continuation to day 24), pNCL::*oriV*_pTAD_ for 13 transfers, and pCLR-Δ*incC* for 18 transfers (15 transfers under selection and 3 transfers without selection). Assuming constant carrying capacity, this corresponds to approximately 53-67 generations depending on plasmid type and experiment duration. For pCLR, an additional step of selection for plasmid hosts (20 µg/ml spectinomycin) was applied between day 8 and 9; results in Fig. 2c and 2d were collected prior to this step.

At each transfer, 300 µl of culture were mixed 1:1 with sterile 100% glycerol (final concentration 50%) and stored at -80°C. For pCLR (day 10) and pCHR (day 8), two clones per replicate of potential fused plasmid carriers were isolated by inoculation into spectinomycin-supplemented overnight cultures, mixed 1:1 with 100% glycerol, and stored at -80°C. Clone nomenclature indicates donor replicate followed by clone number (e.g., 3.2 denotes clone 2 from donor replicate 3).

Quantification of allele transmission into the recipient population was performed daily after the donor cultures transfer. Recipient cultures (*A. baylyi* BD4 Δ*comB-F*) were thawed 1.5 h prior to mating, diluted in prewarmed LB to approximately 2.5 × 10^8^ cells/ml, and allowed to recover for 90 min at 30°C with shaking at 750 rpm. Donor and recipient cultures were mixed at an anticipated 1:1 ratio. For each mating, 100 µl of the mixture (approximately 5 × 10^7^ donor and 5 × 10^7^ recipient cells; total ∼1 × 10^8^ cells) were spread into individual wells of 12-well plates containing 1 ml LB agar. Donor and recipient monocultures were plated in parallel as controls. Mating plates were incubated at 30°C for 3 h (pNCL), 15 h (pCHR), or 21 h (pCLR). At t_0_, serial dilutions were plated on LB+IPTG to determine total donors, on LB+Sp to quantify plasmid hosts, on LB+Tm to quantify cell titer of the recipient population, and on LB+Km+IPTG to detect novel allele carriers (all CFUs) and fused plasmid carriers (GFP^+^ CUFs). Double selection (LB+Sp+Tm+IPTG) and triple selection (LB+Sp+Tm+Km+IPTG) plates were included at t_0_ to control for conjugation occurring prior to incubation. After incubation, cells were recovered in 1 ml PBS, serially diluted (typically 10^-1^ to 10^-6^), and plated. Plasmid harboring recipients were selected on LB+Sp+Tm+IPTG. Within this population, ancestral allele recipients were identified as SpR TmR GFP⁺ colonies, homoplasmic novel allele recipients as SpR TmR GFP^-^colonies, and fused plasmid recipients as SpR TmR KmR GFP⁺ colonies under kanamycin selection. Genotype frequencies were calculated relative to total plasmid harboring recipients (CFU on LB+Sp+Tm+IPTG). To control for conjugation that may occur during double and triple selection^69^, donor and recipient monocultures were mixed 1:1 immediately prior to plating at identical cell densities and plated on all selective media. The count of colonies arising on control plates correspond to background (unplanned) conjugation events that were eventually subtracted from recipient counts. Dilutions yielding >20 background colonies were excluded from analysis. Following recovery, 300 µl of each mating culture were mixed 1:1 with sterile 100% glycerol (final concentration 50%) and stored at -80°C. For pNCL matings, two fused plasmid recipient clones per replicate were isolated by overnight growth in spectinomycin-containing LB and archived as described above.

To determine the frequency of heteroplasmic recipients, streak tests were performed on day 1 recipient populations. Frozen mating populations after incubation were thawed, appropriately diluted, and plated on LB+Sp+Tm+IPTG to select for plasmid harboring recipients. GFP⁺ colonies were subsequently streaked on LB+Km+IPTG and LB+Sp+IPTG. Clones that were SpR and GFP⁺ but kanamycin-sensitive (KmS) were scored as homoplasmic for the ancestral allele. Clones that were SpR and KmR were classified as heteroplasmic if GFP^-^ on LB+Km+IPTG and as fused plasmid carriers if GFP⁺ on LB+Km+IPTG. For pNCL and pCLR, 50 GFP⁺ colonies per replicate were analyzed. For pCHR, 150 GFP⁺ colonies per replicate were scored to compensate for the lower novel allele frequency and to achieve comparable detection limits across backbones. To validate the assumption that GFP^-^ recipient colonies carried the novel allele, five GFP^-^ colonies per replicate were streaked on LB+Sp+Km+IPTG and LB+Sp. All tested GFP^-^ colonies were KmR.

### Validation of donor cell types using ddPCR

Further examination of donor cell type in pNCL donor populations from day 1 of the evolution experiment was performed using the established ddPCR assay (Fig. S4). Frozen donor cultures were thawed, diluted, and plated on LB+IPTG for total viable counts and the frequency of ancestral allele carriers (GFP⁺ CFUs). From each replicate, 25 GFP⁺ and 25 GFP^-^ colonies (or all available colonies if fewer were present) were streaked onto LB+Km+IPTG, LB+Sp, and LB+IPTG.

Among GFP⁺ colonies, three outcomes were possible: growth on all plates with GFP fluorescence under kanamycin selection was scored as fused plasmid carrier; growth on all plates with GFP fluorescence only on LB+IPTG was scored as heteroplasmic (non-fused); and growth only on LB+IPTG and LB+Sp was scored as homoplasmic for the ancestral allele. Streaks lacking growth or GFP signal on LB+IPTG were excluded. Among GFP^-^ colonies, growth on all plates was scored as homoplasmic for the novel allele; growth only on LB+IPTG was scored as plasmid loss. Colonies exhibiting GFP signal in any streak were excluded from analysis. Population-level genotype frequencies were calculated by weighting streak test classifications by the initial proportions of GFP⁺ and GFP^-^ colonies on LB+IPTG plates.

### Detection and quantification of fused plasmids in donor population

Frequencies of fused plasmid carriers in donor populations were determined by differential plating. For pNCL, pCLR, pCHR, and pCLR-Δ*incC*, no GFP⁺ (fluorescent) colonies were detected among kanamycin-resistant colonies immediately after allele introduction. The novel allele thus gains rapid dominance under selection for kanamycin resistance. Accordingly, flourescent colonies under kanamycin selection were interpreted as fused plasmid carriers. For pNCL, frozen donor cultures from every second day were thawed, diluted appropriately, and plated on LB+Sp to quantify total plasmid hosts and on LB+Km+IPTG to detect fused plasmid carriers. For pCLR, fused plasmid carrier frequencies were inferred from the LB+Km+IPTG and LB+Sp plates generated at t₀ of the respective mating assays. For pCHR and pCLR-Δ*incC* (including the parallel pCLR control), diluted donor cultures were plated directly on LB+Sp and LB+Km+IPTG. Frequencies were calculated as CFU (GFP⁺ on LB+Km+IPTG) divided by plasmid host CFU (LB+Sp).

### Detection and validation of plasmid fusion via linkage ddPCR

Physical linkage of the ancestral and novel alleles was performed using linkage ddPCR analysis^70^. Plasmids were isolated as described above. DNA concentrations were estimated by gel electrophoresis, diluted accordingly, and ∼1,000 target molecules (2 µl) were added per ddPCR reaction. Reactions included primer-probe sets A2 (*nptII*) and A11 (*gfp*) at concentrations described above and were amplified using the same cycling program. The expected frequency of droplets containing both plasmid targets was calculated according to Poisson statistics using λ derived from total plasmid target counts per accepted droplet. Droplets positive for both A2 and A11 were classified as double-target droplets. Observed double-target frequencies were calculated as the proportion of accepted droplets positive for both.

### Phenotypic validation of plasmid fusion

Clones suspected for the presence of fused plasmids were propagated in 2 ml LB in culture tubes at 30°C, shaking at 750 rpm and transferred every 24 h at a 1:100 dilution. At the onset of the experiment and at each subsequent transfer, cultures were diluted and plated on LB+IPTG and LB+Km+IPTG. Total viable counts were determined from CFUs on LB+IPTG. GFP⁺ colonies on LB+IPTG were scored as ancestral allele carriers. Colonies growing on LB+Km+IPTG were classified as novel allele carriers, and GFP⁺ colonies under kanamycin selection were considered as fused plasmid carriers. Frequencies were calculated relative to total viable counts.

The presence of plasmids fusion was additionally validated via conjugation assays with tested colonies as the donors. Matings were conducted as described for the allele dynamics assay. For pCLR and pCHR, strain BD4 Δ*comB-F* served as recipient. For pNCL, strain BD4 Δ*dprA* was used as recipient, and gentamicin was applied instead of trimethoprim to select for plasmid harboring recipients. Fused plasmid donor strains were inoculated 1:40 from frozen stocks into LB supplemented with spectinomycin to maintain backbone selection. Strain BD4 Δ*dprA* was inoculated from a single colony streaked from strain stocks. Cultures of BD4 Δ*comB-F* were prepared as described for the evolution experiment. Overnight cultures were harvested by centrifugation, washed in PBS, and donor and recipient cultures were mixed at a 1:1 ratio. Subsequent mating, recovery, and plating steps followed the evolution experiment mating protocols.

### Gel Electrophoresis of pNCL plasmid isolates

Clones suspected for the presence of fused plasmdis were propagated in 300 ml LB supplemented with spectinomycin, and plasmids were isolated as described above. Plasmid preparations were analyzed by native agarose gel electrophoresis (0.6% agarose in TBE, unstained). DNA quantities were normalized prior to loading. Gels were run at 27 V for 52 h at 4°C, stained with ethidium bromide for 20 min, destained for 10 min, and imaged using a Bio-Rad imaging system.

### Detection of heteroplasmic plasmid hosts by streak assay

Streak tests were performed to distinguish fused plasmid carriers from heteroplasmic cells carrying monomeric plasmid configurations (i.e., non-fused plasmids) that segregate over time. GFP⁺ colonies from day 17 of the pCLR donor cultures were picked from LB+IPTG plates and streaked onto LB+Km+IPTG and LB+IPTG. Colonies growing on LB+Km+IPTG without GFP signal but retaining GFP signal on LB+IPTG were classified as segregating heteroplasmic cells. Colonies growing on both plates and GFP⁺ under kanamycin selection were classified as fused plasmid carriers (Fig. S.6a).

Two segregating heteroplasmic clones per donor replicate were resuspended in LB (∼10^6^ CFU/ml), serially diluted, and plated on LB+IPTG, LB+Km+IPTG, and LB+Sp+IPTG. Total plasmid host titer was determined from CFU on LB+Sp. The titer of ancestral allele carriers was determined as GFP⁺ colonies on LB+IPTG. Novel allele carriers were quantified as GFP^-^ colonies on LB+Km+IPTG. Fused plasmid carriers were quantified as GFP⁺ colonies on LB+Km+IPTG. Proportions of each genotype were calculated relative to total plasmid hosts (CFU on LB+Sp).

To quantify genotype composition within GFP⁺ donor colonies, 50 GFP⁺ colonies per replicate were streaked onto LB+Km+IPTG and LB+IPTG. Growth on both plates with GFP signal under kanamycin selection was scored as fused plasmid carrier. Growth on both plates with GFP signal only on LB+IPTG was scored as segregating heteroplasmic cells. Growth exclusively on LB+IPTG was scored as homoplasmic ancestral allele carrier. Streaks lacking growth on LB or lacking GFP signal on LB+IPTG were excluded from further analysis.

For each replicate, proportions of each genotype within GFP⁺ colonies were estimated by dividing counts of each category divided by the total number of valid streaks. Population-level frequencies were calculated by multiplying the cell type proportion by the frequency of ancestral allele carriers in the respective donor population. The frequency of heteroplasmic cells was subtracted from total novel allele carriers to infer the frequency of homoplasmic novel allele cells. In the visualizations the proportion of plasmid-hosts was set to 1.

### Conjugation frequency of novel and ancestral alleles

Overnight cultures of homoplasmic ancestral and homoplasmic novel allele plasmid hosts, as well as a BD4 Δ*comB-F* recipient population, were grown under selective conditions. Cultures were adjusted to an OD600 of 1.6 and subjected to mating assays following the protocol described for the evolution experiment. Conjugation frequencies were calculated relative to the product of total donor and recipient population counts.

### Quantifying multiple plasmid transfer via mating assays

To distinguish single-donor from multiple-donor transfer events, additional mating assays were performed for pNCL, pCLR, and pCHR. Homoplasmic donors carrying either the ancestral or the novel allele were mixed 1:1 (four replicates per model plasmid) and subsequently combined 1:1 with the recipient population. Mating conditions, and data evaluation followed the protocol described for the evolution experiment. After incubation, colonies were documented on LB+Tm+Sp+IPTG plates including GFP fluorescence. Plates were subsequently replica plated onto LB+Km^71^. For pCHR ca. 100 colonies were streak tested instead on LB+Km and LB+Sp+IPTG. The overlay of both plates was used to identify GFP⁺ colonies capable of growth under Km selection, indicative of heteroplasmic seeding cells. Frequencies of plasmid harbouring recipients were calculated relative to the total number of colonies on LB+Tm+Sp+IPTG plates.

To further test whether an established plasmid backbone in the recipient permits secondary acquisition of the same backbone carrying a different allele, plasmid harboring recipients of the ancestral allele (Δ*comB-F* pNCL, Δ*comB-F* pCLR or Δ*comB-F* pCHR) were mated 1:1 with homoplasmic novel allele donors of the respective backbone (pNCL::*nptII*, pCLR::*nptII* or pCHR::*nptII*) in four independent replicates. For comparison control mating assays consisted of homoplasmic novel allele donors or homoplasmic ancestral allele donors mated 1:1 with plasmid-free recipients (two replicates each). Mating conditions, and data evaluation were identical to the ones above with the following alteration: plasmid-harboring recipients were selected for by gentamicin instead of trimethoprim.

### Estimation of the double plasmid conjugation probability

The probability of two different plasmid variants being received by a recipient cell during conjugation is given by

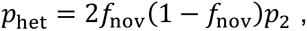

assuming binomial sampling from the donor plasmid pool, where *f*_nov_ denotes the novel allele frequency in the donor plasmid pool and *p*_2_ is the double plasmid conjugation probability. We infer the double transfer probability (*p*_2_) by the measured frequency of heteroplasmic recipients *r*_het_ and the novel allele frequency,

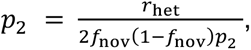

where we set the probability of transfer of two different plasmid variants into one recipient to the frequency of heteroplasmic recipients in the hosts of the recipient population, *r*_het_ = *p*_het_.

## Acknowledgements

We thank Holger Heuer (Julius Kühn-Institute, Quedlinburg, Germany) for kindly providing pHHV216. We thank Johanna E. Trapp for technical assistance during experimental work. We thank Florence Muccino, Alex Gashanja, Jordi Weijers and Hieu Minh Truong for critical comments on the manuscript. This research was supported in part through high-performance computing resources available at the Kiel University computing center.

## Data availability statement

The data underlying this article are available in the article and in its online supplementary information,

## Code availability statement

The scripts required to reproduce the simulations can be found in the GitHub repositories https://github.com/mariosanter/conPlasDyn-conjugation and https://github.com/mariosanter/conPlasDyn-heteroplasmy and were archived on Zenodo.

## Author contributions

L.H., N.H. and T.D. conceived the study and designed the evolution experiment. L.H. performed the experiments and analyzed their results. L.H., M.S. and T.D. conceived model simulations. M.S. designed and performed model simulations. All authors wrote the manuscript.

## References

1. Castañeda-Barba, S., Top, E.M., and Stalder, T. (2024). Plasmids, a molecular cornerstone of antimicrobial resistance in the One Health era. Nat. Rev. Microbiol. 22, 18–32. 10.1038/s41579-023-00926-x.

2. Palomino, A., Gewurz, D., DeVine, L., Zajmi, U., Moralez, J., Abu-Rumman, F., Smith, R.P., and Lopatkin, A.J. (2023). Metabolic genes on conjugative plasmids are highly prevalent in *Escherichia coli* and can protect against antibiotic treatment. ISME J. 17, 151–162. 10.1038/s41396-022-01329-1.

3. San Millan, A., Escudero, J.A., Gifford, D.R., Mazel, D., and MacLean, R.C. (2016). Multicopy plasmids potentiate the evolution of antibiotic resistance in bacteria. Nat. Ecol. Evol. 1, 10. 10.1038/s41559-016-0010.

4. Ilhan, J., Kupczok, A., Woehle, C., Wein, T., Hülter, N.F., Rosenstiel, P., Landan, G., Mizrahi, I., and Dagan, T. (2019). Segregational drift and the interplay between plasmid copy number and evolvability. Mol. Biol. Evol. 36, 472–486. 10.1093/molbev/msy225.

5. Santer, M., and Uecker, H. (2020). Evolutionary rescue and drug resistance on multicopy plasmids. Genetics 215, 847–868. 10.1534/genetics.119.303012.

6. Garoña, A., Hülter, N.F., Romero Picazo, D., and Dagan, T. (2021). Segregational Drift Constrains the Evolutionary Rate of Prokaryotic Plasmids. Mol. Biol. Evol. 38, 5610–5624. 10.1093/molbev/msab283.

7. Garoña, A., Santer, M., Hülter, N.F., Uecker, H., and Dagan, T. (2023). Segregational drift hinders the evolution of antibiotic resistance on polyploid replicons. PLOS Genet. 19, e1010829. 10.1371/journal.pgen.1010829.

8. Ramiro-Martínez, P., Quinto, I.D., Jaraba-Soto, L., Lanza, V.F., Herencias-Rodríguez, C., González Casanova, A., Peña-Miller, R., and Rodríguez-Beltrán, J. (2026). Plasmid mutation rates scale with copy number. Proc. Natl. Acad. Sci. 123, e2526088123. 10.1073/pnas.2526088123.

9. Santer, M., Kupczok, A., Dagan, T., and Uecker, H. (2022). Fixation dynamics of beneficial alleles in prokaryotic polyploid chromosomes and plasmids. Genetics 222, iyac121. 10.1093/genetics/iyac121.

10. Rossine, F., Sanchez, C., Eaton, D., Paulsson, J., and Baym, M. (2025). Intracellular competition shapes plasmid population dynamics. Science 390, eadx0665. 10.1126/science.adx0665.

11. Maddamsetti, R., Shyti, I., Wilson, M.L., Son, H.-I., Baig, Y., Zhou, Z., Lu, J., and You, L. (2025). Scaling laws of bacterial and archaeal plasmids. Nat. Commun. 16, 6023. 10.1038/s41467-025-61205-2.

12. Ramiro-Martínez, P., de Quinto, I., Lanza, V.F., Gama, J.A., and Rodríguez-Beltrán, J. (2025). Universal rules govern plasmid copy number. Nat. Commun. 16, 6022. 10.1038/s41467-025-61202-5.

13. Lawley, T., Wilkins, B.M., and Frost, L.S. (2004). Bacterial conjugation in Gram-negative bacteria. In Plasmid Biology (John Wiley & Sons, Ltd), pp. 203–226. 10.1128/9781555817732.ch9.

14. Virolle, C., Goldlust, K., Djermoun, S., Bigot, S., and Lesterlin, C. (2020). Plasmid transfer by conjugation in Gram-negative bacteria: from the cellular to the community level. Genes 11, 1239. 10.3390/genes11111239.

15. Skurray, R.A., and Reeves, P. (1973). Characterization of lethal zygosis associated with conjugation in *Escherichia coli* K-12. J. Bacteriol. 113, 58–70. 10.1128/jb.113.1.58-70.1973.

16. Garcillán-Barcia, M.P., and de la Cruz, F. (2008). Why is entry exclusion an essential feature of conjugative plasmids? Plasmid 60, 1–18. 10.1016/j.plasmid.2008.03.002.

17. Couturier, A., Fraikin, N., and Lesterlin, C. (2025). Exclusion systems preserve host cell homeostasis and fitness, ensuring successful dissemination of conjugative plasmids and associated resistance genes. Nucleic Acids Res. 53, gkaf898. 10.1093/nar/gkaf898.

18. Couturier, A., Virolle, C., Goldlust, K., Berne-Dedieu, A., Reuter, A., Nolivos, S., Yamaichi, Y., Bigot, S., and Lesterlin, C. (2023). Real-time visualisation of the intracellular dynamics of conjugative plasmid transfer. Nat. Commun. 14, 294. 10.1038/s41467-023-35978-3.

19. Prensky, H., Gomez-Simmonds, A., Uhlemann, A.-C., and Lopatkin, A.J. (2021). Conjugation dynamics depend on both the plasmid acquisition cost and the fitness cost. Mol. Syst. Biol. 17, e9913. 10.15252/msb.20209913.

20. Hall, J.P.J., Williams, D., Paterson, S., Harrison, E., and Brockhurst, M.A. (2017). Positive selection inhibits gene mobilization and transfer in soil bacterial communities. Nat. Ecol. Evol. 1, 1348–1353. 10.1038/s41559-017-0250-3.

21. Lopatkin, A.J., Huang, S., Smith, R.P., Srimani, J.K., Sysoeva, T.A., Bewick, S., Karig, D., and You, L. (2016). Antibiotics as a selective driver for conjugation dynamics. Nat. Microbiol. 1, 16044. 10.1038/nmicrobiol.2016.44.

22. Overballe-Petersen, S., Harms, K., Orlando, L.A.A., Mayar, J.V.M., Rasmussen, S., Dahl, T.W., Rosing, M.T., Poole, A.M., Sicheritz-Ponten, T., Brunak, S., et al. (2013). Bacterial natural transformation by highly fragmented and damaged DNA. Proc. Natl. Acad. Sci. 110, 19860–19865. 10.1073/pnas.1315278110.

23. Heuer, H., Kopmann, C., Binh, C.T.T., Top, E.M., and Smalla, K. (2009). Spreading antibiotic resistance through spread manure: characteristics of a novel plasmid type with low %G+C content. Environ. Microbiol. 11, 937–949. 10.1111/j.1462-2920.2008.01819.x.

24. Pansegrau, W., Lanka, E., Barth, P.T., Figurski, D.H., Guiney, D.G., Haas, D., Helinski, D.R., Schwab, H., Stanisich, V.A., and Thomas, C.M. (1994). Complete nucleotide sequence of Birmingham IncP alpha plasmids. Compilation and comparative analysis. J. Mol. Biol. 239, 623–663. 10.1006/jmbi.1994.1404.

25. Sia, E.A., Roberts, R.C., Easter, C., Helinski, D.R., and Figurski, D.H. (1995). Different relative importances of the par operons and the effect of conjugal transfer on the maintenance of intact promiscuous plasmid RK2. J. Bacteriol. 177, 2789–2797. 10.1128/jb.177.10.2789-2797.1995.

26. Durland, R.H., Toukdarian, A., Fang, F., and Helinski, D.R. (1990). Mutations in the trfA replication gene of the broad-host-range plasmid RK2 result in elevated plasmid copy numbers. J. Bacteriol. 172, 3859–3867. 10.1128/jb.172.7.3859-3867.1990.

27. Roberts, R.C., Ström, A.R., and Helinski, D.R. (1994). The *parDE* operon of the broad-host-range plasmid RK2 specifies growth inhibition associated with plasmid loss. J. Mol. Biol. 237, 35–51. 10.1006/jmbi.1994.1207.

28. Motallebi-Veshareh, M., Rouch, D.A., and Thomas, C.M. (1990). A family of ATPases involved in active partitioning of diverse bacterial plasmids. Mol. Microbiol. 4, 1455–1463. 10.1111/j.1365-2958.1990.tb02056.x.

29. Lukaszewicz, M., Kostelidou, K., Bartosik, A.A., Cooke, G.D., Thomas, C.M., and Jagura-Burdzy, G. (2002). Functional dissection of the ParB homologue (KorB) from IncP-1 plasmid RK2. Nucleic Acids Res. 30, 1046–1055. 10.1093/nar/30.4.1046.

30. Verheust, C., and Helinski, D.R. (2007). The *incC korB* region of RK2 repositions a mini-RK2 replicon in *Escherichia coli*. Plasmid 58, 195–204. 10.1016/j.plasmid.2007.03.004.

31. Levin, B.R., Stewart, F.M., and Rice, V.A. (1979). The kinetics of conjugative plasmid transmission: fit of a simple mass action model. Plasmid 2, 247–260. 10.1016/0147-619X(79)90043-X.

32. Birky, C.W. (2001). The inheritance of genes in mitochondria and chloroplasts: laws, mechanisms, and models. Annu. Rev. Genet. 35, 125–148. 10.1146/annurev.genet.35.102401.090231.

33. Rodríguez-Beltrán, J., DelaFuente, J., León-Sampedro, R., MacLean, R.C., and San Millán, Á. (2021). Beyond horizontal gene transfer: the role of plasmids in bacterial evolution. Nat. Rev. Microbiol. 19, 347–359. 10.1038/s41579-020-00497-1.

34. Plotnikov, E.Y., Khryapenkova, T.G., Vasileva, A.K., Marey, M.V., Galkina, S.I., Isaev, N.K., Sheval, E.V., Polyakov, V.Y., Sukhikh, G.T., and Zorov, D.B. (2008). Cell-to-cell cross-talk between mesenchymal stem cells and cardiomyocytes in co-culture. J. Cell. Mol. Med. 12, 1622–1631. 10.1111/j.1582-4934.2007.00205.x.

35. Hayakawa, K., Esposito, E., Wang, X., Terasaki, Y., Liu, Y., Xing, C., Ji, X., and Lo, E.H. (2016). Transfer of mitochondria from astrocytes to neurons after stroke. Nature 535, 551–555. 10.1038/nature18928.

36. Borcherding, N., and Brestoff, J.R. (2023). The power and potential of mitochondria transfer. Nature 623, 283–291. 10.1038/s41586-023-06537-z.

37. Garcia-Souto, D., Bruzos, A.L., Diaz, S., Rocha, S., Pequeño-Valtierra, A., Roman-Lewis, C.F., Alonso, J., Rodriguez, R., Costas, D., Rodriguez-Castro, J., et al. (2022). Mitochondrial genome sequencing of marine leukaemias reveals cancer contagion between clam species in the Seas of Southern Europe. eLife 11, e66946. 10.7554/eLife.66946.

38. Garoña, A., Hülter, N.F., Romero Picazo, D., and Dagan, T. (2021). Segregational drift constrains the evolutionary rate of prokaryotic plasmids. Mol. Biol. Evol. 38, 5610–5624. 10.1093/molbev/msab283.

39. Hernandez-Beltran, J.C.R., Rodríguez-Beltrán, J., Millán, A.S., Peña-Miller, R., and Fuentes-Hernández, A. (2021). Quantifying plasmid dynamics using single-cell microfluidics and image bioinformatics. Plasmid 113, 102517. 10.1016/j.plasmid.2020.102517.

40. Hülter, N.F., Wein, T., Effe, J., Garoña, A., and Dagan, T. (2020). Intracellular competitions reveal determinants of plasmid evolutionary success. Front. Microbiol. 11. 10.3389/fmicb.2020.02062.

41. Summers, D.K., Beton, C.W.H., and Withers, H.L. (1993). Multicopy plasmid instability: the dimer catastrophe hypothesis. Mol. Microbiol. 8, 1031–1038. 10.1111/j.1365-2958.1993.tb01648.x.

42. Abe, R., Akeda, Y., Sugawara, Y., Matsumoto, Y., Motooka, D., Kawahara, R., Yamamoto, N., Tomono, K., Iida, T., and Hamada, S. (2021). Enhanced carbapenem resistance through multimerization of plasmids carrying carbapenemase genes. mBio 12, 10.1128/mbio.00186-21. https://doi.org/10.1128/mbio.00186-21.

43. Hanke, D.M., and Dagan, T. (2025). SegMantX: a novel tool for detecting dna duplications uncovers prevalent duplications in plasmids. Mol. Biol. Evol. 42, msaf242. 10.1093/molbev/msaf242.

44. Saurugger, P.N., Hrabak, O., Schwab, H., and Lafferty, R.M. (1986). Mapping and cloning of the *par*-region of broad-host-range plasmid RP4. J. Biotechnol. 4, 333–343. 10.1016/0168-1656(86)90047-7.

45. Davis, T.L., Helinski, D.R., and Roberts, R.C. (1992). Transcription and autoregulation of the stabilizing functions of broad-host-range plasmid RK2 in *Escherichia coli*, *Agrobacterium tumefaciens* and *Pseudomonas aeruginosa*. Mol. Microbiol. 6, 1981–1994. 10.1111/j.1365-2958.1992.tb01371.x.

46. Pogliano, J., Ho, T.Q., Zhong, Z., and Helinski, D.R. (2001). Multicopy plasmids are clustered and localized in *Escherichia coli*. Proc. Natl. Acad. Sci. U. S. A. 98, 4486–4491. 10.1073/pnas.081075798.

47. Kolatka, K., Witosinska, M., Pierechod, M., and Konieczny, I. (2008). Bacterial partitioning proteins affect the subcellular location of broad-host-range plasmid RK2. Microbiology 154, 2847–2856. 10.1099/mic.0.2008/018762-0.

48. Bury, K., Wegrzyn, K., and Konieczny, I. (2017). Handcuffing reversal is facilitated by proteases and replication initiator monomers. Nucleic Acids Res. 45, 3953–3966. 10.1093/nar/gkx166.

49. Kittell, B.L., and Helinski, D.R. (1991). Iteron inhibition of plasmid RK2 replication in vitro: evidence for intermolecular coupling of replication origins as a mechanism for RK2 replication control. Proc. Natl. Acad. Sci. 88, 1389–1393. 10.1073/pnas.88.4.1389.

50. Haase, J., Lurz, R., Grahn, A.M., Bamford, D.H., and Lanka, E. (1995). Bacterial conjugation mediated by plasmid RP4: RSF1010 mobilization, donor-specific phage propagation, and pilus production require the same Tra2 core components of a proposed DNA transport complex. J. Bacteriol. 177, 4779–4791. 10.1128/jb.177.16.4779-4791.1995.

51. Tazzyman, S.J., and Bonhoeffer, S. (2014). Plasmids and evolutionary rescue by drug resistance. Evolution 68, 2066–2078. 10.1111/evo.12423.

52. Geoffroy, F., and Uecker, H. (2023). Limits to evolutionary rescue by conjugative plasmids. Theor. Popul. Biol. 154, 102–117. 10.1016/j.tpb.2023.10.001.

53. Rodríguez-Beltrán, J., Sørum, V., Toll-Riera, M., de la Vega, C., Peña-Miller, R., and San Millán, Á. (2020). Genetic dominance governs the evolution and spread of mobile genetic elements in bacteria. Proc. Natl. Acad. Sci. 117, 15755–15762. 10.1073/pnas.2001240117.

54. Rodriguez-Beltran, J., Hernandez-Beltran, J.C.R., Delafuente, J., Escudero, J.A., Fuentes-Hernandez, A., MacLean, R.C., Peña-Miller, R., and San Millan, A. (2018). Multicopy plasmids allow bacteria to escape from fitness trade-offs during evolutionary innovation. Nat. Ecol. Evol. 2, 873–881. 10.1038/s41559-018-0529-z.

55. Effe, J., Santer, M., Wang, Y., Feenstra, T.E., Hülter, N.F., and Dagan, T. (2025). The combination of active partitioning and toxin-antitoxin systems is most advantageous for low-copy plasmid fitness. Nat. Commun. 16, 7078. 10.1038/s41467-025-62473-8.

56. Zhang, X., Deatherage, D.E., Zheng, H., Georgoulis, S.J., and Barrick, J.E. (2019). Evolution of satellite plasmids can prolong the maintenance of newly acquired accessory genes in bacteria. Nat. Commun. 10, 5809. 10.1038/s41467-019-13709-x.

57. Dimitriu, T., Matthews, A.C., and Buckling, A. (2021). Increased copy number couples the evolution of plasmid horizontal transmission and plasmid-encoded antibiotic resistance. Proc. Natl. Acad. Sci. 118, e2107818118. 10.1073/pnas.2107818118.

58. Hanahan, D. (1983). Studies on transformation of *Escherichia coli* with plasmids. J. Mol. Biol. 166, 557–580. 10.1016/S0022-2836(83)80284-8.

59. Hülter, N.F., Sørum, V., Borch-Pedersen, K., Liljegren, M.M., Utnes, A.L.G., Primicerio, R., Harms, K., and Johnsen, P.J. (2017). Costs and benefits of natural transformation in *Acinetobacter baylyi*. BMC Microbiol. 17, 34. 10.1186/s12866-017-0953-2.

60. Wein, T., Dagan, T., Fraune, S., Bosch, T.C.G., Reusch, T.B.H., and Hülter, N.F. (2018). Carrying capacity and colonization dynamics of *Curvibacter* in the hydra host habitat. Front. Microbiol. 9. 10.3389/fmicb.2018.00443.

61. Gibson, D.G., Young, L., Chuang, R.-Y., Venter, J.C., Hutchison, C.A., and Smith, H.O. (2009). Enzymatic assembly of DNA molecules up to several hundred kilobases. Nat. Methods 6, 343–345. 10.1038/nmeth.1318.

62. Kickstein, E., Harms, K., and Wackernagel, W. (2007). Deletions of *recBCD* or *recD* influence genetic transformation differently and are lethal together with a *recJ* deletion in *Acinetobacter baylyi*. Microbiol. Read. Engl. 153, 2259–2270. 10.1099/mic.0.2007/005256-0.

63. Dykhuizen, D.E., and Hartl, D.L. (1983). Selection in chemostats. Microbiol. Rev. 47, 150–168. 10.1128/mr.47.2.150-168.1983.

64. Levin, B.R., Perrot, V., and Walker, N. (2000). Compensatory mutations, antibiotic resistance and the population genetics of adaptive evolution in bacteria. Genetics 154, 985–997. 10.1093/genetics/154.3.985.

65. Starikova, I., Al-Haroni, M., Werner, G., Roberts, A.P., Sørum, V., Nielsen, K.M., and Johnsen, P.J. (2013). Fitness costs of various mobile genetic elements in *Enterococcus faecium* and *Enterococcus faecalis*. J. Antimicrob. Chemother. 68, 2755–2765. 10.1093/jac/dkt270.

66. Hindson, B.J., Ness, K.D., Masquelier, D.A., Belgrader, P., Heredia, N.J., Makarewicz, A.J., Bright, I.J., Lucero, M.Y., Hiddessen, A.L., Legler, T.C., et al. (2011). High-throughput droplet digital PCR system for absolute quantitation of DNA copy number. Anal. Chem. 83, 8604–8610. 10.1021/ac202028g.

67. Holland, P.M., Abramson, R.D., Watson, R., and Gelfand, D.H. (1991). Detection of specific polymerase chain reaction product by utilizing the 5’----3’ exonuclease activity of Thermus aquaticus DNA polymerase. Proc. Natl. Acad. Sci. 88, 7276–7280. 10.1073/pnas.88.16.7276.

68. De Berardinis, V., Vallenet, D., Castelli, V., Besnard, M., Pinet, A., Cruaud, C., Samair, S., Lechaplais, C., Gyapay, G., Richez, C., et al. (2008). A complete collection of single-gene deletion mutants of *Acinetobacter baylyi* ADP1. Mol. Syst. Biol. 4, 174. 10.1038/msb.2008.10.

69. He, Z., Smets, B.F., and Dechesne, A. (2024). Mating assay: plating below a cell density threshold is required for unbiased estimation of plasmid conjugation frequency of RP4 transfer between *E. coli* strains. Microb. Ecol. 87, 109. 10.1007/s00248-024-02427-7.

70. de la Cruz Barron, M., Kneis, D., Elena, A.X., Bagra, K., Berendonk, T.U., and Klümper, U. (2023). Quantification of the mobility potential of antibiotic resistance genes through multiplexed ddPCR linkage analysis. FEMS Microbiol. Ecol. 99, fiad031. 10.1093/femsec/fiad031.

71. Wein, T., Stücker, F.T., Hülter, N.F., and Dagan, T. (2019). Quantification of plasmid-mediated antibiotic resistance in an experimental evolution approach. JoVE J. Vis. Exp., e60749. 10.3791/60749.

