## Supplemental text for "Horizontal transfer promotes allele segregation in multicopy plasmids"

#### Contents

**S1 Model of multicopy plasmid allele dynamics extended for plasmid fusion and partitioning of sister plasmids** **2**

**S2 Population-level model of plasmid allele dynamics with conjugation** **7**

### S1 Model of multicopy plasmid allele dynamics extended for plasmid fusion and partitioning of sister plasmids

#### Model

We simulate the segregation dynamics of two neutral alleles on a non-transmissible multicopy plasmid. We extend the model of random assortment of plasmid copies at cell division introduced in Santer and Uecker (2020) for plasmid fusion and partitioning of isogenic sister plasmids. Each bacterial cell carries a plasmid at a fixed copy number $n$ at cell birth. Plasmids occur in two allelic types—ancestral and novel. At $t = 0$, a small fraction $f_0$ of cells is heteroplasmic, carrying one plasmid copy with the novel allele and $n - 1$ copies with the ancestral allele. All remaining cells are homoplasmic for the ancestral allele. Upon cell division, all plasmid copies are duplicated and subsequently distributed to the two daughter cells such that the plasmid copy number $n$ is restored in each daughter cell. In the baseline model, individual plasmid copies are independently and randomly assorted between daughter cells (Fig. S9A(i)). Like in the implementation in Garoña et al. (2023), we consider a cell population of infinite size. Moreover, here we also neglect bottlenecks and growth saturation (see also Santer et al. (2022) for a related framework).

We extend the baseline model by incorporating independently two mechanisms that promote the maintenance of heteroplasmy: plasmid dimerization, resulting in fused-plasmid carriers (Fig. S9A(ii)) and partitioning of sister plasmids (Fig. S9A(iii)). For plasmid fusion, we assume that (randomly formed) pairs of monomeric plasmids form dimers at probability $d$, which results in heteromultimers or homomultimers depending on the involved plasmids. Conversely, plasmid dimers resolve into monomers at probability $d'$, which we assume to occur prior to dimerization in our model. At the start of the simulation, all plasmids are assumed to be in a monomeric form. For the sister-partitioning mechanism, we assume that sister plasmids——the two copies produced by replication of a single plasmid——are actively segregated into different daughter cells with probability $a$. With complementary probability $1 - a$, plasmids are assorted randomly.

#### Results

Predicting the allele segregation under the (baseline) random assortment model underestimated the maintenance of heteroplasmy in the pNCL plasmid (Fig. S9B(i), plasmid copy number $n = 5$) The maintenance of the pNCL::*ori*$_{\text{pTAD}}$ plasmid employed with the origin of replication from the pTAD matched the prediction from the random assortment model for the first 3–5 transfers (Fig. S9B(ii), see also results in the main text).

Plasmid fusion slows the loss of heteroplasmy by reducing the number of independently segregating plasmid units. Partitioning of sister plasmids counteracts random assortment by ensuring that homogenic plasmid copies are distributed across daughter cells and, thus, hindering allele segregation. Hence, both mechanisms extend the persistence of heteroplasmic cells compared to the baseline random assortment model with temporal dynamics depending on the model parameters $d$, $d'$ and $a$ as well the plasmid copy number $n$ (Fig. S9C).

We compared the simulated allele segregation dynamics under the fusion model and the sister-partitioning model with the experimental dynamics of the low-copy plasmids pNCL and pCLR (PCN=5). We used plasmid copy number $n = 4$ and $n = 6$ since these two models (plasmid fusion and sister partitioning) need an even copy number, and we set the initial proportion of heteroplasmic cells to $f_0 = 0.1$.

Under random assortment model, our mathematical analysis revealed that the simulated heteroplasmic cell frequency declines by a factor $1/n$ per generation, which corresponds to loss of 70 % heteroplasmy for copy number $n = 4$, 78 % for $n = 5$, 85 % for $n = 6$ per transfer. The simulated loss of heteroplasmy under occasional fusion and fission ($d = d' = 0.1$) is substantially slowed. While it takes approximately 1.5 transfers for a reduction of the heteroplasmic frequency of an order of magnitude under random assortment for copy number $n = 4$, occasional fusion and fission extends this duration to approximately 10 transfers (Fig. S9C, plot (ii)). Contrary to the experimental finding, most heteroplasmic cells are fused heterogenic plasmid carriers under these conditions. Hence, we simulated the segregation dynamics under frequent fission ($d' = 0.4$) with occasional and rare fusion events ($d = 0.1$, $d = 0.01$ respectively). For occasional fusion with frequent fission, the allele segregation was slightly prolonged with fused heterogenic plasmid carriers making up ca. 50 % of the heteroplasmic subpopulation (Fig. S9C, plot (iii)). Under rare fusion ($d = 0.01$), the simulated heteroplasmic frequencies resembled the relative proportion of fused plasmid carriers of ca. 10 % in pCLR and pCHR. The heteroplasmic cell frequency under rare fission, however, was very similar to the simulated dynamics under the random assortment model, thus, did not influence the heteroplasmic loss.

Simulating the allele dynamics under a model of sister-plasmid partitioning shows that an intermediate separation probability of $a = 0.4$ yield a slight prolongation of heteroplasmic maintenance similar to occasional fusion with rare fission (Fig. S9C, plot (v)). Strong partitioning of sister plasmids similar to partitioning of sister chromatides could lead to an extended maintenance of heteroplasmy (Fig. S9C, plot (vi)).

#### Mathematical description of the model

##### (i) Baseline model.

Let $N_i$ denote the abundance of cells carrying $i = 0, \dots, n$ copies of the novel allele. The temporal dynamics of the cell population are described by a system of $n + 1$ differential equations,

$$\dot{N}_i = \sum_{j=0}^{n} r_j N_j \left( m_{j \to i} - \delta_{ij} \right) \text{ for all } i, \tag{1}$$

where $r_j \equiv 1$ is the division rate of $j$-type cells, $\delta_{ij}$ is Kronecker's delta, and $N^{(c)}$ denotes the carrying capacity. The term $m_{j \to i}$ is the expected number of $i$-type daughter cells produced by the division of a $j$-type cell,

$$m_{j \to i} = 2 \frac{\binom{2j}{i} \binom{2n-2j}{n-i}}{\binom{2n}{n}}. \tag{2}$$

A detailed derivation of the baseline model is given in Garoña et al. (2023).

The approximate heteroplasmic cell loss rate can be by obtained by the dominant eigenvalue of the matrix $(m_{j \to i})_{i,j \in \{1,\dots,n-1\}}$, which is (Santer et al., 2022)

$$\frac{2n-3}{2n-1}. \tag{3}$$

Along the lines of Eq. A17 of Santer et al. (2022), the dynamics of heteroplasmic cell type frequency, $i \in \{1, \dots, n-1\}$ quickly converges to

$$\dot{x}_{\text{het}} := \frac{\sum_i \dot{N}_i}{N} = \left( \frac{2n-3}{2n-1} - 1 \right) x_{\text{het}} = -\frac{1}{n + \frac{1}{2}} x_{\text{het}} \overset{n \gg 1}{\approx} -\frac{1}{n} x_{\text{het}}. \tag{4}$$

For discrete generations, $g$, we can similarly calculate

$$x_{\text{het}}(g) = x_{\text{het}}(0)\left(1 - \frac{1}{n}\right)^{g}, \tag{5}$$

as well as, for a transfer with a bottleneck factor 100 with $g_t = \log_2(100) \approx 6.7$ generations between two transfers. Thus, the reduction of heteroplasmy per transfer can be calculated by

$$1 - \frac{x_{\text{het}}(g_t)}{x_{\text{het}}(0)} = 1 - \left(1 - \frac{1}{n}\right)^{g_t} \approx \begin{cases} 0.77 & \text{for } n = 5, \\ 0.50 & \text{for } n = 10, \end{cases} \tag{6}$$

#### (ii) Plasmid fusion and fission

To model plasmid fusion, we extend the state space to include both monomeric and dimeric plasmids carrying one or both alleles. A cell is characterized by the tuple

$$\mathbf{i} = (i_a^{(\text{m})}, i_n^{(\text{m})}, i_a^{(\text{d})}, i_n^{(\text{d})}, i_{wn}^{(\text{d})}),$$

where $i_a^{(\text{m})}$ and $i_n^{(\text{m})}$ denote the numbers of ancestral and novel monomers, $i_a^{(\text{d})}$ and $i_n^{(\text{d})}$ the numbers of homoplasmic dimers, and $i_{wn}^{(\text{d})}$ the number of heteroplasmic dimers. Dimers are counted as two plasmid copies, such that the copy number constraint becomes

$$i_a^{(\text{m})} + i_n^{(\text{m})} + 2i_a^{(\text{d})} + 2i_n^{(\text{d})} + 2i_{wn}^{(\text{d})} = n. \tag{7}$$

We compute the expected number of $\mathbf{i}$-type daughter cells produced by a $\mathbf{j}$-type cell by iteratively propagating the probability distribution through three stages: plasmid replication and dimer resolution (monomerization), dimer formation, and random assortment (Fig. 9A).

**Plasmid replication and monomerization (Fission).** All plasmid replicons—dimers and monomers–replicate once and each dimer resolves into monomers independently with probability $d'$. Let $x_a^{(\text{d}\to\text{m})}$, $x_n^{(\text{d}\to\text{m})}$, and $x_{an}^{(\text{d}\to\text{m})}$ denote the numbers of resolving ancestral, novel, and heteroplasmic dimers, respectively. These random variables are binomially distributed,

$$x_\bullet^{(\text{d}\to\text{m})} \sim \text{Bin}(2i_\bullet^{(\text{d})}, d'). \tag{8}$$

Monomerization yields an intermediate cell type

$$\begin{aligned} \mathbf{i}^{(\text{mon})} = \Big(& i_a^{(\text{m})} + 2x_a^{(\text{d}\to\text{m})} + x_{an}^{(\text{d}\to\text{m})},\ i_n^{(\text{m})} + 2x_n^{(\text{d}\to\text{m})} + x_{an}^{(\text{d}\to\text{m})}, \\ & i_a^{(\text{d})} - x_a^{(\text{d}\to\text{m})},\ i_n^{(\text{d})} - x_n^{(\text{d}\to\text{m})},\ i_{wn}^{(\text{d})} - x_{an}^{(\text{d}\to\text{m})}\Big). \end{aligned} \tag{9}$$

**Dimerization (Fusion).** Monomeric plasmids are randomly paired. The total number of distinct pairings of $N = i_a^{(\text{mon})} + i_n^{(\text{mon})}$ monomers is

$$\frac{N!}{2^{N/2}(N/2)!}. \tag{10}$$

Conditioned on the numbers of ancestral, novel, and heteroplasmic pairs, the number of configurations yielding $x_a$ ancestral, $x_n$ novel, and $x_{an}$ heteroplasmic pairs is

$$\binom{i_a^{(\text{mon})}}{2x_a}\frac{(2x_a)!}{2^{x_a}x_a!}\binom{i_n^{(\text{mon})}}{2x_n}\frac{(2x_n)!}{2^{x_n}x_n!}x_{an}!. \tag{11}$$

The ratio of (11) and (10) gives the probability of forming a given pairing configuration.

Each plasmid pair forms a dimer independently with probability $d$. Let $x_\bullet^{(\mathrm{m}\to\mathrm{d})} \sim \mathrm{Bin}(x_\bullet, d)$ denote the number of successful dimerization events. The resulting intermediate cell type is

$$\begin{aligned}\mathbf{i}^{(\mathrm{dim})} = \Big(&i_a^{(\mathrm{mon})} - 2x_a^{(\mathrm{m}\to\mathrm{d})} - x_{an}^{(\mathrm{m}\to\mathrm{d})},\ i_n^{(\mathrm{mon})} - 2x_n^{(\mathrm{m}\to\mathrm{d})} - x_{an}^{(\mathrm{m}\to\mathrm{d})},\\ &i_a^{(\mathrm{d})} + x_a^{(\mathrm{m}\to\mathrm{d})},\ i_n^{(\mathrm{d})} + x_n^{(\mathrm{m}\to\mathrm{d})},\ i_{wn}^{(\mathrm{d})} + x_{an}^{(\mathrm{m}\to\mathrm{d})}\Big). \end{aligned} \tag{12}$$

**Random assortment.** In the final step, plasmids are randomly assorted into daughter cells while preserving their dimerization state. To maintain a fixed plasmid copy number $n$, exactly $n$ plasmid copies are assigned to each daughter cell.

We compute the probability distribution of daughter cell types by sequentially removing plasmids from the post-dimerization state $\mathbf{i}^{(\mathrm{dim})} = (i_a, i_n, i_{ww}, i_{nn}, i_{wn})$ until $n$ copies remain in the cell. Here,

$$N = i_a + i_n + 2i_{ww} + 2i_{nn} + 2i_{wn}$$

denotes the total number of plasmid copies, and

$$R = i_a + i_n + i_{ww} + i_{nn} + i_{wn}$$

the total number of replicons (monomers and dimers).

If $N = n + 1$, only a single monomeric plasmid can be removed. The corresponding transition probabilities are

$$P(i_a - 1, i_n, i_{ww}, i_{nn}, i_{wn}) = \frac{i_a}{i_a + i_n} P(i_a, i_n, i_{ww}, i_{nn}, i_{wn}), \tag{13}$$

$$P(i_a, i_n - 1, i_{ww}, i_{nn}, i_{wn}) = \frac{i_n}{i_a + i_n} P(i_a, i_n, i_{ww}, i_{nn}, i_{wn}). \tag{14}$$

If $N > n + 1$, either monomers or dimers can be removed. The transition probabilities are then given by

$$P(i_a - 1, i_n, i_{ww}, i_{nn}, i_{wn}) = \frac{i_a}{R} P(i_a, i_n, i_{ww}, i_{nn}, i_{wn}), \tag{15}$$

$$P(i_a, i_n - 1, i_{ww}, i_{nn}, i_{wn}) = \frac{i_n}{R} P(i_a, i_n, i_{ww}, i_{nn}, i_{wn}), \tag{16}$$

$$P(i_a, i_n, i_{ww} - 1, i_{nn}, i_{wn}) = \frac{i_{ww}}{R} P(i_a, i_n, i_{ww}, i_{nn}, i_{wn}), \tag{17}$$

$$P(i_a, i_n, i_{ww}, i_{nn} - 1, i_{wn}) = \frac{i_{nn}}{R} P(i_a, i_n, i_{ww}, i_{nn}, i_{wn}), \tag{18}$$

$$P(i_a, i_n, i_{ww}, i_{nn}, i_{wn} - 1) = \frac{i_{wn}}{R} P(i_a, i_n, i_{ww}, i_{nn}, i_{wn}). \tag{19}$$

By iterating these transitions, we obtain the probability distribution over all daughter cell types with exactly $n$ plasmid copies. The final transition probabilities are obtained by summing over all intermediate states arising from monomerization and dimerization.

#### Partitioning of sister plasmids

Sister-plasmid partitioning is modeled as a two-step process. First, sister plasmids are selected for active segregation with probability $a$. Second, remaining plasmids are randomly assorted.

For a $j$-type cell with $j_n = j$ novel and $j_a = n - j$ ancestral plasmids, the numbers of actively partitioned sister pairs are

$$J_a^{(\mathrm{par})} \sim \mathrm{Bin}(j_a, a), \tag{20}$$

$$J_n^{(\mathrm{par})} \sim \mathrm{Bin}(j_n, a). \tag{21}$$

The remaining plasmids are randomly drawn via a hypergeometric distribution,

$$j_a^{(\mathrm{free})} = 2j_a - J_a^{(\mathrm{par})}, \tag{22}$$

$$j_n^{(\mathrm{free})} = 2j_n - J_n^{(\mathrm{par})}. \tag{23}$$

Hence, the number of ancestral-type and novel-type plasmids after cell division is,

$$I_a = J_a^{(\mathrm{par})} + \mathrm{HG}(n - J_a^{(\mathrm{par})} - J_n^{(\mathrm{par})}, J_a^{(\mathrm{free})}, J_a^{(\mathrm{free})} + J_n^{(\mathrm{free})}), \tag{24}$$

$$I_m = n - I_a, \tag{25}$$

where $p_{\mathrm{HG}}(n_{\mathrm{draw}}, n_{\mathrm{succ}}, n_{\mathrm{total}})$ denotes a hypergeometric distribution with probability mass function

$$p_{\mathrm{HG}}(k) = \frac{\binom{n_{\mathrm{succ}}}{k}\binom{n_{\mathrm{total}}-n_{\mathrm{succ}}}{n_{\mathrm{draw}}-k}}{\binom{n_{\mathrm{total}}}{n_{\mathrm{draw}}}}. \tag{26}$$

#### Computer simulations

The model was implemented in Python programming language and computer simulations were performed using a Jupyter notebook with Python (version 3.13.5). The code is available on Github (`github.com/mariosanter/conPlasDyn-heteroplasmy`) and archived using Zenodo (`https://doi.org/10.5281/zenodo.18891692`).

### S2 Population-level model of plasmid allele dynamics with conjugation

We simulate plasmid allele dynamics using a random segregation model (Garoña et al., 2023) combined with a deterministic mass-action kinetics model of plasmid conjugation. As in the section above, bacterial cells carry a conjugative plasmid with two allelic variants. Here, we neglect active partitioning and plasmid multimerization to isolate the effect of conjugation on allele dynamics. Cells carrying at least one copy of the novel allele divide at rate $r$, whereas cells lacking the novel allele divide at a reduced rate $r(1-s)$, where $s$ denotes the selection coefficient. Upon division, plasmid-host cells either produce two daughter cells that retain the parental plasmid copy number with probability $p$, or experience segregational plasmid loss, generating one plasmid-free daughter cell. Unless stated otherwise, results shown in the main text assume faithful plasmid segregation ($p = 1$).

#### Population-level conjugation dynamics

We first ignore plasmid genotype and assume that all plasmids carry the ancestral allele. We distinguish four cell types: (i) plasmid-carrying *donor* cells ($D$), (ii) plasmid-free donors that have lost the plasmid ($D_f$), (iii) plasmid-carrying *transconjugants* ($T$), and (iv) plasmid-free *recipient* cells ($R$). The temporal dynamics of their abundances are given by

$$\begin{aligned}
\dot{D} &= rpD + \beta D_f(D + t_c T), \\
\dot{D}_f &= rD_f + r(1-p)D - \beta D_f(D + t_c T), \\
\dot{T} &= rpT + \beta R(D + t_c T), \\
\dot{R} &= rR + r(1-p)T - \beta R(D + t_c T),
\end{aligned} \tag{27}$$

where $\beta$ is the conjugation transfer-rate constant (Huisman et al., 2022). The parameter $t_c \in \{0, 1\}$ indicates whether transconjugant cells are themselves competent donors ($t_c = 1$); throughout this work, we set $t_c = 1$.

#### Allele-resolved dynamics of plasmid-carrying cells

We next extend the model to include plasmid allele dynamics. Following Garoña et al. (2023), we denote by $i$ the number of novel-allele plasmid copies in a cell with total plasmid copy number $n$. Accordingly, $D_i$ and $T_i$ denote the abundances of donor and transconjugant cells carrying $i$ novel-type and $n - i$ ancestral-type plasmid copies.

In the absence of conjugation, the dynamics of donor cells are governed by random plasmid segregation during cell division:

$$\dot{D}_i = \sum_{j=0}^{n} \underbrace{r(1-\delta_{j0}s)}_{:=r_j} pD_j \left(m_{j\to i} - \delta_{ij}\right), \tag{28}$$

which is analogous to Eq. (1), but without a saturation term. Here, $m_{j\to i}$ denotes the probability that a cell with $j$ novel plasmid copies produces a daughter cell with $i$ such copies upon division. The growth rate $r_j$ reflects selection against cells lacking the novel allele ($j = 0$).

#### Conjugation dynamics with allele resolution

Plasmid conjugation between donor cells of type $i$ and plasmid-free recipients occurs at rate $\beta D_i R$. In the *baseline model*, a single plasmid copy is transferred per conjugation event. The transferred plasmid is selected uniformly at random from the donor's plasmid pool, such that the recipient receives a novel allele with probability $i/n$ and an ancestral allele with probability $1 - i/n$. After replication of the transferred plasmid, the transconjugant cell becomes homoplasmic for the corresponding allele as all plasmid copies originate from the transferred plasmid.

The dynamics of homoplasmic transconjugants carrying only ancestral ($T_0$) or only novel ($T_n$) alleles are given by

$$\dot{T}_0 = r_0 p T_0 + \beta R \left( \sum_{i=0}^{n} \left(1 - \frac{i}{n}\right) D_i + t_c T_0 \right), \tag{29}$$

$$\dot{T}_n = r_n p T_n + \beta R \left( \sum_{i=0}^{n} \frac{i}{n} D_i + t_c T_n \right). \tag{30}$$

The abundance of plasmid-free recipient cells evolves according to

$$\dot{R} = r_0 R + (1-p)(r_0 T_0 + r_n T_n) - \beta R \left( \sum_{i=0}^{n} D_i + t_c (T_0 + T_n) \right). \tag{31}$$

#### Replenishment of donor cells by conjugation

Conjugation can also replenish donor cells that have lost their plasmids. Including this process, the dynamics of donor cells of type $i$ are

$$\dot{D}_i = \sum_{j=0}^{n} r_j p D_j \left( m_{j \to i} - \delta_{ij} \right) + \beta D_f \begin{cases} \left( \sum_{j=0}^{n} D_j \left(1 - \frac{j}{n}\right) \right) + t_c T_0, & i = 0, \\ \left( \sum_{j=0}^{n} D_j \frac{j}{n} \right) + t_c T_n, & i = n. \end{cases} \tag{32}$$

Finally, the abundance of donor cells that have lost the plasmid follows

$$\dot{D}_f = \sum_{j=0}^{n} r_j (1-p) D_j + r D_f - \beta D_f \left( \sum_{i=0}^{n} D_i + t_c (T_0 + T_n) \right). \tag{33}$$

#### Initial conditions and model dimensionality

Together, the model comprises $n + 5$ coupled ordinary differential equations. We integrate them simultaneously to obtain the abundances (or relative frequencies) of homoplasmic ancestral donors ($D_0$), homoplasmic novel donors ($D_n$), heteroplasmic donors ($D_1 + \cdots + D_{n-1}$), plasmid-free donors ($D_f$), homoplasmic transconjugants ($T_0$, $T_n$), and recipient cells ($R$).

Simulations start from an initial total population size $N_0$ with $R(0) = f_R N_0$ recipient cells ($f_R = 50\,\%$ throughout this study) and $D_1(0) = f_A (1 - f_R) N_0$ donor cells carrying exactly one novel-type plasmid copy. All remaining donor cells are homoplasmic for the ancestral allele, $D_0(0) = N_0 - D_1(0) - R(0)$. All other cell types arise only through cell division and conjugation, and are initialized as $D_i(0) = 0$ for $i \geq 2$, and $T_0(0) = T_n(0) = D_f(0) = 0$.

#### Model extension for multi-copy plasmid transfer

Here, we extend the allele-dynamics model to allow for the transfer of multiple plasmid copies during conjugation. We distinguish two scenarios: (i) *single-donor conjugation*, in which several plasmid copies are transferred from a single donor cell to a recipient, and (ii) *multi-donor conjugation*, in which multiple plasmid copies may be transferred from different donor cells during a single conjugation event. To reduce complexity, we omit selection, segregational plasmid loss, and competence of transconjugant cells by setting $s = 0$, $p = 1$, and $t_c = 0$ in the following.

Under these assumptions, the temporal dynamics of donor cell types $D_i$ are governed by Eq. (28). The dynamics of plasmid-free recipient cells simplify to

$$\dot{R} = r_0 R - \beta R \left( \sum_{i=0}^{n} D_i \right) . \tag{34}$$

In contrast to the baseline model, multi-copy transfer can generate heteroplasmic transconjugants due to the simultaneous transfer of both plasmid variants. Therefore, we track the abundances of all transconjugant cell types $T_i$ with $i = 0, \ldots, n$ novel-type plasmid copies. Their temporal dynamics are given by

$$\dot{T_i} = \sum_{j=0}^{n} r T_j \left( m_{j \to i} - \delta_{ij} \right) + \beta R \sum_{j=0}^{n} q_{j \to i} D_j , \tag{35}$$

where $q_{j \to i}$ denotes the probability that conjugation involving a donor cell of type $j$ produces a transconjugant cell of type $i$. We derive these transition probabilities below.

#### Transition probabilities for multi-copy transfer

For both transfer scenarios, we denote by $p_{c^{\text{tot}}}$ the probability that a total of $c^{\text{tot}} = 1, 2, \ldots, n$ plasmid copies are transferred during a conjugation event.

**Single-donor conjugation.** For single-donor transfer, the probability that $c^{\text{nov}}$ novel-type and $c^{\text{anc}} = c^{\text{tot}} - c^{\text{nov}}$ ancestral-type plasmid copies are transferred from a donor cell of type $j$ is

$$p_{c^{\text{tot}}} \, \mathrm{HG}\big(c^{\text{nov}}; \, c^{\text{tot}}, j, n\big) , \tag{36}$$

where $\mathrm{HG}$ denotes the probability mass function of the hypergeometric distribution, reflecting sampling without replacement from the donor's plasmid pool.

Following conjugation, the transferred plasmids replicate until the total plasmid copy number $n$ is reached. Replication proceeds deterministically in complete rounds, where each plasmid copy replicates once per round, followed—if necessary—by a final partial replication round in which a subset of plasmids is selected uniformly at random for replication.

The probability that replication results in a transconjugant cell carrying $t^{\text{nov}}$ novel-type and $t^{\text{anc}} =$

$n - t^{\mathrm{nov}}$ ancestral-type plasmid copies is

$$P_{\mathrm{rep}}\big(t^{\mathrm{nov}};\, c^{\mathrm{nov}}, c^{\mathrm{tot}}\big) = \begin{cases} \delta_{t^{\mathrm{nov}},\, c^{\mathrm{nov}} 2^R}, & \text{if there exists } R \in \mathbb{N}_0 \text{ such that } n = c^{\mathrm{tot}} 2^R, \\ \dfrac{\dbinom{m^{\mathrm{nov}}}{t^{\mathrm{nov}} - m^{\mathrm{nov}}} \dbinom{m^{\mathrm{tot}} - m^{\mathrm{nov}}}{t - (t^{\mathrm{nov}} - m^{\mathrm{nov}})}}{\dbinom{m^{\mathrm{tot}}}{t}}, & \text{otherwise.} \end{cases} \tag{37}$$

Here,

$$R = \max\{r \in \mathbb{N}_0 : c^{\mathrm{tot}} 2^r \leq n\}$$

is the number of complete replication rounds,

$$m^{\mathrm{tot}} = c^{\mathrm{tot}} 2^R, \qquad m^{\mathrm{nov}} = c^{\mathrm{nov}} 2^R,$$

are the total and novel-type plasmid copy numbers after the complete rounds, and

$$t = n - m^{\mathrm{tot}}$$

is the number of plasmids produced in the final partial replication round. The second case corresponds to random selection of plasmid copies for replication during this partial round.

The resulting transition probability for the joint conjugation–replication process is

$$q_{j \to i} = \sum_{\substack{c^{\mathrm{nov}}, c^{\mathrm{anc}} \\ c^{\mathrm{nov}} + c^{\mathrm{anc}} = c^{\mathrm{tot}} \leq n}} p_{c^{\mathrm{tot}}} \, \mathrm{HG}\big(c^{\mathrm{nov}};\, c^{\mathrm{tot}}, j, n\big) \; P_{\mathrm{rep}}\big(i;\, c^{\mathrm{nov}}, c^{\mathrm{tot}}\big) \, . \tag{38}$$

**Multi-donor conjugation.** For multi-donor conjugation, we assume that plasmid copies are sampled independently from the pooled plasmid population of all donor cells. This corresponds to sampling with replacement, which is appropriate for sufficiently large donor populations. The probability that $c^{\mathrm{tot}}$ plasmid copies are transferred, of which $c^{\mathrm{nov}}$ are novel-type, is

$$p_{c^{\mathrm{tot}}} \, \mathrm{Bin}\big(c^{\mathrm{nov}};\, p = f_A,\, c^{\mathrm{tot}}\big) \, , \tag{39}$$

where

$$f_A = \frac{\sum_i i D_i}{n \sum_i D_i}$$

is the frequency of the novel allele among donor plasmids. The joint transition probabilities $q_{j \to i}$ are obtained analogously by combining this distribution with the replication kernel $P_{\mathrm{rep}}$ defined above.

#### Computer simulations

The model was implemented in the Wolfram Language and computer simulations were performed with Mathematica (version 14.0). Mathematica notebooks are available on Github (`github.com/mariosanter/conPlasDyn-conjugation`) and archived using Zenodo (`doi.org/10.5281/zenodo.18891290`).

### References

A. Garoña, M. Santer, N. F. Hülter, H. Uecker, and T. Dagan. Segregational drift hinders the evolution of antibiotic resistance on polyploid replicons. *PLOS Genetics*, 19:e1010829, 2023.

Jana S. Huisman, Fabienne Benz, Sarah J.N. Duxbury, J. Arjan G.M. de Visser, Alex R. Hall, Egil A.J. Fischer, and Sebastian Bonhoeffer. Estimating plasmid conjugation rates: A new computational tool and a critical comparison of methods. *Plasmid*, 121:102627, 2022.

Mario Santer and Hildegard Uecker. Evolutionary Rescue and Drug Resistance on Multicopy Plasmids. *Genetics*, 215(3):847–868, 2020.

Mario Santer, Anne Kupczok, Tal Dagan, and Hildegard Uecker. Fixation dynamics of beneficial alleles in prokaryotic polyploid chromosomes and plasmids. *Genetics*, 222(2):iyac121, 2022.
