## Supplementary Figures and Tables for "Horizontal transfer promotes allele segregation in multicopy plasmids"

Lisa M. Hartmann, Mario Santer, Nils F. Hülter, Tal Dagan
Institute of General Microbiology, Kiel University, Kiel, Germany

**This pdf includes:**
Supplementary Figures S1 to S17
Supplementary Table S1 to S6

### Supplementary Figures

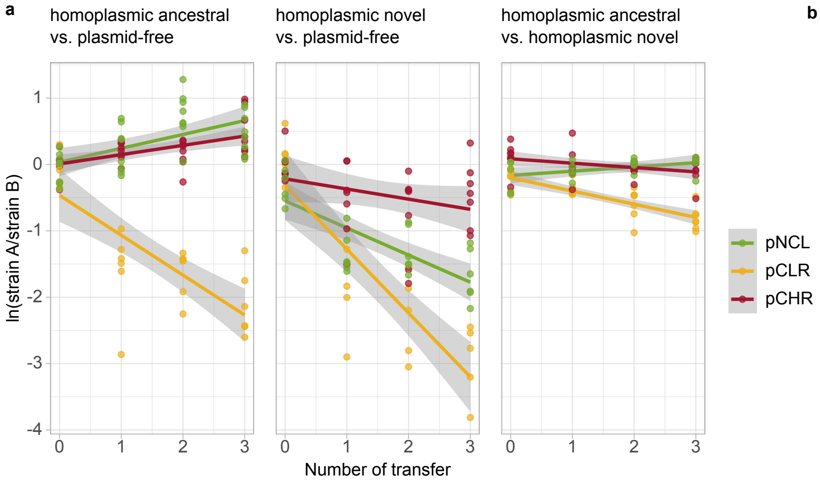


b

| competiton | model plasmid | *w* ± 95% CI |
|---|---|---|
| homoplasmic ancestral vs. plasmid-free | pNCL | 1.038 ± 0.009 |
| | pCLR | 0.856 ± 0.033 |
| | pCHR | 1.035 ± 0.017 |
| homoplasmic novel vs. plasmid-free | pNCL | 0.900 ± 0.027 |
| | pCLR | 0.778 ± 0.033 |
| | pCHR | 0.969 ± 0.027 |
| homoplasmic ancestral vs. homoplasmic novel | pNCL | 1.013 ± 0.011 |
| | pCLR | 0.957 ± 0.008 |
| | pCHR | 0.986 ± 0.004 |

**Figure S1. Linear regression of competition and relative fitness of model plasmids.**

(a) Linear regression of competition assay measurements. Points represent natural log-transformed ratios of the competing strains over time; the grey shaded region indicates the 95% confidence interval of the regression. (b) Relative fitness of model plasmids calculated from differences in the Malthusian parameter.

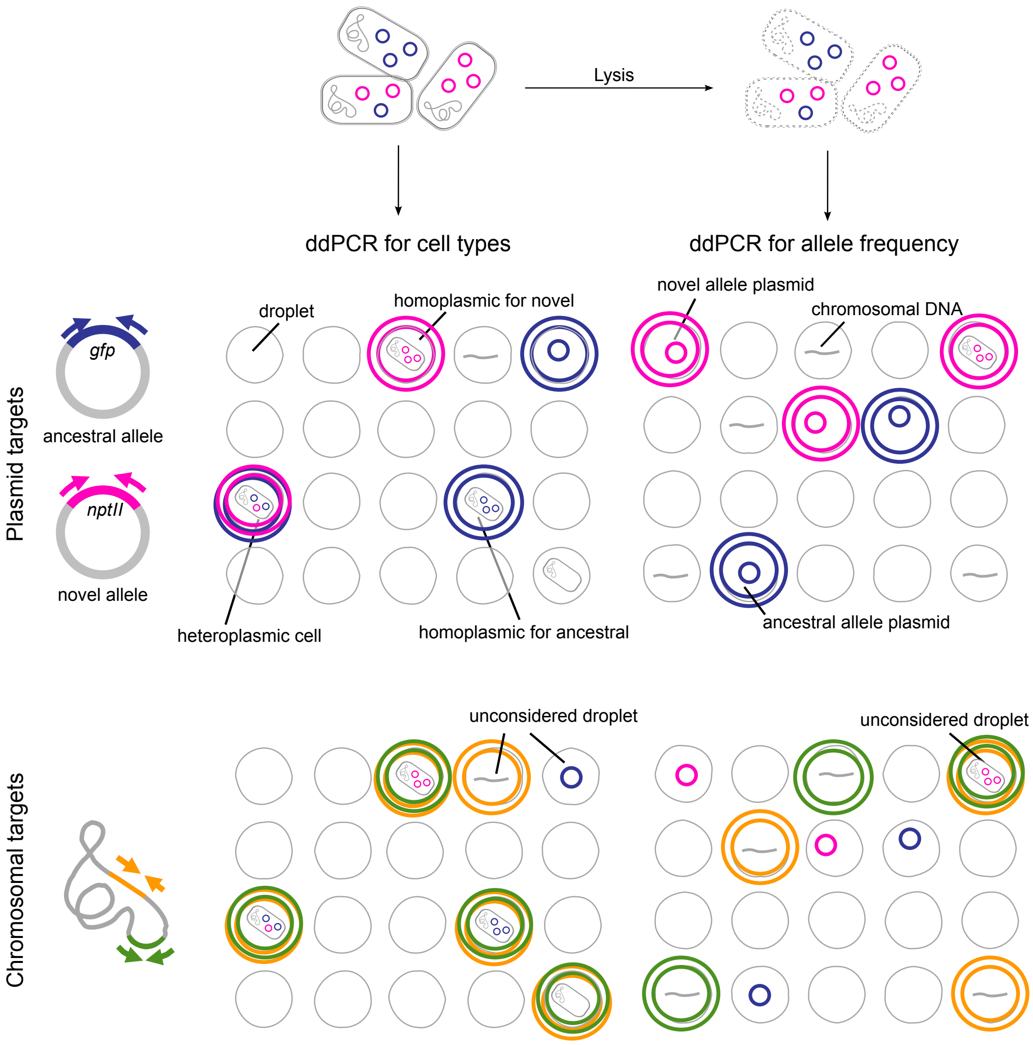


**Figure S2. ddPCR assays used during the evolution experiment.**
Schematic overview of droplet digital PCR (ddPCR) assays used to quantify cell type frequencies (left) or allele frequencies and plasmid copy number (right). Grey circles represent example droplets generated during reaction mix preparation. Colored double outlines indicate fluorescence emission detected during droplet reading: ancestral allele (blue), novel allele (pink), and the two chromosomal targets (green and orange). Droplets in the top row show fluorescence signals derived exclusively from plasmid-target probes; the bottom row shows the same droplets with fluorescence resulting from chromosomal target detection. The PCR template within each droplet may consist of either intact cells or free DNA originating from lysed cells (see labels). For determination of cell types, ddPCR was performed using untreated samples from serial donor cultures. After appropriate dilution, single cells are encapsulated into individual droplets, enabling single-cell assessment of plasmid composition. Droplets containing only one chromosomal marker and/or plasmid target(s) are interpreted as containing free DNA from lysed cells and are excluded from analysis. Droplets positive for both chromosomal markers are classified as cell-containing. Among these droplets, detection of only one allele indicates homoplasmy, whereas detection of both alleles indicates heteroplasmy. For measurements of allele frequencies in the total plasmid pool or of plasmid copy number (PCN), cells were lysed prior to addition to the ddPCR reaction mixture. In this case, droplets positive for both chromosomal markers are excluded because they may contain intact cells. The remaining droplets are analyzed to determine allele content.

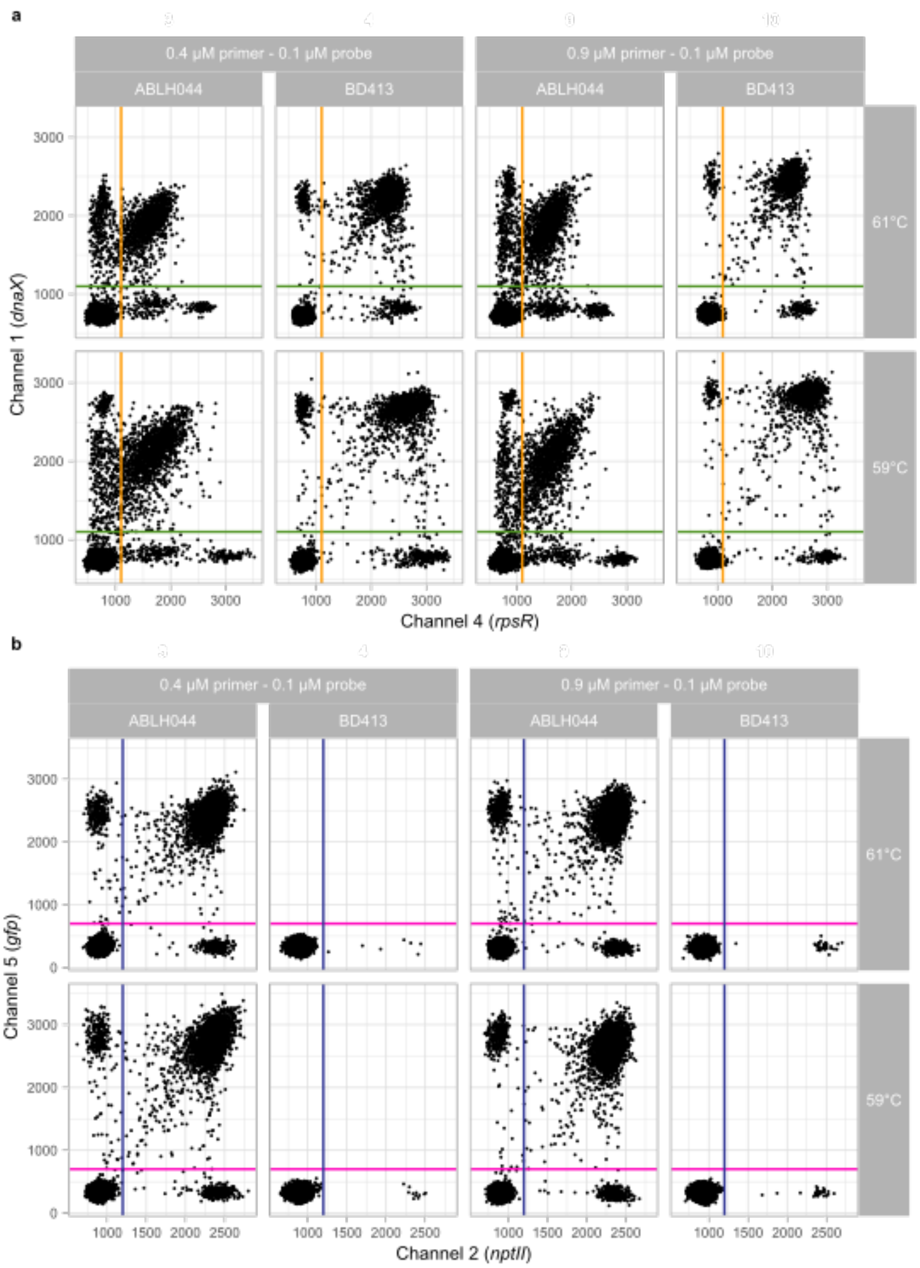


**Figure S3. Optimization of ddPCR conditions**
ddPCR test runs were performed with varying primer/probe concentrations and annealing temperatures using plasmid-bearing strain ABLH044 (plasmid copy number, PCN≈10) and plasmid-free BD413. Dots represent individual droplets. Subplots in (a) show chromosomal targets, and the corresponding samples in (b) show plasmid targets. Vertical and horizontal lines indicate fluorescence amplitude thresholds used to classify droplets as positive or negative. (a) In ABLH044, double-positive chromosomal droplets show reduced fluorescence amplitude compared to single-positive droplets, consistent with competition for amplification resources. This effect is not observed in BD413. Increasing annealing temperature reduces positive–negative cluster separation. (b) Plasmid target cluster separation remains robust across conditions. Lower annealing temperatures increase signal separation. Low-frequency false positives are detected for the novel allele but are minimized at lower primer/probe concentrations. Final assay conditions were 59°C annealing temperature with 0.4 μM primers and 0.1 μM probes.

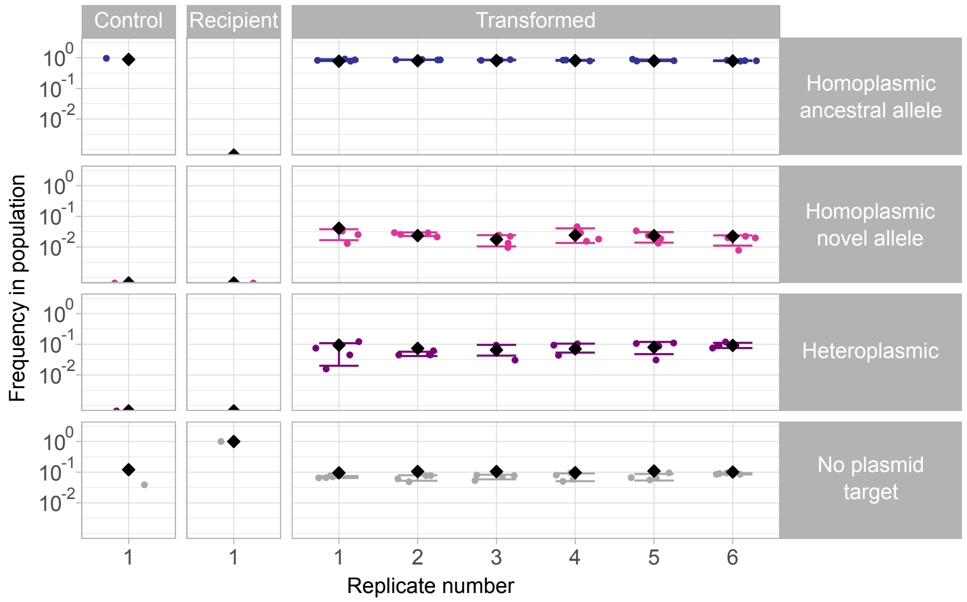


**Figure S4. Comparison of ddPCR results and phenotypic classification**
Cell-type frequencies in the pNCL population at day 1 determined by ddPCR (diamonds) and colony streak tests (dots; 95% confidence intervals). Estimates of of heteroplasmic and homoplasmic cell frequencies are consistent between the two methods, whereas plasmid-free cells are slightly overestimated by ddPCR.

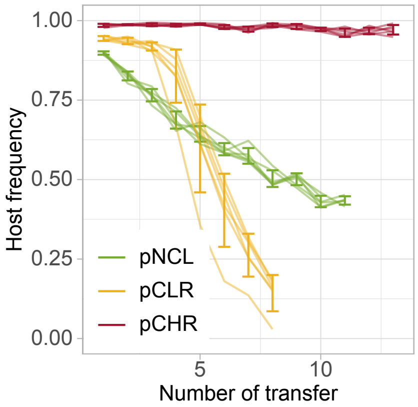


**Figure S5. Stability of model plasmids in donor culture throughout the evolution experiment**
Frequency of plasmid-hosts in the donor populations over the course of the evolution experiment. The frequency of pCLR and pNCL hosts declined over time, whereas pCHR is stably maintained within the host population. Error bars indicate SD.

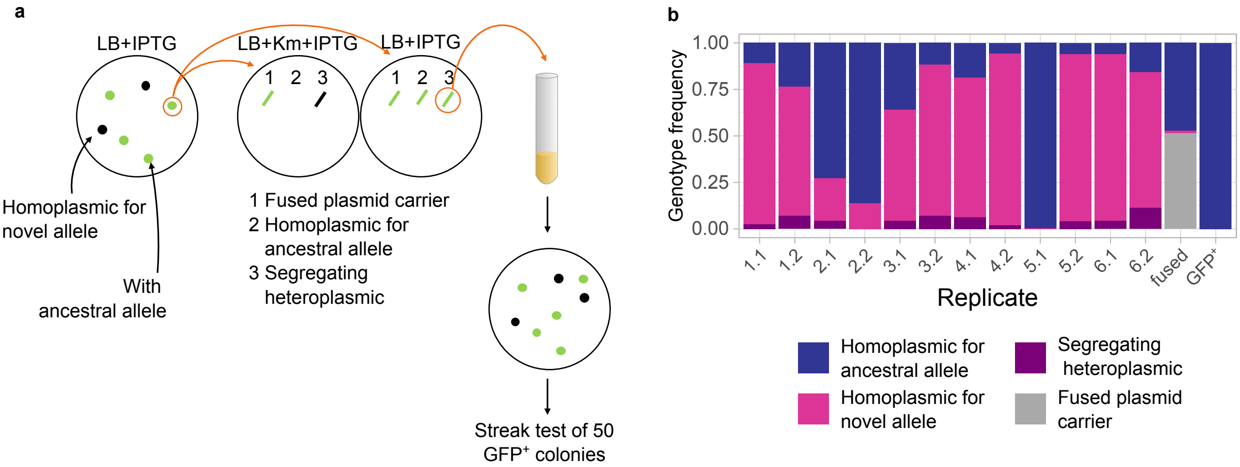


**Figure S6. Segregation of heteroplasmic, non-fused plasmids in pCLR donor cultures.** (a) Illustration of the workflow used to identify the subpopulation of heteroplasmic donor cells that carry non-fused plasmids and to assess their ability to segregate. Donor cultures were plated on non-selective LB plates supplemented with IPTG, to induce expression of the ancestral allele (*gfp*). GFP$^-$ colonies were classified as homoplasmic for the novel allele, whereas GFP$^+$ colonies carried the ancestral allele. GFP$^+$ colonies were subsequently streaked on LB agar supplemented with kanamycin (Km) and IPTG, as well as on LB + IPTG plates, to distinguish cells carrying fused plasmids, homoplasmic ancestral hosts, and heteroplasmic cells harboring both alleles on separate plasmid copies. Two heteroplasmic seeding cell streaks per replicate were analyzed further for their cell type distribution. (b) Cell type distribution of heteroplasmic seeding cells after streak tests (as described in a). Non-fused heteroplasmic cells occur at low frequency and segregate into homoplasmic cell types.

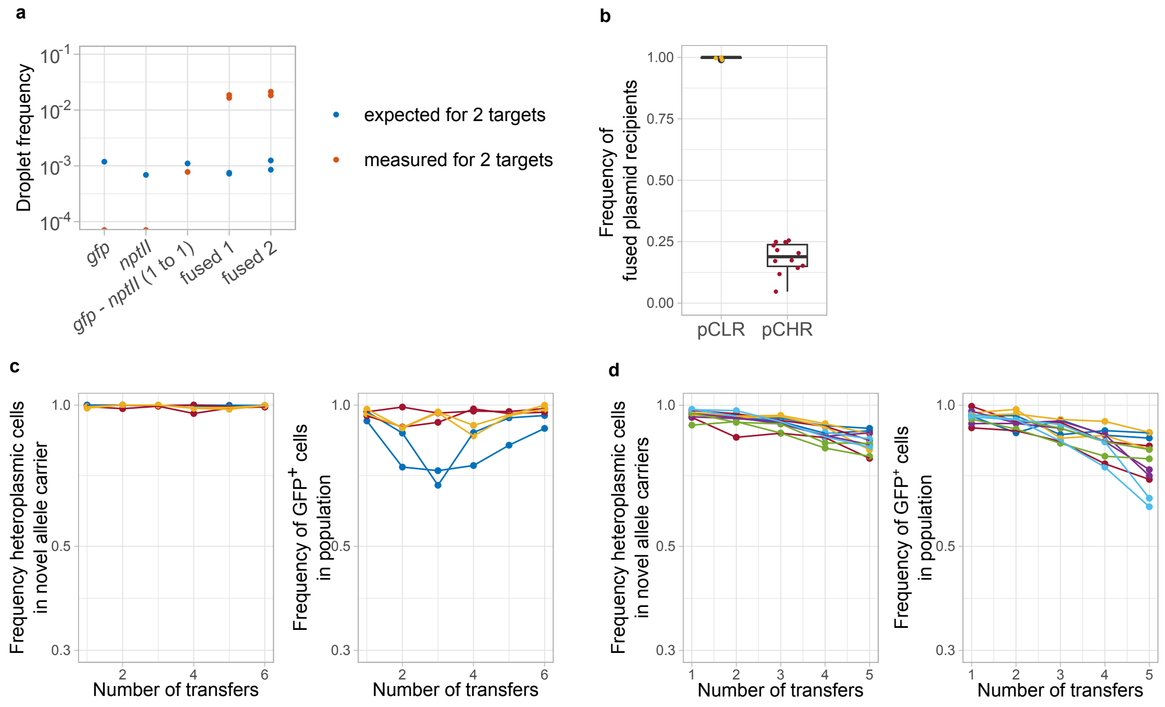


**Figure S7. Characterization of fused plasmid carriers in pCLR and pCHR donor populations.**

(a) Linkage ddPCR of purified pCLR plasmid variants: the ancestral-type plasmid (carrying *gfp*), the novel-type plasmid (carrying *nptII*), a 1:1 mixture of the ancestral- and the novel allele plasmids, and two candidates of fused plasmids (fused 1 and fused 2). Expected ratios of droplets containing two or more targets (blue) calculated from the input target amount and the total droplet count for every reaction. The ratios of droplets containing the novel and the ancestral allele per total droplet count were recorded per sample (orange). In samples carrying only the ancestral or only the novel plasmid variant, no double-target droplets were detected. In the 1:1 plasmid mixture, expected and measured double-target ratios are similar. In fused plasmid candidates, the measured frequency of double-target droplets exceeds the expected ratio by more than one order of magnitude, consistent with plasmid fusion. (b) Frequency of heteroplasmic plasmid harboring recipients following mating with fused plasmid donor clones from pCLR and pCHR donor cultures. The observed frequency of heteroplasmic recipients exceeds stochastic expectations, consistent with plasmid fusion in the donor. (c, d) Serial propagation of isolated fused-plasmid carriers from pCLR (c) and pCHR (d). Left: frequency of phenotypically heteroplasmic cells among novel-allele carriers (colony-forming units on selective plates). Right: frequency of GFP$^+$ cells in the total population. Limited segregation into homoplasmic novel allele cells is observed, whereas retention of the ancestral allele remains stable. Colors denote clones originating from the same replicate.

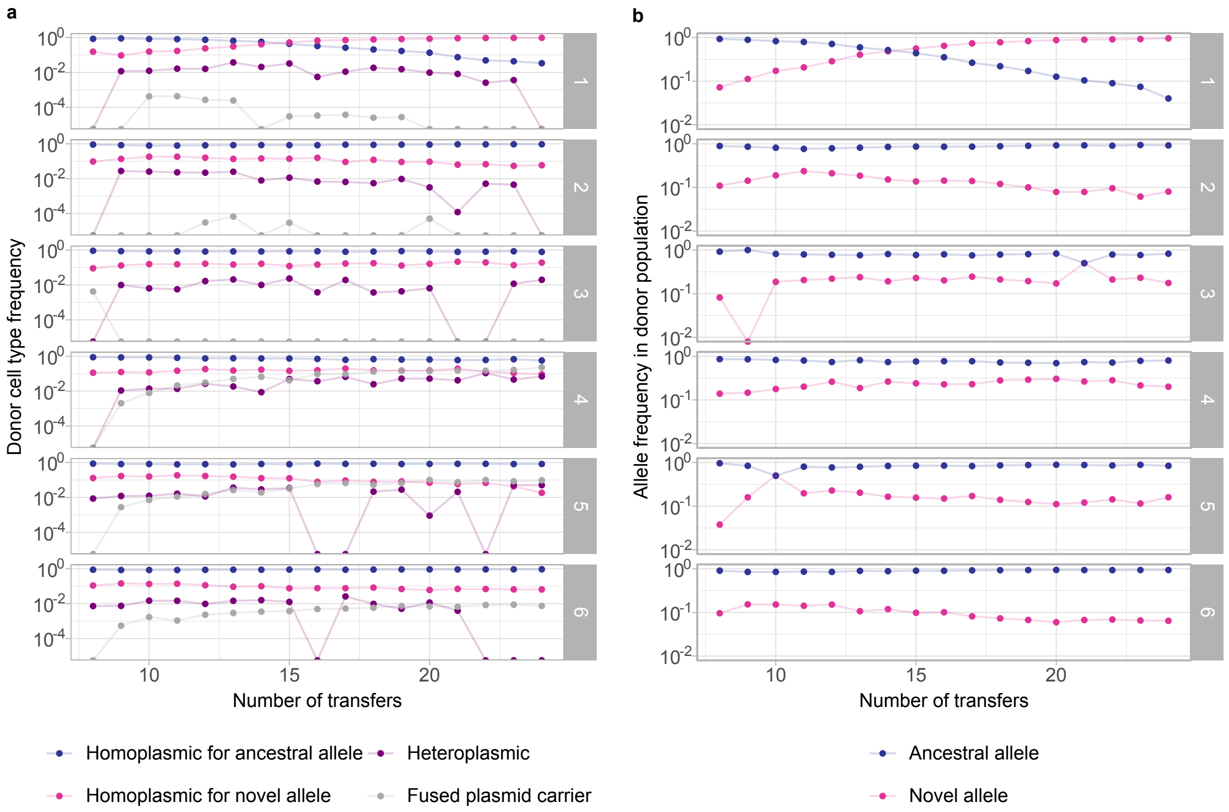


**Figure S8. Allele dynamics during extended experimental evolution of the pCLR donor populations.**

(a) Cell type frequencies among plasmid host cells in the continued serial pCLR donor culture (after selection for plasmid hosts) over time, shown for each replicate. In replicate 1, homoplasmic novel cells increase to frequencies exceeding those of homoplasmic ancestral cells. In most replicates, heteroplasmic cells persist, including cells harboring both alleles on two separate plasmids or on a fused plasmid. Fused-plasmid carriers increase in frequency in replicates 4-6 but are not detected at late time points in replicates 1-3, indicating that plasmid fusion is not required for maintenance of heteroplasmy. (b) Corresponding allele frequencies for the donor populations shown in (a). The novel allele frequency remains stable in most replicates but increases continuously in replicate 1.

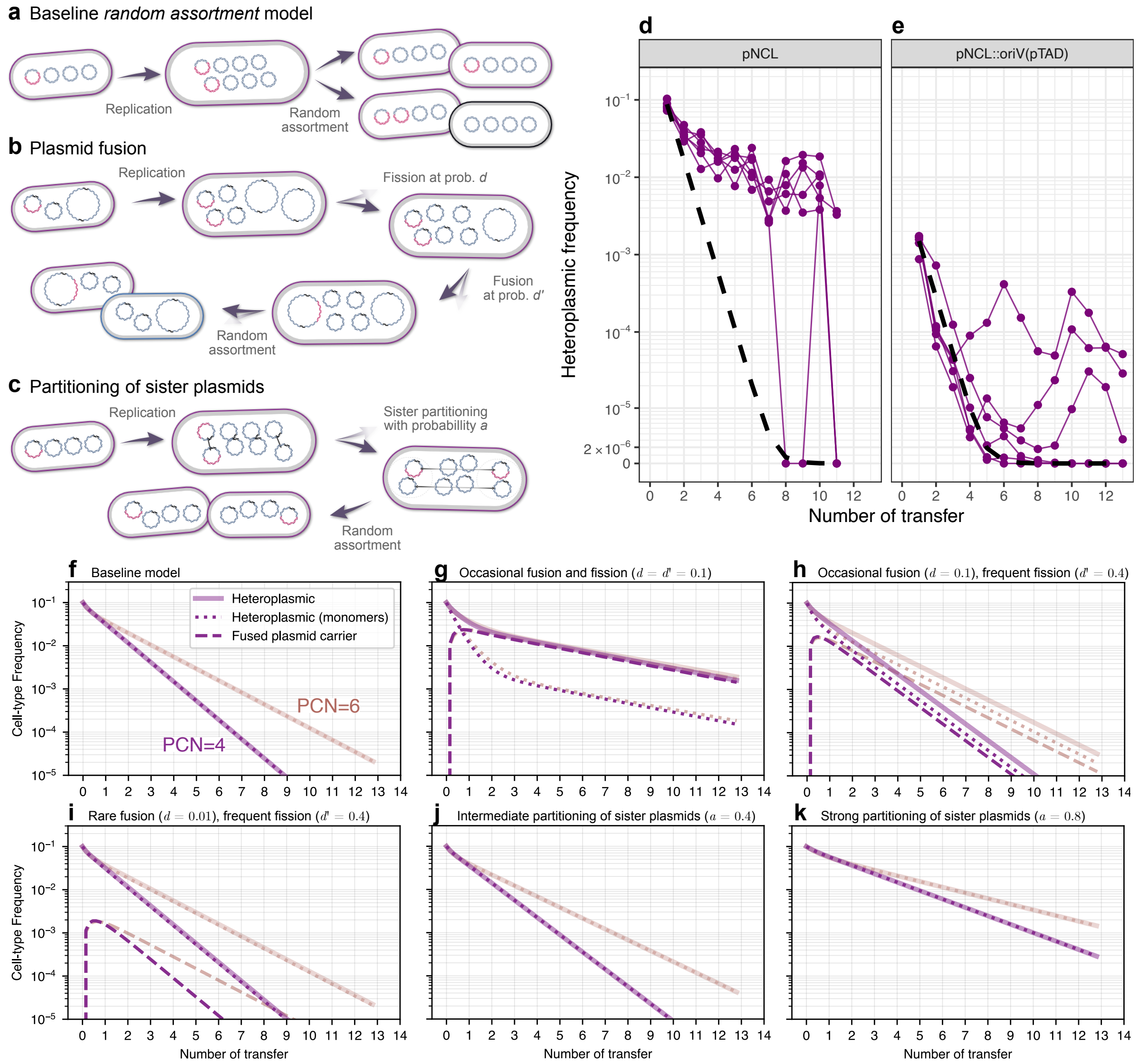


**Figure S9. Modeling and simulation of plasmid-allele segregation dynamics with plasmid fusion and sister-plasmid partitioning.**

Models of allele segregation for plasmids allowing plasmid fusion and fission or active separation of sister plasmids. Each plasmid copy is replicated prior to cell division. (a) Baseline model: plasmid copies are randomly assorted in equal numbers into the two daughter cells. (b) Plasmid fusion model: plasmid dimers resolve into monomers with probability $d'$; monomers subsequently pair with another plasmid copy and dimerize with probability $d$. (c) Sister-plasmid separation model: the two plasmids generated during replication are actively segregated into different daughter cells with probability $a$ (double-headed arrows); otherwise, the remaining plasmids are randomly assorted between daughter cells in equal number. (d, e) Comparison of experimental heteroplasmic frequencies for pNCL and pNCL::oriV(pTAD) with simulated heteroplasmic frequencies under the baseline model (Supplemental text, Eq. 5). Simulations were performed using the corresponding plasmid copy number ($n = 5$) and initial frequencies calibrated to the experimental starting values. The number of generations per transfer in the simulations, $\log_2(100)$, reflects population growth between two serial transfers with a bottleneck factor of 100. (f–k) Simulated heteroplasmic cell-type frequencies for plasmid copy numbers (PCN) $n = 4$ (violet) and $n = 6$ (light orange) under the three models with various parameters.

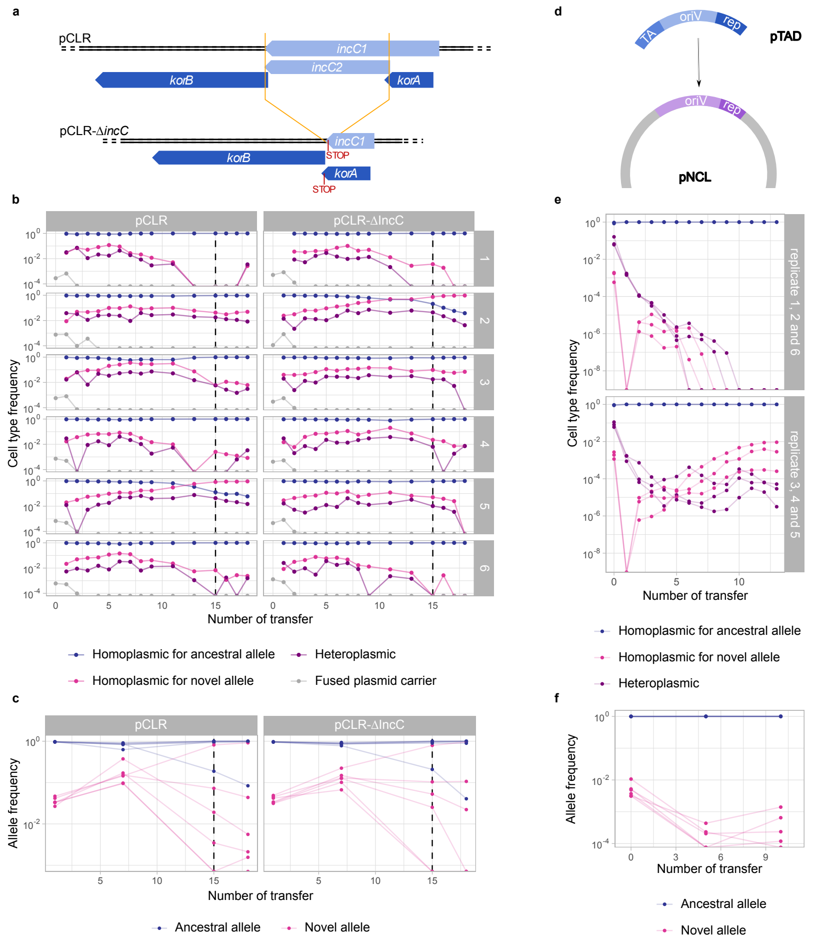


**Figure S10. Experimental evolution of pCLR-Δ*incC* and pNCL::*ori*$_{\text{pTAD}}$.**
(a) Genetic configuration of pCLR-Δ*incC*. The *incC* region not overlapping with *korA* was deleted; a stop codon was introduced in frame with the N-terminal part of *incC1* while maintaining the stop codon of *korA*. (b) Cell type frequencies among plasmid hosts during experimental evolution of pCLR and pCLR-Δ*incC* over time, shown per replicate. Dashed lines mark a shift from selection for the plasmid backbone to non-selective conditions. In both plasmids, one replicate shows an increase of novel homoplasmic cells above ancestral homoplasmic cells. Heteroplasmic cells persisted in most replicates, whereas fused plasmids were lost early in all replicates. (c) Corresponding allele frequencies in the donor populations shown in (b). Novel allele frequency increased during the first seven days in all replicates of pCLR and pCLR-Δ*incC* and subsequently diverged among replicates. (d) Genetic configuration of pNCL::oriV$_{\text{pTAD}}$. The native *oriV* and replication initiation protein (*rep*) of pNCL were replaced by the *oriV*, *rep*, and toxin-antitoxin (TA) system of pTAD. (e) Cell type frequencies among plasmid hosts during serial culture of pNCL::*ori*$_{\text{pTAD}}$, determined by differential plating; replicates are grouped by the pattern of allele dynamics. Heteroplasmic cells declined rapidly in all replicates, either falling below the detection limit or stabilizing at low frequencies. Novel homoplasmic cells arose in all replicates. In replicates 1, 2, and 6, novel homoplasmic cell frequencies remain below those of heteroplasmic cells, whereas in replicates 3, 4, and 5 they increased one to two orders of magnitude above heteroplasmic cell frequencies and remained stable until the end of the experiment. (f) Corresponding allele frequencies in (e). Novel allele frequency declines during the first five transfers and subsequently stabilizes in most replicates.

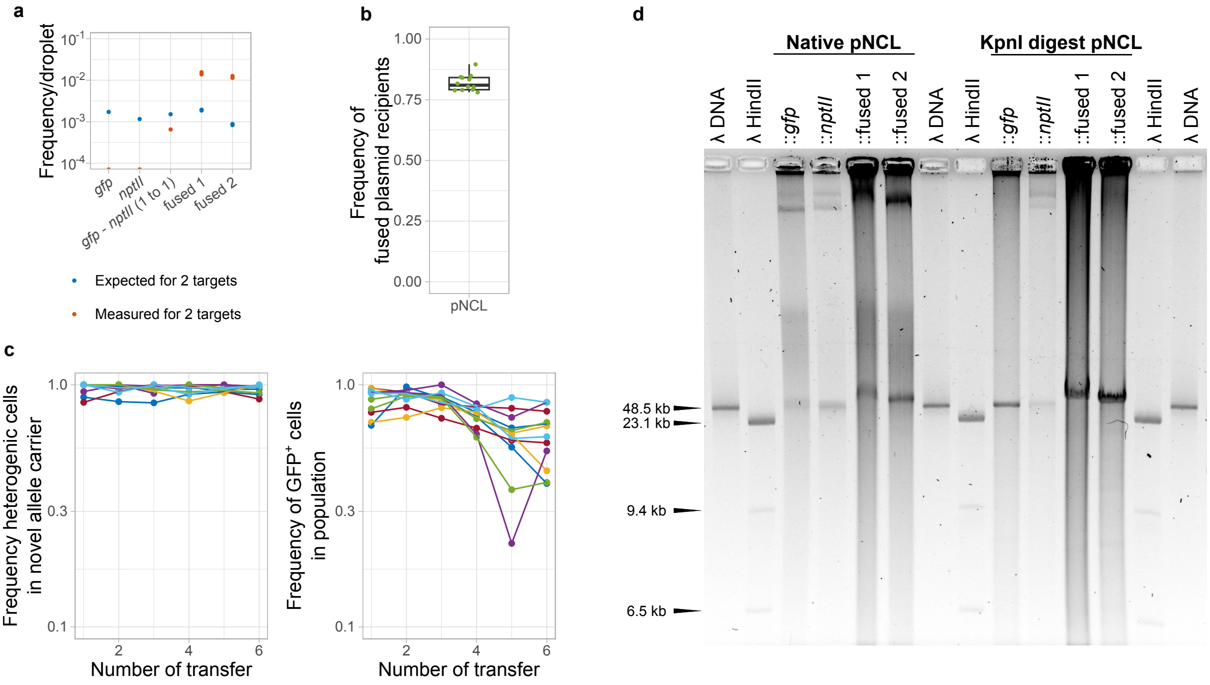


**Figure S11. Characterization of fused plasmid carriers in pNCL recipients.**

(a) Linkage ddPCR of purified plasmid variants isolated from pNCL recipients: ancestral (*gfp*), novel (*nptII*), a 1:1 mixture of ancestral and novel plasmids, and two fused plasmid candidates (fused 1 and fused 2). Expected ratios of double-target droplets (blue) were calculated from input target amounts and total droplet counts; measured ratios are shown relative to total droplets. No double-target droplets were detected in samples containing only ancestral or only novel plasmids, and the 1:1 mixture shows similar expected and measured ratios. Fused candidates exhibit double-target droplet frequencies more than one order of magnitude above expectation, consistent with plasmid fusion. (b) Frequency of heteroplasmic plasmid harboring recipients after mating of fused plasmid recipient clones from pNCL recipient cultures. Observed heteroplasmic recipients exceed stochastic expectations, consistent with plasmid fusion. (c) Serial propagation of isolated fused-plasmid recipients. Left: frequency of phenotypically heteroplasmic cells among novel-allele carriers. Right: frequency of $GFP^{+}$ cells in the total population. Limited segregation into homoplasmic novel cells was observed, whereas retention of the ancestral allele remained stable. Colors denote clones originating from the same experimental evolution replicate. (d) Agarose gel electrophoresis of plasmid preparations (native and *KpnI*-digested). *KpnI* cuts within the ancestral allele but not within the novel allele. Fused plasmids migrate above monomeric ancestral and novel plasmids in both untreated and digested samples, consistent with increased plasmid size. Fused 1 migrated above fused 2, indicating size differences between candidates. No additional bands are detected after digestion, consistent with homoplasmic fused-plasmid carriers containing a single ancestral allele copy. Gel resolution did not permit precise determination of plasmid size or multimeric state. Brightness and contrast were adjusted uniformly across the entire uncropped image.

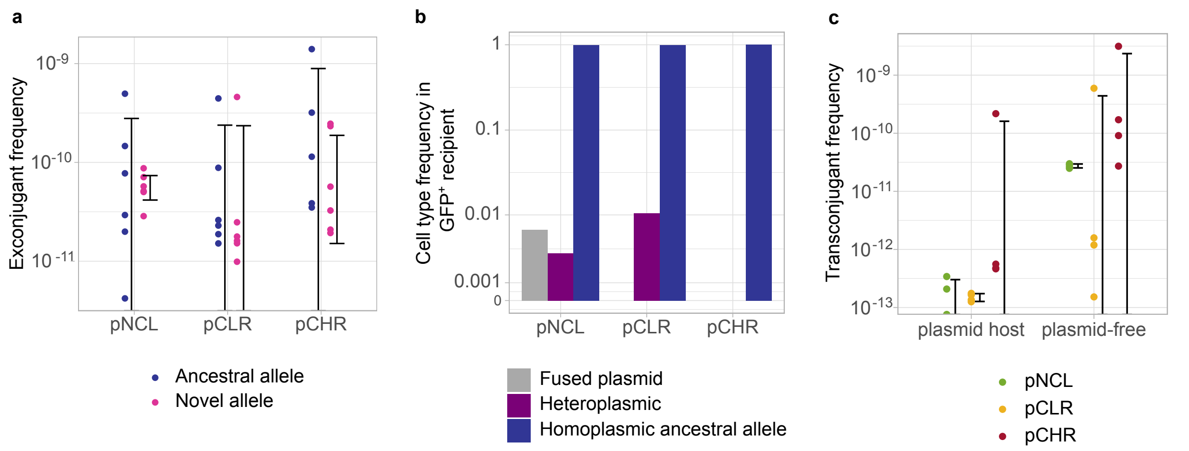


**Figure S12. Conjugative frequency of model plasmids.**
(a) Exconjugant frequency, T/(DR), following matings of ancestral homoplasmic (blue) or novel homoplasmic (pink) cells with plasmid-free recipients under conditions similar to the evolution experiment. Conjugation frequencies are similar between both alleles for each model plasmid. Error bars indicate 95% confidence intervals. (b) Phenotypic classification of plasmid harboring recipients at day 1 of the evolution experiment. In pNCL and pCLR recipient populations, segregating heteroplasmic recipients occured at low frequencies (0.4% and 1%, respectively). The pNCL recipient population additionally contained fused plasmids. No heteroplasmic recipients (fused or segregating) were detected in the pCHR population, indicating that transfer of multiple plasmids into a single recipient was rare. (c) Exconjugant frequency (T/(DR)) after matings of homoplasmic novel allele donors with ancestral allele recipients (plasmid hosts, left) compared to homoplasmic ancestral or novel allele donors (2 each) with with a plasmid-free recipient population. Conjugation frequencies to plasmid-free recipients was about two magnitudes higher than for the matings with plasmid harboring recipients. Error bars indicate 95%-CI.

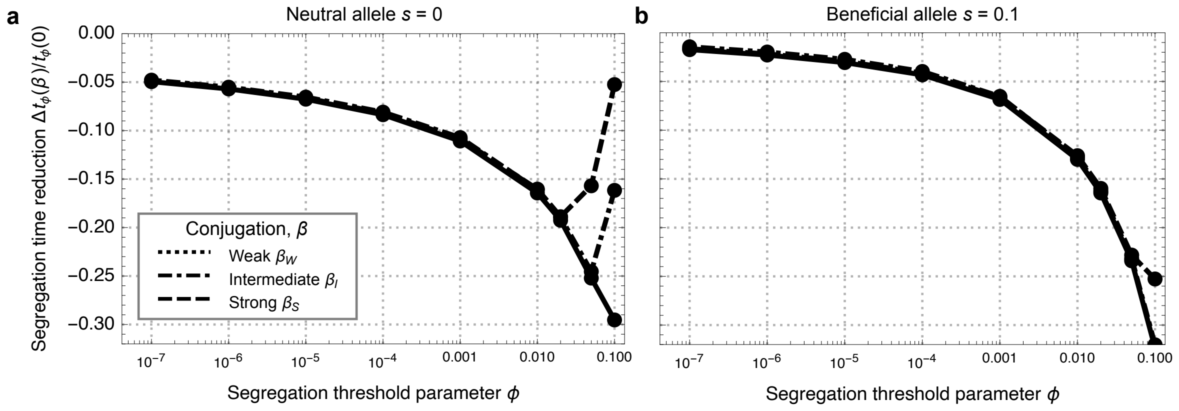


**Figure S13. Effect of conjugation on segregation time.** Relative reduction in segregation time due to conjugation, $\Delta t_{\phi}(\beta)/t_{\phi}(0)$, for different segregation threshold parameters $\phi$ for (a) neutral alleles ($s = 0$) and (b) beneficial alleles ($s = 0.1$). Parameter: plasmid copy number $n_{\mathrm{PCN}} = 5$.

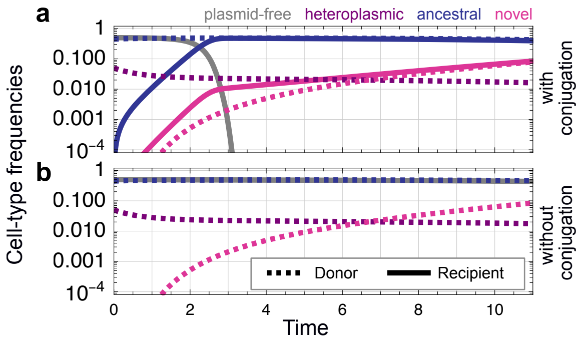


**Figure S14. Simulated segregation dynamics for beneficial plasmid alleles with selection parameter ($s = 0.1$)** (analogous to Fig. 4b)**.** Frequencies of plasmid-free, heteroplasmic, homoplasmic ancestral, homoplasmic novel cells (color code shown above the plot) under (a) plasmid conjugation with a high transfer-rate coefficient, $\beta_H = 10^{-10}$, and (b) no conjugation ($\beta = 0$) assuming plasmid copy number $n_{\mathrm{PCN}} = 10$.

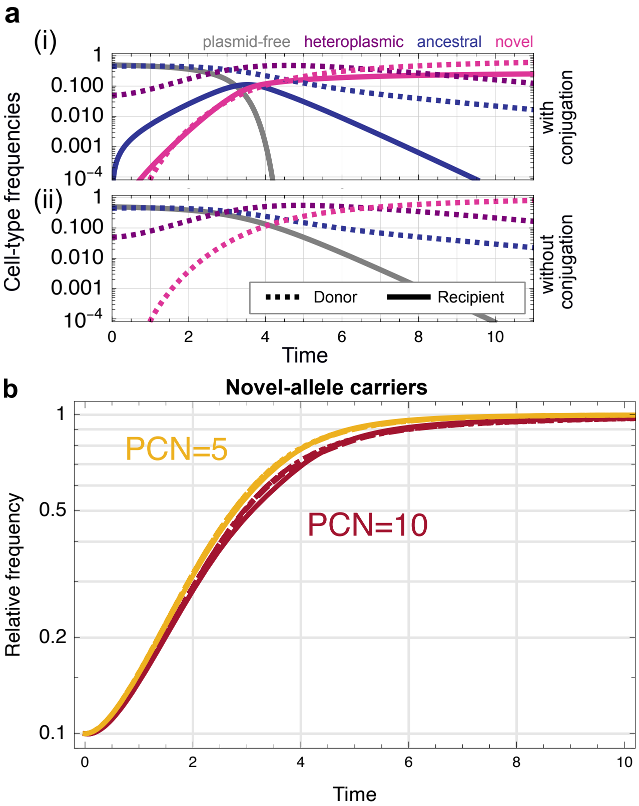


**Figure S15. Simulated segregation dynamics for beneficial plasmid alleles under strong selection ($s = 0.5$).** (a) Frequencies of plasmid-free, heteroplasmic, ancestral-homoplasmic, novel-homoplasmic cell types (color code shown above) under (i) plasmid conjugation with a high transfer-rate coefficient ($\beta = \beta_H = 10^{-10}$, and (ii) without conjugation ($\beta = 0$), assuming for plasmid copy number $n_{\mathrm{PCN}} = 10$. (b) Relative frequency of novel-allele carriers, defined as the proportion of plasmid-host cells (donors and transconjugants) carrying at least one plasmid copy encoding the novel allele.

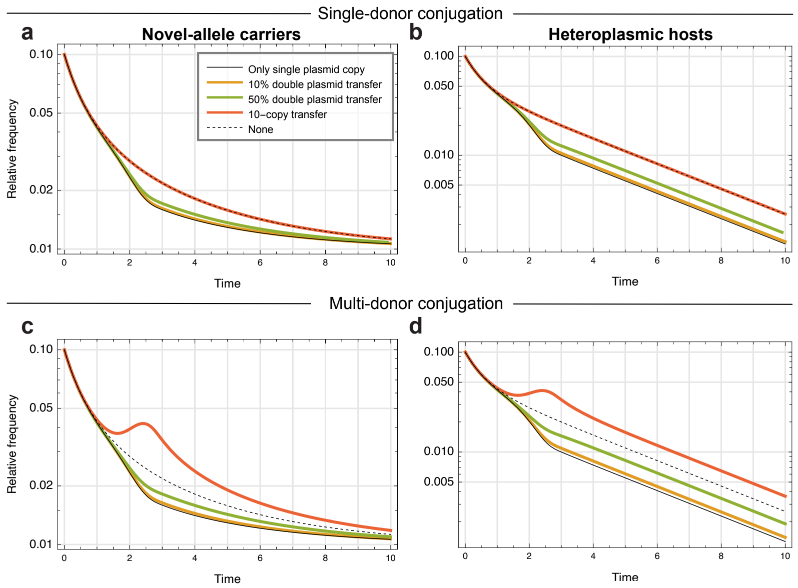


**Figure S16. Multi-plasmid transfer from a single donor.** Simulated plasmid-allele segregation dynamics for multi-copy transfer with single-donor conjugation (a, b) and multi-donor conjugation (c, d). Relative frequency of novel-allele carriers (left plots) and heteroplasmic hosts (right plots) under multi-copy transfer (bold colored lines), single-copy transfer (thin solid line), and no conjugation (dotted line). For multi-copy transfer, simulations include double-plasmid transfer with probabilities of 10% and 50% (corresponding to 90% and 50% single-copy transfer, respectively) and the transfer of all $n = 10$ plasmid copies with probability 1. Parameter: plasmid copy number $n = 10$; transfer-rate coefficient, $\beta = \beta_H = 10^{-10}$ (high).

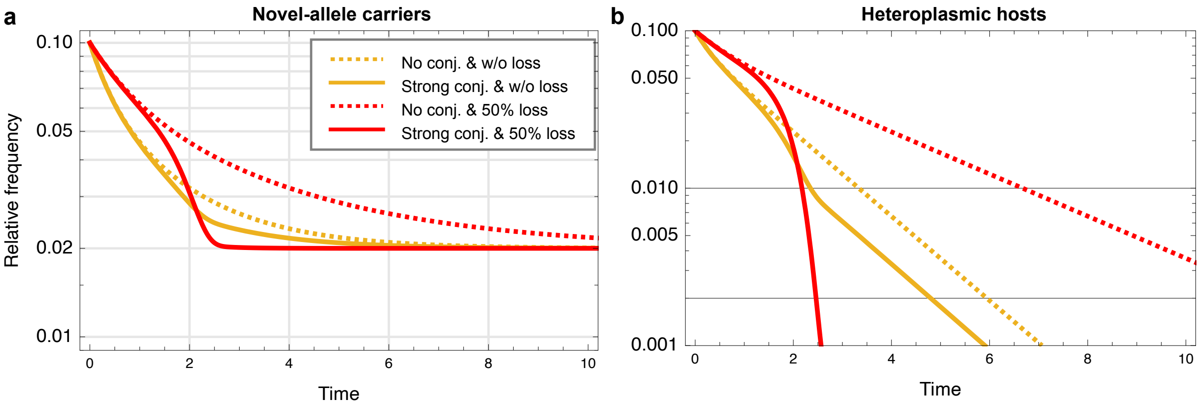


**Figure S17. Plasmid-allele segregation dynamics under plasmid loss.** Comparison of faithful plasmid inheritance ($p = 1$) and unfaithful plasmid inheritance ($p = 0.5$). (a) Relative frequency of novel-allele carriers, defined as the proportion of plasmid-host cells (donors and transconjugants) carrying at least one plasmid copy encoding the novel allele. (b) Relative frequency of heteroplasmic hosts, defined as the proportion of plasmid-host cells carrying both allelic variants. Plasmid copy number $n = 10$.

**Supplementary Tables**
**Table S1. *Acinetobacter baylyi* strains used in this study**
Strain number, strain name, and phenotypic marker are indicated.

| number | Strain name | Phenotypic marker |
|---|---|---|
| NH24 | BD4 Δ*dprA* | GmR |
| ABLH007 | BD4 | prototrophic |
| ABLH008 | BD413 | tryptophan auxotrophic |
| ABLH017 | BD413 pCLR | SpR, $GFP^+$ on IPTG |
| ABLH027 | BD4 Δ*comB-F* | TmR |
| ABLH030 | BD4 Δ*comB-F* pNCL | TmR, SpR |
| ABLH031 | BD4 Δ*comB-F* pCLR | TmR, SpR |
| ABLH032 | BD413 pNCL | SpR, $GFP^+$ on IPTG |
| ABLH044 | BD413 pCHR-[*sacB-nptII*] | SpR, $GFP^+$ on IPTG, KmR, SucS |
| ABLH048 | BD413 pCHR | SpR, $GFP^+$ on IPTG |
| ABLH053 -ABLH056 | BD413 pNCL::*nptII* | SpR, KmR |
| ABLH057-ABLH060 | BD413 pCLR::*nptII* | SpR, KmR |
| ABLH061-ABLH064 | BD413 pCHR::*nptII* | SpR, KmR |
| ABLH049 | BD413 pNCL::$oriV_{pTAD}$ | SpR, $GFP^+$ on IPTG |
| ABLH071 | BD413 pCLR-Δ*incC* | SpR, KmR, $GFP^+$ on IPTG |
| ABLH077 | BD4 Δ*comB-F* pCHR | TmR, SpR, $GFP^+$ on IPTG |

**Table S2. Cloning and validation primers used in this study**
Primer name, sequence, and annealing temperature ($T_M$) are listed. Overhangs are indicated by underline.

| Number | Sequence (5'-3') |
|---|---|
| CP1 | CCTCACTGATTAAGCACTGGTAACGGTGAAATCATTGTCAATTAAGGC |
| CP2 | CTTTGTCGTATCCACGTTCTAGAGCTTATCATGTCAGCCTATTTGC |
| CP3 | GCCTTCTTGACGAGTTCTTCTAGACCTGATTAGGCATTCTGTACCG |
| CP4 | AAAAGCTGGAGATCAAGTTCTTTAGCAGCAGTAGTTTCTATTCAACC |
| NH138 | TCGAACCCCAGAGTCCCGC |
| NH139 | GTAGCCCAGCGCGTCGCTTGC |
| NH226 | GCGCGTGGTATGGGAGCCTAGGGCTTATTTGCCGACTACCTTGGTG |
| NH227 | ACGCCGGTAGGTAATTGCTAGCGGATCGTAAGCTGTAATGCAAGTAGCG |
| NH286 | TGTAATGCTGTTGTATTTTGCCCAG |
| NH287 | GCCCTAGGCTCCCATACCACGCGCTCGCTCTATTCCCTTGAGGTTG |
| NH290 | CCGCTAGCAATTACCTACCGGCGTAAACAGGACATTTTAAATGGTTTATCG |
| NH291 | CTTACTAACAGCTTTATCTTTTGCC |
| NH293 | TCCAGACGATGTGGTTGAGC |
| oLH009 | CCCTGATCGAGGTCGAGGGGGCATCGCACGCCGGTAGGTAATTGCTAG |
| oLH010 | GCCAAACCGTCCCTCCGAGTGGTTCGAAGCACGTGCGCGTGGTATGGGAG |
| oLH011 | CTGCATCGAAATTGGCCTTGC |
| oLH012 | ACGTGCTTCGAACCACTCG |
| oLH013 | GCGATGCCCCCTCGAC |
| oLH014 | CCGACGGCAAGCAATGG |
| oLH023 | GGTTGTGCTGTGGCTCG |
| oLH034 | AGGGGTGACGCCAAAGTATAC |
| oLH035 | TGCTTTACGGTATCGCCGC |
| oLH036 | ACGTGGATACGACAAAGAGTCC |
| oLH038 | CGGTATCATTGCAGCACTGG |
| oLH040 | TAACTCGTCAAGAAGGCGATAGAAGG |
| oLH041 | GGCAAACCTAGCCAACG |
| oLH042 | GCGTTCGTTACCAGTGCTTATATTCAGCCTATTTGCAACAGTGCCAG |
| oLH043 | GGCACTGTTGCAAATAGGCTGAATATAAGCACTGGTAACGAACGC |
| oLH044 | TATGTCGAACGATGTGGACCTATTGAAATCGAACCCCAGAGTCCC |

| Number | Sequence (5’-3’) |
|---|---|
| oLH045 | CTGAGAGCGGGACTCTGGGGTTCGATTTCAATAGGTCCACATCGTTCGAC |
| oLH046 | CCTCAACTTTGGAATCGTTTGG |
| oLH047 | TGTTGCCCGTCTCACTGG |
| oLH048 | CGCTTCCTCGTGCTTTACG |
| oLH049 | ACGACTTACTGCATCCAGGAC |
| oLH052 | GGTTTCTGCCGACGTACTTCG |
| oLH068 | GCTGGCACGACAGGTTTCC |
| oLH079 | ACCCTCACTAAAGGGAACAAAAGCTGGAGACATTGCACTCGATTCCAGAC |
| oLH092 | CGAGGATCGCTTTCACTGG |
| oLH093 | AGTCATTCTGCCCCGACC |
| oLH116 | AGAGCGGCAGAGATGAACACG |
| oLH117 | CTGCTGCACTGCTTCCGCGTC |
| oLH118 | ACGCGGAAGCAGTGCAGCAGC |
| oLH123 | CTCACTAAAGGGAACAAAAGCTGGAGATCGACGGCACAGGCTACATCC |
| oLH124 | CTTTGTCGTATCCACGTTCTAGAAGAGCGGCAGAGATGAACACG |
| oLH126 | CTCCAGCTTTTGTTCCCTTTAGTG |
| oLH127 | TGCTTAATCAGTGAGGCACCTATC |
| oLH128 | CCACGCGCAACAAGAAAATCC |
| oLH129 | TGTCGGCCCTGAAGAAAGC |
| oLH138 | CATTAAGGTTTGACGACCGCTAATGATTTCACCCGAATGGCTCGCGTTGG |
| oLH140 | CTGACGACACGCAAACTGG |
| oLH142 | TCGCCCAGGCAATGATCACC |
| oLH143 | GACTTCATTGCCGATAAGGTGG |
| oLH144 | CACATCGCAGGAGTGGTCATG |
| oLH146 | TGGGTCTTAAAACGCAAATGGTG |
| oLH147 | ACGCAAGTTTTAGCTATGGTGC |
| oLH148 | CCACCGGTGGAGATCGAAC |
| oLH149 | GCCCCAAGCAATTGCAAGG |
| oLH150 | GCGTGCTAGACGAATAACAACC |
| oLH151 | GTATGCGAGCTAATCCTCAAGC |
| oLH153 | GCCTGACCTTGAGCAGTCG |
| oLH154 | ATAGGTGCCTCACTGATTAAGCACTGACTGCTCCCATTTGATTTAGTAGC |
| oLH158 | GCTGGACTCTTTGTCGTATCCACGTTCTAGACTGCGCTCGGATCTTGATG |
| oLH159 | TGGCATTGCTGCACATTGAATAAATCTACCACGTTATGACCAAGAACAGC |

| Number | Sequence (5'-3') |
| --- | --- |
| oLH160 | AGGCTGTTCTTGGTCATAACGTGGTAGATTTATTCAATGTGCAGCAATGC |
| oLH165 | TCATCTTGCGTCCTTTAAGCTC |
| oLH166 | CTTCTTGACGAGTTCTTCTAGTCTAGAGGTAGCCCGATACGATTGATGG |
| oLH167 | GAGATAGGTGCCTCACTGATTAAGCACTGGTGAGGCAATCCCGCAAGGAG |
| oLH171 | GGTGCCTCACTGATTAAGCACTGGTGACTGCTCCCATTTGATTTAGTAGC |
| oLH172 | CTTCTTGACGAGTTCTTCTAGTCTAGACGATCAGGACACAGACTATCAGG |
| oLH175 | GCTTGGATAAAAGCTGCAGCG |
| oLH178 | AGCTCCTGGTCTACGTTCG |
| oLH179 | GTATAAATCGGCTTGTTCCACG |
| oLH180 | ATACGCTGGACTCTTTGTCGTATCCACGTCTAGACGTTTAGTTTCCTGTTTTTTCTTGGC |
| oLH181 | ATAGGTGCCTCACTGATTAAGCACTGGTGAGTGTTGACGTGCGAGAAATG |
| oLH182 | ATCGCCTTCTTGACGAGTTCTTCTAGTCTAGAATGACTGCGGCTCAAGCC |
| oLH183 | CTAAAGGGAACAAAAGCTGGAGATCAAGTTGTAGTCGTTGTCGATGAACG |
| oLH184 | GACCTGTTGGTGATGGAGC |
| oLH185 | GCCTTTCTTCTTGCCCTTCG |
| oLH186 | GGCAACCGGCTTTGTTGG |
| oLH187 | GGTCTTGGCTTGAGCCGCAGTCATCGTTTAGTTTCCTGTTTTTTCTTGGC |
| oLH188 | GCGGACGCCAAGAAAAAACAGGAAACTAAACGATGACTGCGGCTCAAGCC |
| oLH189 | CCTTCTATCGCCTTCTTGACGAGTTCTTCTAGAAGGGAGGCGTCTTGAGCA |

**Table S3. Strain construction details generated in this study**

The table summarizes the genetic modifications, plasmids, primers, and intermediate strains involved in the construction of each strain used in this work. Detailed explanations of all column headings are provided below.

| construct | descriptive name and description | parental | organism | DNA delivery | DNA | flanking region 1 | | | selection cassette | | | flanking region 2 A | | | flanking region 2 B | | | backbone | | | Valdiation Primer Pair 1 | | Validation Primer | | Validation Primer | | sequencing primer | | | | | | | | |
|---|---|---|---|---|---|---|---|---|---|---|---|---|---|---|---|---|---|---|---|---|---|---|---|---|---|---|---|---|---|---|---|---|---|---|---|
| | | | | | | template | primer 1 | primer 2 | template | primer5 | primer6 | template | primer 3 | primer 4 | template | primer 7 | primer 8 | template | primer 9 | primer 10 | primer A | primer B | primer C | primer D | primer E | primer F | for primerA-primerB product | | | for primerC-primerD product | | for primerE-primerF product | | for isolated plasmid | |
| BD4 ΔcomBF | BD4 comBF::dhfr1<br>allelic replacement of comBF operon with dhfr1 | BD4 | A. baylyi BD4 | natural transformation | KOM130 gDNA | | | | | | | | | | | | | | | | oLH151 | oLH153 | | | | | oLH150 | oLH154 | | | | | | | |
| BD4 ΔdprA | BD4 dprA::aacC1<br>allelic replacement of dprA gene with aacC1 | BD4 | A. baylyi BD4 | natural transformation | SOEing PCR product | | | | | | | | | | | | | | | | | | | | | | | | | | | | | | |
| pLH13 | vector for allelic replacement of AD-gfp with AD-nptII during evolution experiment by amplification of flanking regions and AD-nptII of pCLR::nptII | pJET | E. coli DH5alpha | chemical transformation | ligation product after employing pJET protocol | pCLR::nptII | oLH178 | oLH140 | | | | | | | | | | | | | | | | | | | | | | | | | | whole plasmid sequencing | |
| pLH14 | vector for allelic replacement of AD-gfp with AD-nptII during evolution experiment by amplification of flanking regions and AD-nptII of pNCL::nptII | pJET | E. coli DH5alpha | chemical transformation | ligation product after employing pJET protocol | pTF4-AD-nptII | oLH179 | NH293 | | | | | | | | | | | | | | | | | | | | | | | | | | whole plasmid sequencing | |
| pGT42-junk-sub | vector for replacement of the region 31,983–53,830 (21.8 kb) with [sacB nptII]-cassette in pHHV216 | pGT42 | E. coli DH5alpha | chemical transformation | Gibson Assembly | pHHV216 | CP1 | CP2 | [sacB nptII] cassette (created when pGT42 is cut with AleI & HpaI) | | | pHHV216 | CP3 | CP4 | | | | pGT42 (AleI & HpaI digested vector fragment) | | | | | | | | | | | | | | | | | |
| pTF2 | pHHV216 with region 31,983–53,830 replaced by [sacB nptII]-cassette | pHHV216 | A. baylyi BD413 | natural transformation | pGT42-junk-sub | | | | | | | | | | | | | | | | | | | | | | | | | | | | | | |
| pTF3 | pTF2 with insertion of aadA replacing the gene int (V216_47) | pTF2 | A. baylyi BD413 | natural transformation | SOEing PCR product | pHHV216 | NH286 | NH287 | pTKRED (GenBank: GU327533.1) | NH226 | NH227 | pHHV216 | NH290 | NH291 | | | | | | | | | | | | | | | | | | | | | |
| pGT42-junk-del | vector for deletion of the [sacB nptII]-cassette in pTF3 | pGT42 | E. coli DH5alpha | chemical transformation | ligation product (recircularization of XbaI-digested pGT42-junk-sub vector fragmenmt (i.e., devoid [sacB nptII]) | | | | | | | | | | | | | | | | | | | | | | | | | | | | | | |
| pTF4 | | pTF3 | A. baylyi BD4 | natural transformation | pGT42-junk-del | | | | | | | | | | | | | | | | | | | | | | | | | | | | | | |
| pNCL::nptII | pTF4-AD-nptII<br>insertion of lacIadi and a funtional nptII upstream of rumA | pTF4 | A. baylyi BD413 | natural transformation | SOEing PCR product | pTF4 | oLH041 | oLH042 | pTAD-R-nptII | oLH043 | oLH044 | pTF4 | oLH045 | oLH046 | | | | | | | oLH047 | oLH049 | oLH048 | oLH052 | | | oLH047 | oLH023 | | oLH046 | oLH048 | | | | |
| pNCL | pTF4-AD-gfp<br>allelic replacement of functional nptII with funtional gfp under control of Plac flanked by two nptII fragments (with a 10 bp deletion) | pNCL::nptII | A. baylyi BD413 | natural transformation | PCR product | | | | pTAD-R | NH138 | NH139 | | | | | | | | | | oLH138 | oLH189 | | | | | oLH040 | | | | | | | | |
| pLH07 | pGT42-pNCL::oriVpTAD-[sacB-nptII]<br>template vector for replacement of oriV-rep of pNCL with TAsystem-OriV-rep of pTAD with flanking regions of pNCL and a selection/counter-selection cassette sacB-nptII | pGT42 | E. coli DH5alpha | chemical transformation | Gibson Assembly | pNCL | oLH172 | oLH079 | pGT42 | oLH040 | oLH036 | AG19 | oLH158 | oLH159 | pNCL | oLH171 | oLH160 | pGT42 | oLH126 | oLH127 | oLH038 | oLH149 | oLH068 | oLH165 | | | | | | | | | | oLH034 | oLH048 |
| pNCL::oriVpTAD-[sacB-nptII] | allelic replacement of oriV-rep of pNCL with TAsystem-OriV-rep of pTAD and a selection/counter-selection cassette sacB-nptII | pNCL | A. baylyi BD413 | natural transformation | pLH07 | | | | | | | | | | | | | | | | oLH144 | oLH034 | oLH148 | oLH093 | oLH175 | oLH149 | oLH144 | oLH147 | oLH034 | oLH048 | oLH165 | oLH175 | oLH149 | | |
| pLH12 | pGT42-pNCL::oriVpTAD-Δ[sacB-nptII]<br>template vector for replacement of oriV-rep of pNCL with TAsystem-OriV-rep of pTAD | pLH07 | E. coli DH5alpha | chemical transformation | ligation product (recircularization of XbaI-digested pLH07 vector fragmenmt (i.e., devoid [sacB nptII]) | | | | | | | | | | | | | | | | oLH068 | oLH147 | | | | | | | | | | | | whole plasmid sequencing | |
| pNCL::oriVpTAD | deletion of selection/counter-selection casette sacB-nptII for markerfree exchange of pNCL OriV-rep with TAsystem-OriV-rep of pTAD by allelic replacement | pNCL::oriVpTAD-[sacB-nptII] | A. baylyi BD413 | natural transformation | pLH12 | | | | | | | | | | | | | | | | oLH175 | oLH146 | oLH147 | oLH093 | | | oLH175 | oLH146 | | oLH147 | oLH093 | | | | |
| RK2-aphA::aadA | deletion of aphA (KmR) and IS21 by allelic replacement with aadA (SpR) | RK2 | A. baylyi BD413 | natural transformation | SOEing PCR product | RK2 | oLH011 | oLH012 | aadA | oLH009 | oLH010 | RK2 | oLH013 | oLH14 | | | | | | | | | | | | | | | | | | | | | |
| pCLR::nptII | RK2-aphA::aadA-AD-nptII<br>deletion of Tn1 by allelic replacement with lacIadi and a funtional nptII | RK2-aphA::aadA | A. baylyi BD413 | natural transformation | SOEing PCR product | | | | pTAD-R-nptII | | | | | | | | | | | | oLH035 | oLJ052 | | | | | oLH035 | | | | | | | | |
| pCLR | RK2-aphA::aadA-AD-gfp<br>allelic replacement of functional nptII with funtional gfp under control of Plac flanked by two nptII fragments (with a 10 bp deletion) | pCLR::nptII | A. baylyi BD413 | natural transformation | PCR product | | | | pTAD-R | NH138 | NH139 | | | | | | | | | | oLH138 | oLH189 | | | | | oLH138 | | | | | | | | |
| pLH02 | pGT42-pCHR(trf271)-[sacB-nptII]<br>template vector for replacement of trfA(WT) with trfA(271C) with flanking regions of pCLR, an inserted point mutation at position 271aa from Arg to Cys (by primer permutation) and a selection/counter-selection cassette sacB-nptII | pGT42 | E. coli DH5alpha | chemical transformation | Gibson Assembly | pCLR | oLH166 | oLH123 | pGT42 | oLH040 | oLH036 | pCLR | oLH167 | oLH118 | pCLR | oLH124 | oLH117 | pGT42 | oLH126 | oLH127 | oLH034 | oLH129 | oLH035 | oLH068 | | | | | | | | | | oLH034 | oLH048 |
| pCHR-[sacB-nptII] | allelic replacement of trfA(WT) with trfA(271C),an inserted point mutation at position 271aa from Arg to Cys, and a selection/counter-selection cassette sacB-nptII | pCLR | A. baylyi BD413 | natural transformation | pGT42-pCHR-[sacB-nptII] plasmid prep | | | | | | | | | | | | | | | | oLH092 | oLH148 | oLH034 | oLH143 | | | oLH148 | oLH092 | oLH128 | | | | | | |
| pLH10 | pGT42-pCHR-Δ[sacB-nptII]<br>template vector for replacement of trfA(WT) with trfA(271C) with flanking regions of pCLR and an inserted point mutation at position 271aa from Arg to Cys (by primer permutation) | pLH02 | E. coli DH5alpha | chemical transformation | pGT42-pCHR-[sacB-nptII] ligation after XbaI digest | | | | | | | | | | | | | | | | oLH068 | oLH129 | | | | | | | | | | | | whole plasmid sequencing | |
| pCHR | deletion of selection/counter-selection casette sacB-nptII for markerfree exchange ofof trfA(WT) with trfA(271C),an inserted point mutation at position 271aa from Arg to Cys by allelic replacement | pCHR-[sacB-nptII] | A. baylyi BD413 | natural transformation | p LH10 pGT42-pCHR-Δ[sacB-nptII] plasmid prep | | | | | | | | | | | | | | | | oLH142 | oLH128 | oLH143 | oLH116 | | | oLH142 | oLH128 | | oLH143 | pLH116 | | | | |
| pCLR-ΔIncC-[sacB-nptII] | deletion of incC from aa position 104 to the end with an STOP insertion at aa104 without altering the aa sequence of overlapping korA by allelic replacement with selection/counter-selection cassette sacB-nptII | pCLR | A. baylyi BD413 | natural transformation | SOEing PCR product | pCLR | oLH180 | oLH181 | pGT42 | oLH040 | oLH036 | pCLR | pLH182 | pLH183 | | | | | | | oLH186 | oLH034 | oLH185 | oLH48 | | | oLH035 | oLH184 | | oLH35 | oLH184 | | | | |
| pLH14 | pGT42-pCLR-ΔIncC<br>template vector for scarless deletion of incC from aa104 on | pGT42 | E. coli DH5alpha | chemical transformation | Gibson Assembly | pCLR | oLH187 | oLH181 | | | | pCLR | pLH188 | pLH183 | | | | pGT42 | oLH126 | oLH127 | oLH068 | oLH038 | | | | | | | | | | | | oLH038 | oLH068 |
| pCLR-ΔIncC | scarless deletion of incC from aa104 on by alleleic replacement of selection/counter-selection cassette sacB-nptII | pCLR-ΔIncC-[sacB-nptII] | A. baylyi BD413 | natural transformation | pLH14 | | | | | | | | | | | | | | | | oLH186 | oLH185 | | | | | oLH184 | | | | | | | | |

### Description of column headings in Table S3

| column | description |
|---|---|
| **construct** | the strain or plasmid that is to be constructed |
| **descriptive name and description** | the name of the construct refering back to the previous construct; denoting the changes and the description of the changes itself |
| **parental strain/plasmid** | the strain or plasmid the changes were introduced in |
| **organism** | the organism carrying the contruct if applicable i.e. when plasmids were constructed |
| **DNA delivery** | method of DNA delivery to the organism |
| **DNA** | type of DNA that was to be delivered |
| **flanking region 1** | template and primers used for amplification of the flanking region of the locus that is to be altered; homologous to the site where it is to be integrated |
| **selection cassette** | template and primers used for amplification of the casette used to select for the correctly integradted DNA fragment |
| **flanking region 2 A** | template and primers used for amplification of the flanking region of the locus that is to be deleted; homologous to the site where it is to be integrated<br>or for site directed mutation via primer amplification: template and primers used for the aplification and introduction of the point mutation or introcution of new loci: template and primer amplifiying the loci to be newly integrated |
| **flanking region 2 B** | template and primers used for amplification of the flanking region of the locus that is to be altered with overhangs to flanking region 2A; homologous to the site where it is to be integrated |
| **backbone** | template and primers used to amplify the backbone of the cloning vector |
| **Valdiation Primer Pair 1-3** | primer pairs used in colony PCR |
| **sequencing primer** | primers used when sending Colony PCR (specified for the amplicons) or plasmid preps for Sanger Sequencing |

**Table S4. ddPCR primers and probes used in this study**
Assay name, primer or probe identifier, its sequence, fluorophore, quencher, melting temperature ($T_M$), target locus, and expected amplicon length are indicated.

| assay | Primer/probe number | sequence | Fluoro-phore | Quen-cher | Tm [°C] | locus | Amplicon length [bp] |
|---|---|---|---|---|---|---|---|
| A2 | A2_1 | CAATGCGGCGGCTGCATACG | | | 64 | *nptII* | 103 |
| | A2_2 | ACAAGACCGGCTTCCATCCG | | | 62 | | |
| | A2_s1 | TGCCCATTCGACCACCAAGCG | HEX (qPCR Probe) | BMN-Q535 | 65.3 | | |
| A3 | A3_1 | cagcaggcggaagttgattg | | | 60.5 | *dnaX* | 106 |
| | A3_2 | TGCGTGCTTAGATTGTTGGC | | | 58.4 | | |
| | A3_s1 | CGGTTGAGAAGATGCATGTACTG | 6-FAM | BMN-Q535 | 62.9 | | |
| A4 | A4_1 | TACCGTCGTCGCAAGTTCTG | | | 60.5 | *rpsR* | 124 |
| | A4_2 | CAGTGATACGGCTAGGAACG | | | 60.6 | | |
| | A4_s1 | CGCTTTACAGCTGAGAATGTTGC | Cyanine 5.5 | BMN-Q620 | 62.9 | | |
| A11 | A11_1 | CCATGGCCAACACTTGTCAC | | | 60.5 | *gfp* | 106 |
| | A11_2 | CTTCGGGCATGGCACTCTTG | | | 62.5 | | |
| | A11_S1 | CGGTTATGGTGTTCAATGCTTTGC | Rox | BMN-Q620 | 63.6 | | |

**Table S5. Conjugation frequencies in donor culture setup**
Donor, culturing type, replicate number as well as the cell titer per ml (for liquid cultures) or the total amount (for solid cultures) of plasmid harboring cells are indicated. For solid surface matings the samples were treated like in the evolution experiment but with an incubation time of 24 h. The liquid culture started with equal amounts of the donor and the recipient population and treated as specified for the donor culture in the evolution experiment. Cell counts were evaluated after 24 h.

| Donor | Culturing type | Replicate | Plasmid harboring recipient cell titer/count per ml |
|---|---|---|---|
| BD413 pNCL | liquid | 1 | $2 \times 10^4$ |
| | | 2 | 0 |
| | | 3 | 0 |
| | | 4 | 0 |
| | solid | 1 | $2.35 \times 10^8$ |
| BD413 pCLR | liquid | 1 | 0 |
| | | 2 | 0 |
| | | 3 | 0 |
| | | 4 | 0 |
| | solid | 1 | $2.0 \times 10^6$ |
| BD413 pCHR | liquid | 1 | 0 |
| | | 2 | 0 |
| | | 3 | 0 |
| | | 4 | 0 |
| | solid | 1 | $3.2 \times 10^6$ |

**Table S6. Transformation frequencies of model plasmids at evolution experiment onset**
The transformation frequency (the frequency with which the novel allele has been integrated) at evolution experiment onset is indicated for each model plasmid.

| Model plasmid | Transformation frequency ± 95% CI |
|---|---|
| pNCL | $5.48 \pm 0.60 \times 10^{-3}$ |
| pCLR | $5.54 \pm 1.37 \times 10^{-3}$ |
| pCHR | $2.72 \pm 0.39 \times 10^{-3}$ |
| pCLR-Δ*incC* | $7.96 \pm 2.45 \times 10^{-2}$ |
| pCLR (controlling pCLR-Δ*incC*) | $14.34 \pm 5.87 \times 10^{-2}$ |
| pNCL::$oriV_{pTAD}$ | $7.26 \pm 5.47 \times 10^{-3}$ |
